# Multi-scale Effects of Habitat Loss and the Role of Trait Variation

**DOI:** 10.1101/2023.04.09.536156

**Authors:** Rishabh Bagawade, Koen J. van Benthem, Meike J. Wittmann

**Affiliations:** Department of Theoretical Biology, Faculty of Biology, Bielefeld University, Bielefeld, Germany; Faculty of Science and Engineering, University of Groningen, Groningen, the Netherlands

## Abstract

Habitat loss (HL) is a major cause of species extinctions. Although effects of HL beyond the directly impacted area have been previously observed, they are not very well understood, especially in an eco-evolutionary context. To start filling this gap, we study a two-patch deterministic consumer-resource model, with one of the patches experiencing loss of resources. Our model allows foraging and mating within a patch as well as between patches. We then introduce heritable variation in consumer traits to investigate eco-evolutionary dynamics and compare results with constant or no trait variation scenarios. Our results show that HL indeed reduces consumer densities in the neighboring patch, but when the resources are overexploited, HL in one patch can increase the consumer densities in the neighbouring patch. Yet at the landscape scale, the effect of HL on consumer densities is consistently negative. In presence of HL, patch isolation has positive effects on consumer density in the patch experiencing HL and mostly negative effects on the neighbouring patch. The landscape level pattern depends on which of these effects are dominant at the local scale. Evolution always increased resistance of consumers in the affected patch to HL, with varied effects at the landscape level. Finally, we also show a possibility of landscape level consumer extinction due to HL in a local patch when the cross-patch dependence is high, and foraging and mating preferences are coupled. Eco-evolutionary dynamics can rescue consumers from such extinction in some cases if their death rates are sufficiently small. Our findings show that HL at a local scale can affect the neighbouring patch and the landscape as a whole, and that heritable trait variation can provide some resistance against HL. We thus suggest joint consideration of multiple spatial scales and trait variation when assessing and predicting the impacts of HL.

## 1 Introduction

Habitat loss (HL) is one of the leading causes of biodiversity decline (Fahrig, 2003; McWilliams et al., 2019) and its effects often extend beyond the directly affected habitat. For example, otherwise viable habitat fragments can be affected by the degradation of surrounding matrix or buffer zone, as shown both empirically (Bierregaard et al., 1992; Friesen et al., 1995) and theoretically (e.g. due to increased mortality in the matrix, Cantrell et al., 1998; Cantrell & Cosner, 1999). Effects may also take place at larger spatial scales. For example, in migratory populations, HL at a single wintering site can affect the population densities in the summer breeding sites due to increased competition in the remaining winter habitat by the displaced individuals (Sutherland & Dolman, 1994). Local HL can reduce habitat connectivity, culminating in extinctions at both local and regional scales (Horváth et al., 2019). Furthermore, local HL can affect the stability of the entire community, for example, by constraining the mobility of the remaining community in a smaller region leading to increased encounter rates and thereby increasing the interaction strengths and destabilizing the system (McWilliams et al., 2019). This tendency of HL effects to go beyond the local spatial scale necessitates the study of HL effects over multiple spatial scales.

Besides affecting ecological properties such as interaction strengths and population stability, HL may also affect evolutionary dynamics by altering phenotypic trait variation and the underlying genetic diversity, which in turn may affect ecological processes. Genetic diversity, for example, can be reduced by HL through a reduction in reproductive output and increase in inbreeding (Lowe et al., 2005). Furthermore, HL can influence evolution of species either directly by differential loss of specific types of habitats and associated changes in selection pressures or indirectly by reducing population sizes and thereby also reducing genetic diversity (McClure et al., 2008). A reduction in genetic diversity can even trigger an extinction vortex if the populations become too small (Nabutanyi & Wittmann, 2021, 2022). HL can also influence evolution, for example, of dispersal distance such that removal of entire patches would select for reduced dispersal, but degradation (reducing carrying capacity) of patches would select for longer dispersal in a multi-patch landscape (North et al., 2011).

Conversely, trait variation can play a role in mitigating the population dynamic consequences of HL and environmental change. For example, genetic and phenotypic diversity tend to reduce the vulnerability of populations to environmental change (reviewed in Forsman & Wennersten, 2016), and diversity in individual behavioural traits (risk taking vs avoiding) can promote coexistence in communities experiencing HL (Rohwäder & Jeltsch, 2022). Furthermore, when the trait variation is heritable, it can also help mitigate the effects of environmental change through “evolutionary rescue” (Bell & Gonzalez, 2011; Boeye et al., 2013; Gonzalez et al., 2013), or conversely aggravate the negative effects through “evolutionary trapping” (Ferriere & Legendre, 2013). These reciprocal effects can lead to eco-evolutionary feedbacks, between the population dynamic consequences of HL and trait variation, which are increasingly being recognised (Legrand et al., 2017; Faillace et al., 2021; Gawecka et al., 2022).

Apart from feedbacks between HL effects and trait variation, multi-scale spatial processes can also interact with trait variation and evolution, thereby influencing population dynamics. For example in a two-patch model, multi-scale density dependence, where density in one patch influences the dynamics in the adjacent patch and vice versa, can lead to adaptation to emerging low and high density patches (van Benthem & Wittmann, 2020). Moderate immigration from a nearby source patch can provide the necessary genetic material for local adaption and eventually lead to evolutionary rescue in a sink habitat (Gomulkiewicz et al., 1999). Spatial processes such as range expansion can lead to bursts of evolutionary change under genetic drift, or even directional evolution under spatially structured selection gradients (Polly, 2019). Furthermore, spatially structured intraspecific trait variation (ITV) can promote species coexistence when species’ responses to habitat conditions are different (Banitz, 2019).

In summary, HL affects populations over multiple spatial scales, and these effects can be altered by trait variation. The interplay between HL and multi-scale interactions can lead to eco-evolutionary feedbacks. Currently, such eco-evolutionary processes for interacting populations exposed to HL are not well understood. Knowing that mathematical models can act as a proof of concept to address such complex phenomena (Servedio et al., 2014), we take a step in that direction by modeling a deterministic two patch consumer-resource system with two spatial scales: local (patch-level) and landscape (both patches combined). We use the term “spatial scale” in a general sense, and not in the specific context of sampling (as used by Wiens, 1989). In our model, consumers can mate and forage within and between patches, with one of the patches experiencing HL. We explore how HL effects propagate from local to landscape scale, how effects of HL change due to the presence of a nearby unharmed patch, and whether heritable trait-variation in the consumer traits helps mitigate these effects through eco-evolutionary dynamics.

## 2 Methods

We use a system of coupled ordinary differential equations (ODEs) to describe a two-patch model where each patch contains a consumer population and a resource population (Fig. 1a). We define *R*_1_ and *R*_2_ as the resource densities in patch 1 and patch 2, respectively. *N*_1_ and *N*_2_ are the densities of consumers whose birthplace is patch 1 and patch 2 respectively. Temporary movement between patches does not change *N*_1_ and *N*_2_. Such movement occurs in the event of cross-patch foraging, with individuals foraging for resources in the other patch, and cross-patch mating, with individuals mating with a partner from the other patch. Resources, by contrast, do not move between patches. We model habitat loss (HL) as the death or removal of the resources in patch 2 because of external anthropogenic factors such as harvesting, contamination, or degradation of resources. Therefore, consumers are the species of interest whose fate under HL is investigated in this study. We further assume that HL does not cut off the access of consumers to that patch, i.e., consumer breeding sites can still exist in that patch. Another important assumption is that the cross-foraging and mating preferences do not change after HL, unless allowed to evolve under the eco-evolutionary model. Below we define a purely ecological model, and then extend this model by including eco-evolutionary dynamics.

**Figure 1:**
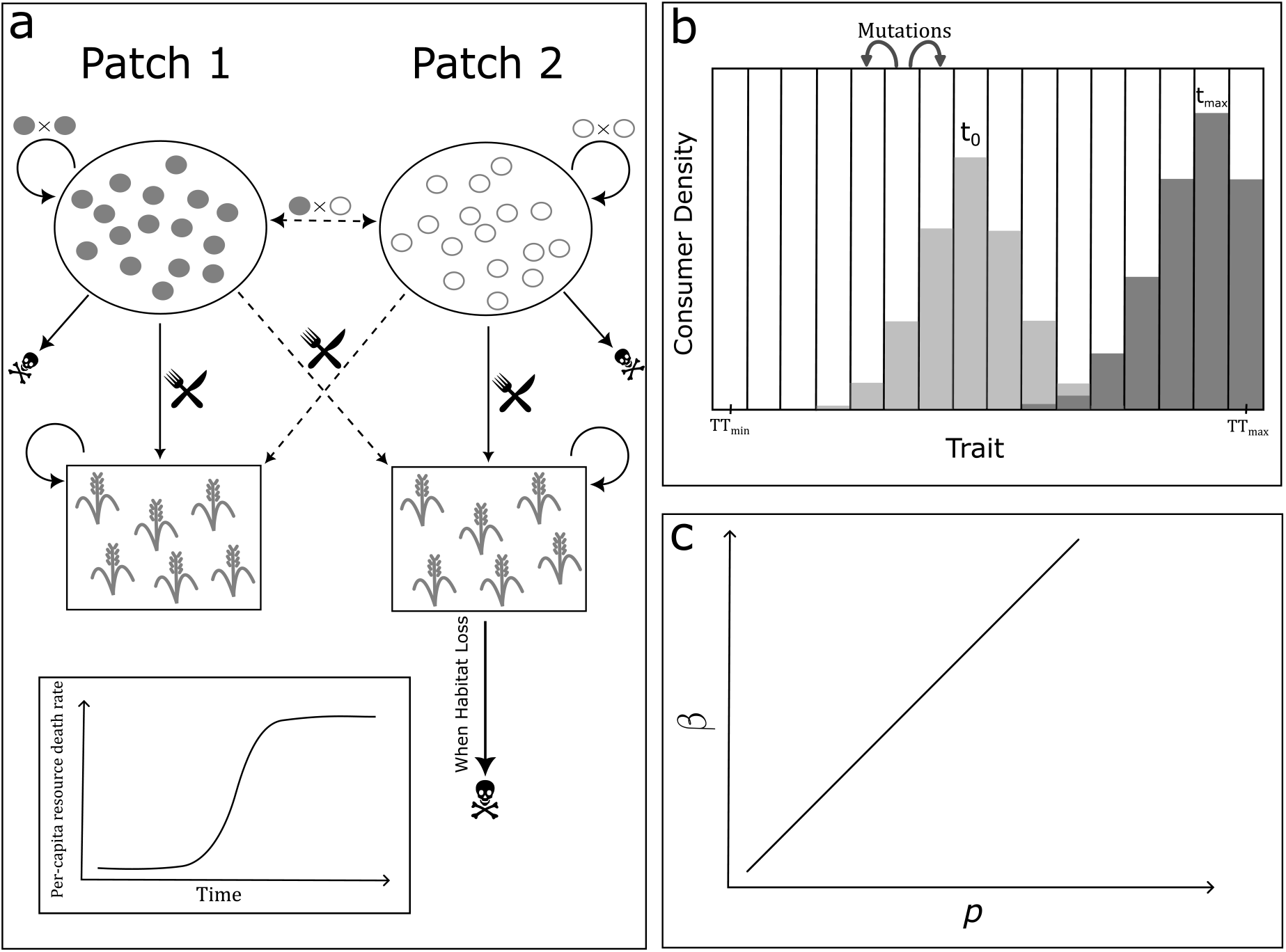
Schematic overview of the model. a. Schematics of the two patch system with consumers and resources. Dotted arrows indicate cross-patch interactions (foraging or mating). In the presence of habitat loss, resources in patch 2 are depleted through a logistically increasing resource death or removal rate (inset). b. In the presence of trait variation, populations are divided into bins where their growth rate depends on the trait value assigned to that bin. Example trait distributions are shown for initial (at *t*_0_, light grey) and final (at *t*_max_, dark grey) time points. The trait distribution is kept constant in the case of constant variation, and it evolves in the case of heritable variation. Note that the no-variation scenario is equivalent to there being only one bin with the trait value equal to the initial mean trait value of the other two variation scenarios. c. Linear relationship when traits *p* and *β* are coupled such that *p* = *β*.

### Ecological model

The resources are replenished according to a logistic growth model:

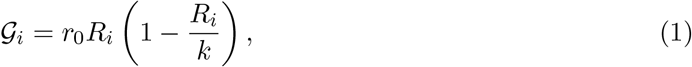

where 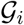 is the population growth rate for the resource in patch *i* ∈ {1, 2}, *r*_0_ is the maximum per-capita growth rate, and *k* is the carrying capacity. All parameters with their default values are noted in table 1.

**Table 1:** Parameters, their description, intervals of possible values and default values

| Parameter | Description | Interval | Default Value |
| --- | --- | --- | --- |
| $a_0$ | per-capita death rate | $[0, \infty]$ | 0.1 |
| $b_0$ | maximum per-capita resource consumption rate | $[0, \infty)$ | 30 |
| $b_1$ | half-saturation constant for resource consumption | $[0, \infty)$ | 500 |
| $p$ | within-patch foraging preference | $[0, 1]$ | 0.7 |
| $\beta$ | within-patch mating preference | $[0, 1]$ | 0.8 |
| $\theta$ | Allee effect parameter | $[0, \infty)$ | 2 |
| $\epsilon$ | per-capita resource conversion efficiency | $[0, \infty)$ | 0.2 |
| $\epsilon_c$ | cross-patch foraging efficiency relative to within-patch | $[0, 1]$ | 0.9 |
| $r_0$ | maximum growth rate of resources | $[0, \infty)$ | 0.5 |
| $k$ | carrying capacity of resources | $[0, \infty)$ | 50 |
| $D$ | maximum rate of resource degradation or removal | $[0, \infty)$ | 0.5 |
| $s$ | steepness of logistic resource removal | $[0, \infty)$ | 0.003 |
| $t_{1/2}$ | time when the resource removal rate is $D/2$ | $[0, \infty)$ | 3500 |
| $N_{0,1}, N_{0,2}$ | initial consumer densities for patch 1, patch 2 | $[0, \infty)$ | 3, 4 |
| $R_{0,1}, R_{0,2}$ | initial resource densities for patch 1, patch 2 | $[0, \infty)$ | 10, 10 |
| $\mu$ | mutation rate | $[0, 1]$ | 0.3 |
| $B$ | number of bins when trait variation is present | $\mathbb{N}$ | 21 |

Assuming a type II functional response, the per-capita resource consumption rate by consumers in patch *i* of the resource in patch *j*, 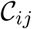, is

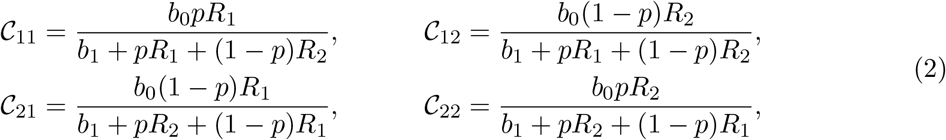

where *b*_0_ is the maximum per-capita resource consumption rate, *b*_1_ is the half-saturation constant, *p* is the within-patch foraging preference, and consequently (1 – *p*) is the cross-patch foraging preference. Note that *pR*_1_ + (1 – *p*)*R*_2_ corresponds to the average resource density weighted by the amount of within-patch and cross-patch foraging for consumers in patch 1 (analogously, *pR*_2_ + (1 – *p*)*R*_1_ for patch 2 consumers).

As mentioned earlier we model HL via resource loss in patch 2 at the rate

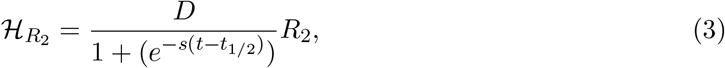

where *D* is the maximum rate of resource removal, *t*_1/2_ is the time at which the resource removal rate is *D*/2, and *s* controls the steepness (i.e. how sudden the resource removal rate changes around *t*_1/2_). Thus, resource removal starts slowly, then becomes increasingly fast, before finally approaching a maximum (Fig. 1a, inset). See SI S1 for further discussion about the shape of HL function 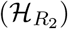. For the scenarios without HL, *D* = 0 i.e. 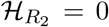. Taking everything together, the ecological dynamics of resources in the two patches are given by:

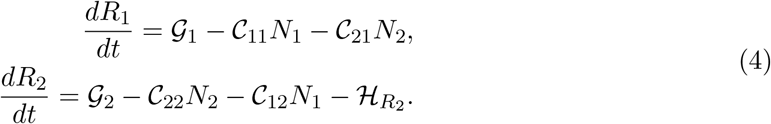

Consumer population growth rates in patch 1 and 2 are modeled as:

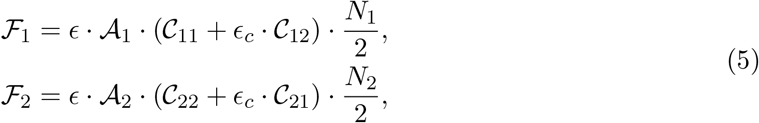

where *ϵ* is the efficiency of resource conversion, *ϵ_c_* ∈ [0, 1] is the cross-patch foraging efficiency relative to the within-patch foraging efficiency, 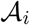 is a mate finding Allee effect term (see below) in patch *i*, 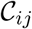 are the per-capita consumption rates as defined in equation (2). Assuming a constant 1:1 sex ratio, the population densities (*N_i_*) are divided by a factor of 2 to model that only half of the population (females) gives birth.

When HL leads to reduced population sizes, populations may experience an Allee effect (Swift & Hannon, 2010), i.e. a decrease in survival and/or reproduction with decreasing population size in small populations, for example because it becomes more difficult to find mating partners (Fauvergue, 2013). We thus model the probability of finding at least one male mating partner if there are *M* available males as (1 – *e^-θM^*) (Dennis, 1989; Fauvergue, 2013). Here, *θ* is inversely proportional to the strength of the Allee effect. To determine the value of *M*, we have to account for cross-patch mating. We define the within-patch mating preference as *β* ∈ [0, 1], and consequently (1 – *β*) equals the cross-patch mating preference. The number of males that a female sees is determined as the average of males in her own patch and in the other patch, weighted by *β* and (1 – *β*), respectively. The probability of at least one successful mating event for a female in patch *i*, 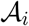, thus becomes:

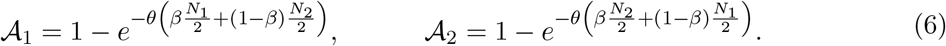

We define *q_ij_* as the fraction of newborns produced by females in patch *i* via mating with males in patch *j* as:

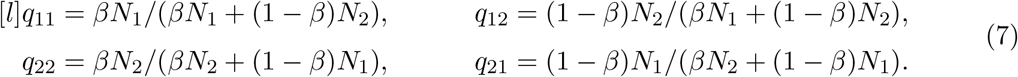

The females that mate within-patch are assumed to produce their offspring in their own patch, whereas females that engage in cross-patch mating are assumed to be equally likely to produce their offspring in either of the patches. This means, for patch 1, offspring coming from withinpatch mating 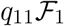 stays in the patch 1, whereas, half of the offspring from cross-patch mating (i.e. 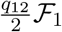) stays in patch 1, and the other half goes to patch 2. Dynamics for patch 2 happen analogously. Lastly, consumers die with a constant per-capita death rate *a*_0_. Therefore, the consumer ecological dynamics are:

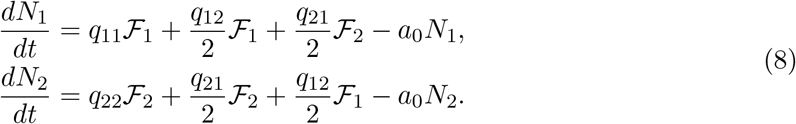

The resource dynamics in eq. (4) and consumer dynamics in eq. (8) together form the full ecological model.

### Eco-evolutionary model

We now add heritable trait variation and mutations to the consumer population such that the consumer parameters are considered as traits (similar to van Benthem & Wittmann, 2020). We investigated intraspecific variation in per-capita resource conversion efficiency *ϵ*, maximum percapita resource consumption rate bo, or half-saturation constant for resource consumption *b*_1_, with only one parameter varying at a time. Moreover, we looked at a scenario with two coupled traits (Fig. 1c): within-patch foraging preference *p* and within-patch mating preference *β* such that *p* = *β*. This choice makes biological sense, because spending more time foraging in one patch might also lead to a higher chance of finding a mate in the same patch. To better understand how ecological and evolutionary consequences of trait variation affect the population dynamics, we also implement runs with constant trait distribution. Despite lacking evolution, this scenario may still deviate from the no variation case through non-linear averaging (Bjørnstad & Hansen, 1994; Ruel & Ayres, 1999).

We assume traits to be determined by an infinite number of loci with alleles of small effect, which would give rise to a continuous trait distribution. We approximate this trait distribution by discretizing it into *B* bins, with a population density of *n_i,b_* and trait value *z_b_* in bin *b* in patch *i* (Fig. 1b). The total consumer density in patch *i* is thus 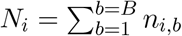 Population dynamics for each bin are determined by the population densities in the patches, the resource densities and the bin-specific trait value *Z_b_*. The first and last bin correspond to the minimum and maximum trait values, and are thus defined as *z*_1_ = *TT_min_* and *z_B_* = *TT_max_*, with equally spaced trait values for the intermediate bins. The initial trait distribution is obtained by truncating and re-normalizing a normal distribution with mean *TT_mean_* and standard deviation *TT_sd_*. That is, to ensure that the initial distribution stays within the defined interval, we set those bins that are beyond *TT_min_* or *TT_max_* to zero, and, to maintain the mean and keep the distribution symmetric, we also set their counterparts on the other side of *TT_mean_* to zero. In the eco-evolutionary model, the trait distribution evolves over time through changing number of individuals in the different bins. In the scenarios where *p* and *β* are coupled (*p* = *β*), each bin now represents two traits of identical value.

Fecundity is determined by the female’s trait value. When a female mates, she randomly picks a mating partner in proportion to bin densities, i.e. there is no sexual selection and males can mate with an arbitrary number of females. Offspring trait values are determined by the average trait value of the two parents. If the mid-parent trait value is on the boundary between two adjacent bins, the offspring density is evenly split between these two bins. The offspring trait distribution of any given female therefore depends on the paternal trait distributions in both patches, and on the chances that she mates with males in each of the two patches. In the scenarios with heritable *β*, the population can evolve for females to have a preference for mating with males in either the own patch or in the other patch.

Putting everything together (see SI S2 for the technical details), we then obtain *f_i,b_*, the rate at which offspring are added to bin *b* in patch *i* due to reproduction in the whole population. Then, in the absence of mutations, our consumer population would develop as follows:

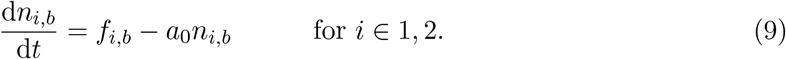

The trait distribution of the offspring is then further affected by mutations. These are modelled deterministically such that a fraction *μ* (the mutation rate) of the offspring density in bin *b* mutates away from the bin and is evenly distributed between bins *b* + 1 and *b* – 1. For the first and the last bin, mutation occurs only in one direction with half the rate (*μ*/2). Thus the population dynamics for the bins in patch *i* are:

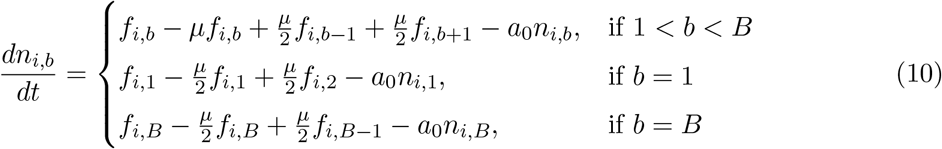

where *i* ∈ {1, 2}.

The resource dynamics for the two patches, in presence of trait variation in consumers, are given by:

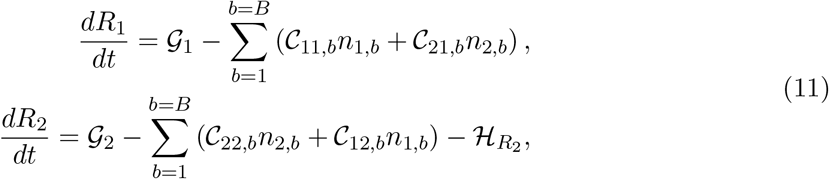

where the consumption terms 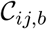 depend on the bin *b* whenever consumer trait values affect the consumption rates.

As a baseline scenario to the eco-evolutionary case, we also model a constant variation case where the densities in the trait bins are redistributed after every time step such that the trait distribution over time always remains identical to the initial distribution of the eco-evolutionary model. These models thus disregard both mutations and inheritance. We further set the trait value in the ecological (no variation) scenario to be identical to the mean trait from the initial distribution in the other two scenarios.

The system of equations is solved numerically using the `deSolve` (Soetaert et al., 2010) package in R (R Core Team, 2022). The numerical solver is run until time 8000 for all the results, unless stated otherwise. We made sure this was long enough for the population densities to stabilize, as confirmed by visual inspection of all underlying time series. Since *a*_0_ = 0.1 (in the default settings) is defined as the per-capita consumer death rate per unit time, 1/*a*_0_ = 10 roughly correspond to the life-span of the consumers such that the simulation time corresponds roughly to 800 generations.

We focus on the parameter region where the populations reached stable (non fluctuating) densities by the end of the runs (but see Fig. S5 for an example with fluctuating dynamics). Density at the last time point is thus considered as the equilibrium density. We are aware that alternative stable states may exist for our system, which is why we compare scenarios with consistent initial conditions. We do not calculate all the fixed points and stability of the full ecological model, however we do provide analytical treatment for a simplified approximation of the full ecological model in SI S3.

### Scenarios explored

The scenarios explored in this study are based on presence or absence of three aspects: habitat loss (HL), trait variation, and patch isolation. First, we compare HL and no HL scenarios. Second, we compare no trait variation (ecological model), and heritable variation (eco-evolutionary model) along with constant variation as the baseline for the eco-evolutionary model. Lastly, we compare the effects of HL when the patches are connected (this is the default scenario) to when they are isolated. Patch isolation is modeled by assuming that the two patches have no access to each other (because of distance or physical barriers) i.e. *N*_2_, *R*_2_ = 0 for patch 1 dynamics and vice versa for patch 2. Here we assume that the consumers still go out of the patch with foraging preference 1 – *p* but cannot access the resources nor potential mating partners in the other patch.

Using the models and scenarios described above, we investigate the effects of local HL over multiple spatial scales and how eco-evolutionary dynamics influence these effects. We specifically address the following questions: 1. How does HL in one of the patches affect the consumer densities in that patch, the neighbouring patch, and the landscape as a whole? 2. How does the presence of a neighbouring patch affect the dynamics in the patch experiencing HL? 3. Can eco-evolutionary dynamics mitigate the multi-scale effects of HL?

## 3 Results

We start with the ecological model, for which we compare the dynamics with and without habitat loss (HL) (Fig. 2). After sufficient time, the consumer and resource densities equilibrate. When HL is absent (blue lines), densities in patch 1 and 2 are equal at equilibrium. In the presence of HL, the resources in patch 2 start to decline and disappear completely around time 4000 (Fig. 2b,f, red dotted lines), with the consumers in patch 2 already going extinct well before time 4000 (Fig. 2a,e, red dotted lines). As a consequence, between time 2000 and 4000, there is a transient increase in resource density in patch 2 and consequently in consumer density in patch 1 because there is no more competition from patch 2 consumers.

**Figure 2:**
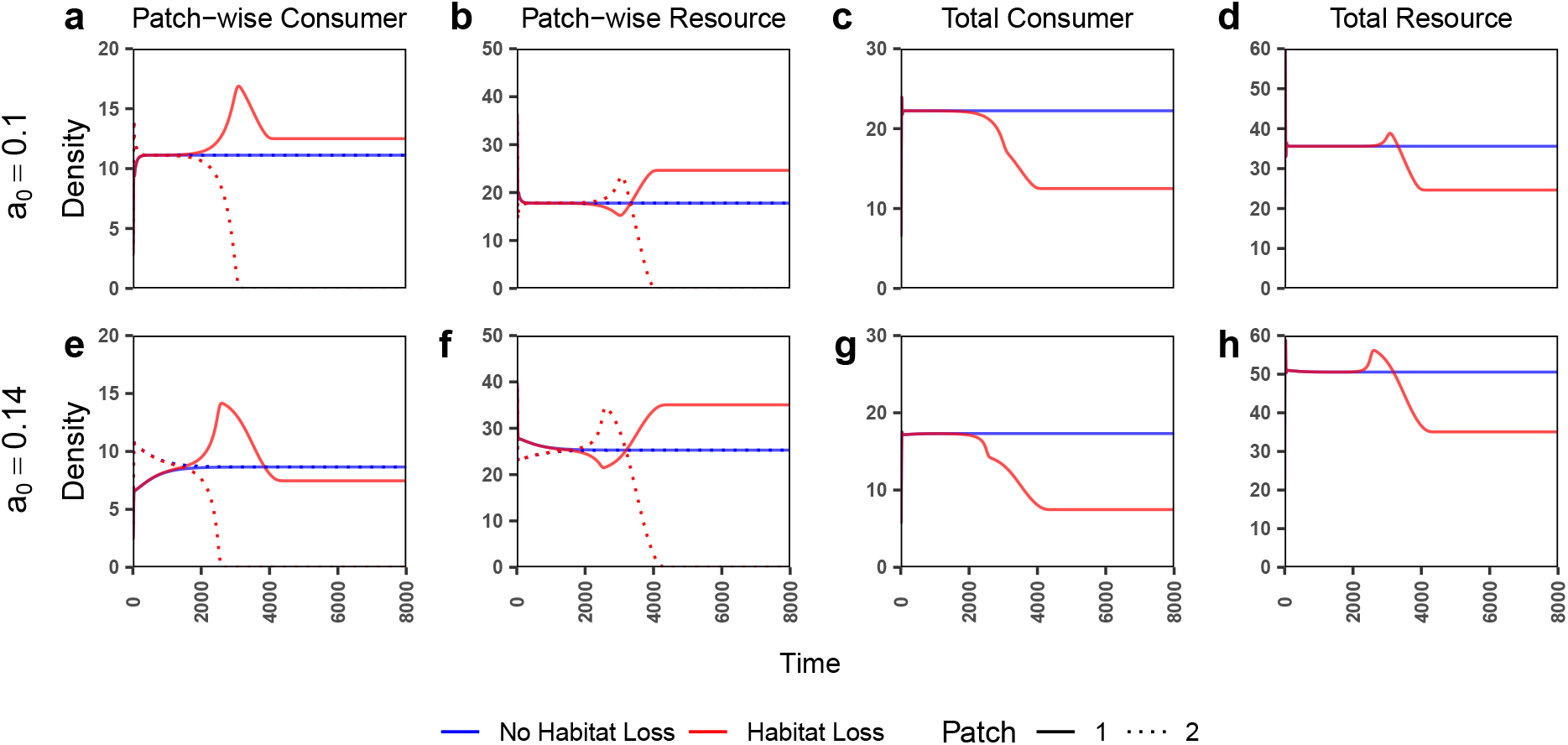
Time-series plots under the ecological model (no trait variation) for patch-wise, and total (landscape scale) consumer (*N*_1_ + *N*_2_) and resource (*R*_1_ + *R*_2_) densities in presence or absence of habitat loss. Note that for panels c, d, g, and h solid lines imply total densities (i.e. there is no dotted line). All parameter values are default values from table 1, except for the *a*_0_ values which are mentioned for each row.

Resource degradation in patch 2 may lead to higher patch 1 equilibrium consumer density when the consumer death rate *a*_0_ is low (Fig. 2a, observe red solid line above the blue line at equilibrium), but not when *a*_0_ is high (Fig. 2e). This is counterintuitive, because when the resources in patch 2 are lost, patch 1 consumers can no longer cross-forage and thereby the effective per-capita resource consumption must reduce irrespective of *a*_0_ (SI S4). Abrams (2002) has shown that when resources are overexploited in a consumer-resource system, reduction in per-capita consumption can increase the consumer density. Our simplified model does indeed work analogous to the work by Abrams (See SI S4). Thus, we infer that the system is in the overexploitation regime at low *a*_0_ (0.1), where the HL-induced reduction in effective consumption rate leads to an increase in consumer density (Fig. 2a). At higher a0, the system has no overexploitation, and hence HL leads to lower consumer densities in patch 1 (Fig. 2e). Along with *a*_0_, other parameters related to consumer efficiency such as *b*_0_, *ϵ*, and *b*_1_ can also show such overexploitation-based increase in patch 1 consumer density after HL in patch 2 (See SI S4 and S5). Importantly, irrespective of the observations for individual patches, the total consumer (and resource) density at the landscape levels is always lower after HL (Fig. 2c,d,g,h).

Including trait variation in the ecological model does not necessarily lead to qualitative changes in population dynamics (Fig. 3). For example, when variation is present in resource conversion efficiency *ϵ* or maximum resource consumption rate *b*_0_, the constant variation (dashed lines) and no variation (solid lines) scenarios are indistinguishable (Fig. 3). Both *ϵ* and *b*_0_ affect the growth rate linearly, and the mean trait value for the two scenarios is always identical. Therefore the effect of constant variation is not expected to be different from no variation (Ruel & Ayres, 1999). However, the dynamics of the no variation and constant variation case differ (even with identical mean trait value) if there is non-linear dependence between growth rate and the trait, for example when trait variation is present in *b*_1_ (Fig. S8). This can happen through non-linear averaging (Bjørnstad & Hansen, 1994; Ruel & Ayres, 1999).

**Figure 3:**
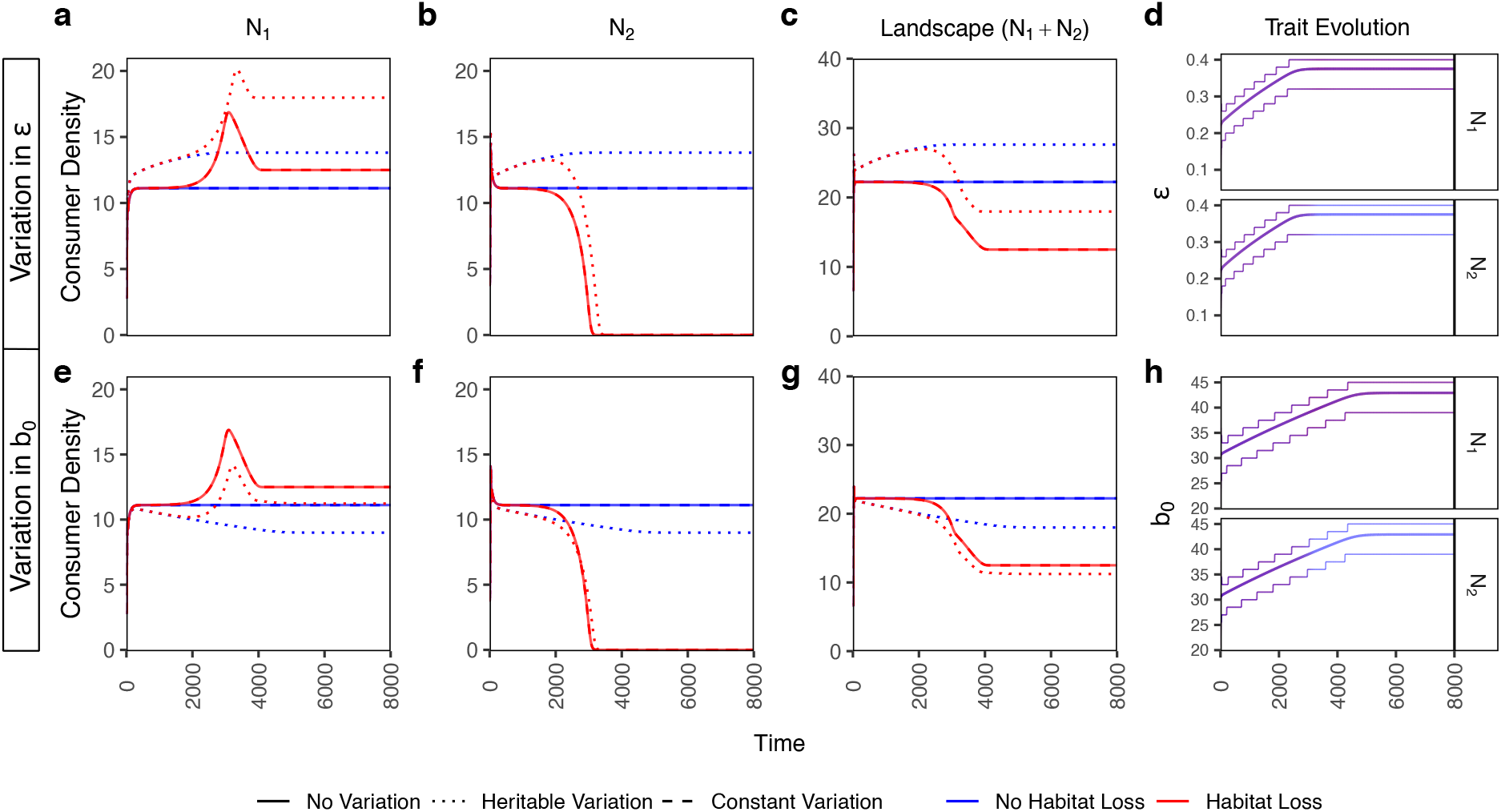
Time series plots for patch and landscape level consumer densities for the three variation scenarios in the presence and absence of habitat loss. The evolving trait for the first row is resource conversion efficiency *ϵ* with initial mean *TT_mean_* = 0.2, and standard deviation *TT_sd_* = 0.08 with maximum possible trait value *TT_max_* = 0.4, and minimum *TT_min_* = 0. The evolving trait for the second row is maximum resource consumption rate *b*_0_ with *TT_mean_* = 30, *TT_sd_* = 5, *TT_max_* = 45, and *TT_min_* = 15. The last column shows the trait evolution where the central line denotes the mean trait value and the thinner outer lines denote the range which contains 90% of the density. Line transparency in the trait evolution plots denote consumer density. Note that the red and blue lines are overlapping in the trait evolution plots. All other parameter values are at their default values from table 1.

However, even with a linear dependency, when the trait variation is heritable (eco-evolutionary scenario), it can lead to either higher or lower equilibrium densities than the other two variation cases depending upon which trait is allowed to evolve. For example, even though both *ϵ* and *b*_0_ evolve to higher trait values, when *b*_0_ evolves the equilibrium density is lower in the eco-evolutionary model (Fig. 3, bottom row, dotted vs. solid line), while it is higher when *ϵ* evolves (Fig. 3, top row). The different behaviour arises because *b*_0_ influences the per-capita consumption rate whereas *ϵ* does not. Thus, an increase in *b*_0_ leads to more overexploitation and thereby decreases consumer density, whereas an increase in *ϵ* just benefits the consumers and thereby increases consumer density (also see Fig. S4b,c).

We further explore the mutual effect that the patches have on each other, by running scenarios in which we artificially set consumer and resource density in the other patch to zero when determining foraging and mating for either patch (Fig. 4). This setting represents isolated patches. We specifically focus on the effect of the resource removal rate *D* under all three variation scenarios. In the absence of HL (*D* = 0), patch isolation has a positive effect on both consumer and resource density in the individual patches as well as on the landscape level (orange lines above turquoise lines at *D* = 0 in Fig. 4). This again is a consequence of overexploitation at lower consumer death rate (*a*_0_ = 0.1 in Fig. 4). In the isolated case, consumers still spend a fraction 1 – *p* of effort searching for resources outside their patch which reduces the resource consumption pressure compared to the connected patches. As expected, the effect disappears at higher consumer death rates when the system is not in the overexploitation regime (*a*_0_ = 0.14 in Fig. S9).

**Figure 4:**
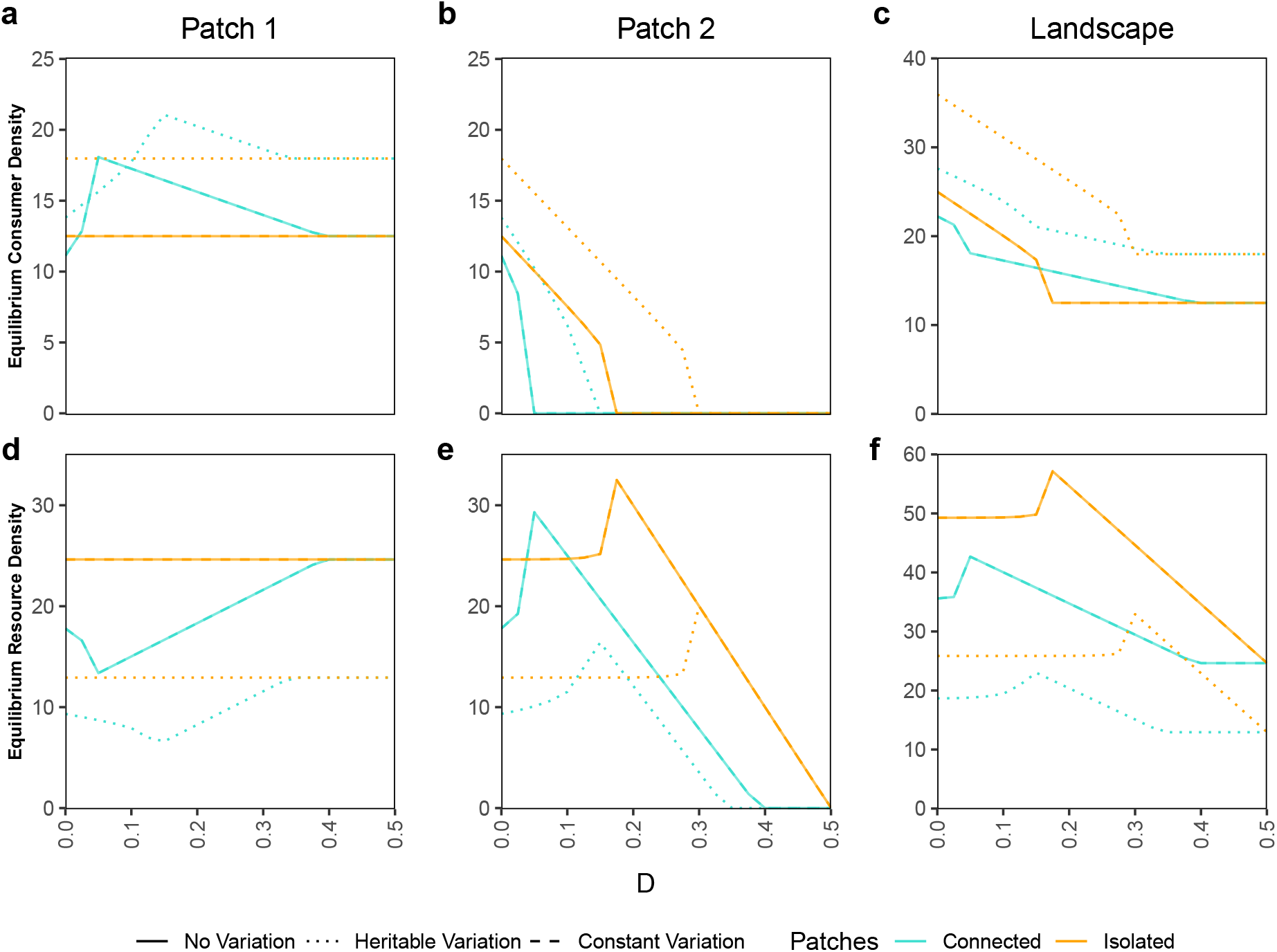
Individual patch and landscape scale equilibrium consumer (row 1) and resource (row 2) densities under HL scenario with heritable variation, constant variation, or no variation. The three trait variation scenarios when the patches are either connected (neighbouring patch is accessible) or isolated (neighbouring patch is not accessible) are compared over a range of maximum resource degradation rate *D* values. The varying (or evolving) trait is the resource conversion efficiency *ϵ* with initial mean *TT_mean_* = 0.2, and standard deviation *TT_sd_* = 0.08 with maximum possible trait value *TT_max_* = 0.4, and minimum *TT_min_* = 0. No variation and constant variation lines are overlapping. Recall that resources do not have trait variation. Resource densities shown here are for the corresponding consumer trait variation scenarios. Note that the equilibrium densities are calculated for 21 equidistant points in each parameter range but they are depicted by lines for better clarity. All other non-varying parameter values are at their default values from table 1.

In the presence of HL (*D* > 0), the pattern changes: Here, patch 1 consumers generally have higher equilibrium densities in the connected case compared to the isolated case. This happens likely because in patch 2 the consumers go extinct at lower *D* value than the resources (Fig. 4b,e), which provides cross-foraging opportunities to patch 1 consumers in connected patches as long as there are some resources existing in patch 2 (*D* < 0.4; Fig. 4a,e). For patch 2, which directly experiences HL, foraging competition by patch 1 consumers in connected patches causes extinction of patch 2 consumers at even lower values of *D* as compared to patch isolation (Fig. 4b, turquoise and orange lines; also see Fig. S10b,e for corresponding results without Allee effect). As a result of these observations in the individual patches, at the landscape scale, patch isolation gives higher consumer density at lower values of *D*, but as the value of *D* increases, patch connectedness gives a higher consumer density (barely visible in the heritable variation case), and finally the difference between the two disappears at even higher *D* values (Fig. 4c, turquoise and orange lines).

With *ϵ* as the evolving trait, patch 2 consumers with heritable variation can withstand higher *D* values compared to the other two variation scenarios (Fig. 4b, compare dotted lines with solid and dashed lines). Consumers with heritable variation also generally have higher equilibrium densities everywhere (Fig. 4, also see Fig. S7), except for a small range of *D* values for patch 1 consumers when the patches are connected (Fig. 4a). Here heritable variation prevents consumer extinction in patch 2, thereby maintaining higher competition for patch 1. At the landscape scale, this effect is averaged out and we observe consistently higher consumer density in the presence of heritable trait variation (Fig. 4c, dotted lines above solid lines). Interestingly, with trait variation in *b*_1_ (Fig. S11), heritable variation leads to considerably lower equilibrium consumer densities at the landscape level, while still helping patch 2 consumers to sustain higher *D* values (Fig. S11b, dotted lines).

Independent evolution of *p*, without corresponding evolution in *β* may be unrealistic, because higher within-patch foraging likely also implies higher within-patch mating. We therefore explore the case where *p* and *β* are coupled, specifically *p* = *β* (Fig. 5). Here, we have increased patch inter-dependency by setting the initial within-patch foraging (and mating) preference lower (*p* = *β* = 0.65) than the default scenario. At lower death rates (*a*_0_ = 0.1) and without HL, the consumers in the two patches can coexist, while at higher death rates (*a*_0_ = 0.15 or 0.18) the patch with higher initial density (here patch 1) outcompetes the other patch (Fig. 5, blue lines; also see Fig. S12 where patch 1 starts with lower initial density and gets out-competed instead). Furthermore, without HL, the trait (*p* = *β*) evolves towards higher values when consumers in the two patches are coexisting (Fig. 5d, blue lines), and towards lower trait values when they are not coexisting (Fig. 5h,l, blue lines).

**Figure 5:**
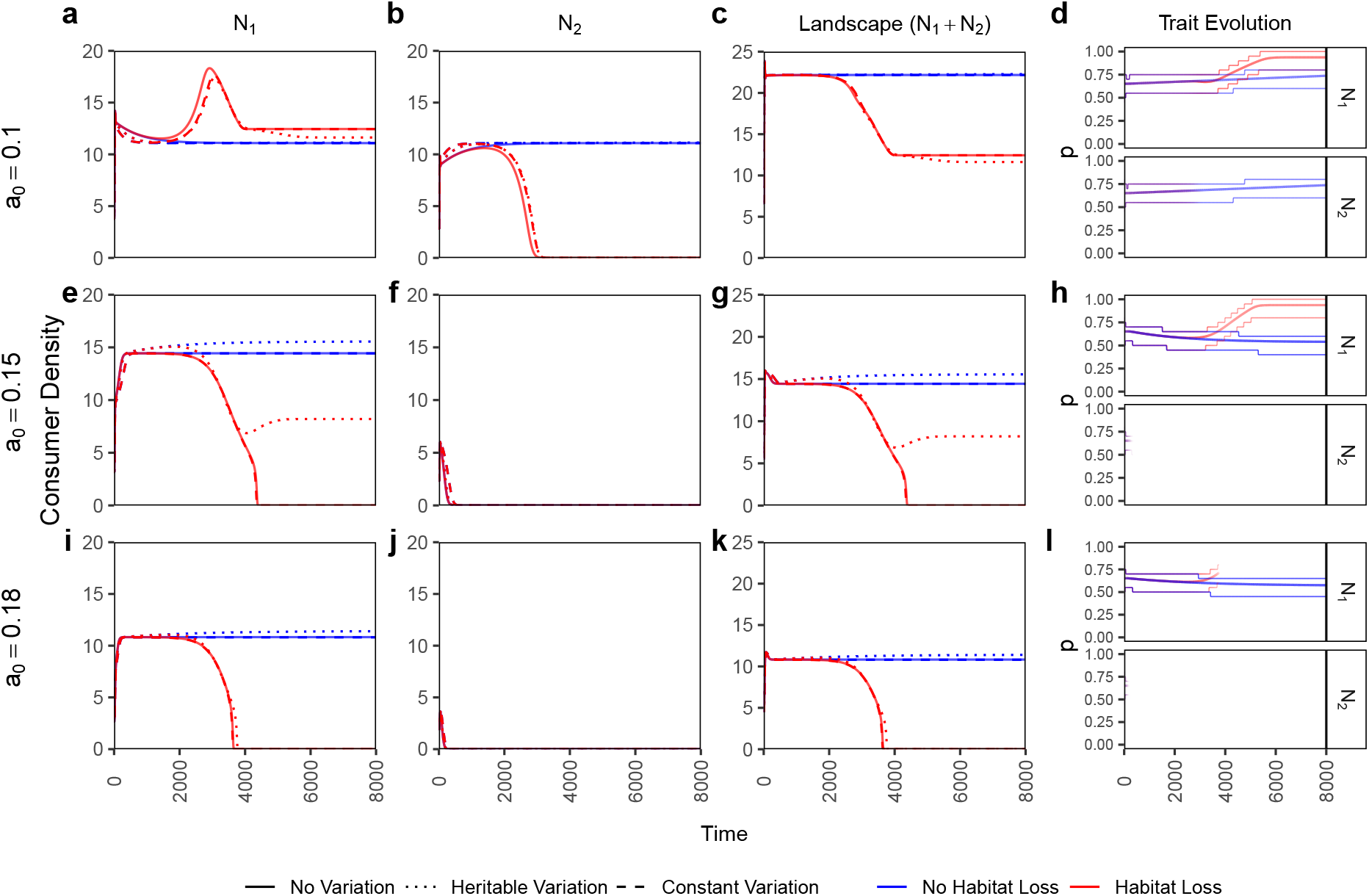
Time series plots of a scenario with coupled within-patch foraging preference *p* and within-patch mating preference *β* such that *p* = *β*. Patch and landscape level consumer densities are shown for the three variation scenarios in the presence and absence of habitat loss. The last column shows the trait evolution of *p* = *β* where the central line denotes the mean trait value and the thinner outer lines denote the range which contains 90% of the density. Line transparency in the trait evolution plots denote consumer density. *p* and *β* both are evolving together with initial mean *TT_mean_* = 0.65, and standard deviation *TT_sd_* = 0.1 with maximum possible trait value *TT_max_* = 1, and minimum *TT_min_* = 0. Initial consumer densities are switched from the default i.e. *N*_0,1_ = 4 and *N*_0,2_ = 3. All other parameter values are default values from table 1 except the *a*_0_ values that are denoted for each row.

In the HL scenario, patch 2 consumers go extinct (Fig. 5 second column, red lines) and then the patch 1 consumers quickly evolve towards higher trait (*p* = *β*) values (Fig. 5 fourth column). Patch 1 consumers survive at the lowest death rate (Fig. 5a, red lines), and when *a*_0_ = 0.18 patch 1 always goes extinct (Fig. 5i, red lines). At this high death rate patch 1 consumers rely on resources in patch 2 for survival; once these get degraded (HL case) the patch 1 consumers also suffer extinction. An interesting case occurs at intermediate death rates, where the extinction process is slow enough such that evolutionary rescue occurs in patch 1 for the heritable variation case only (*a*_0_ = 0.15; Fig. 5e red dotted line). These dynamics at the patch scale translate to the landscape scale such that, under the given parameter conditions, the consumer density in the whole landscape can go extinct after HL in a local patch (Fig. 5g,k), even long after the consumers in the directly affected patch have gone extinct (Fig. 5f,j).

## 4 Discussion

In this study, we investigated the effects of habitat loss (HL) at the local (patch) and landscape scale in the presence or absence of trait variation using a two-patch consumer-resource system. Our results show that: 1. HL has detrimental effects for consumers in the directly affected patch and on the landscape as a whole, but it can sometimes positively affect the consumer density in the adjacent patch (Fig. 2). 2. Patch connectedness can be detrimental to consumers in the patch experiencing HL and beneficial to the adjacent patch. Whether patch connectedness leads to higher consumer density on the landscape scale depends on the intensity of HL, and the existence of overexploitation and/or evolution (Fig. 4; Fig. S9). 3. Evolution allows a patch directly affected by HL to sustain higher intensities of HL, irrespective of whether it increases or decreases the consumer density in the adjacent patch or the landscape (Fig. 4; Fig. S11). 4. Lastly, we also show a scenario where HL in a single patch can lead to consumer extinction at the landscape scale, which could be avoided (at low and intermediate consumer death rate) via evolutionary rescue (Fig. 5). These findings highlight the importance of joint consideration of trait variation and multiple spatial scales when predicting the effects of HL, the prerequisite for management decisions.

### Multi-scale context

The importance of choosing an appropriate spatial scale for ecological measurements has been extensively discussed in the literature (Wiens, 1989; Levin, 1992; Holland & Yang, 2016). Even HL and fragmentation have been found to have a scale-dependent relationship with ecological response variables such as population status (sustained or declining) (Fuhlendorf et al., 2002), and habitat suitability (Treglia et al., 2018). Conservation and management is also benefited by simultaneous consideration of local and landscape scales (Garden et al., 2010; Bergman et al., 2012). However, the majority of these studies are related to differences in statistical measurements due to considering more or less area under the domain of analysis. Our results, on the other hand, not only show how effects of HL can differ at the patch and landscape scale, but also shed a light on how effects of HL can propagate from one patch to another, thus providing one possible explanation of why measurements can be sensitive to scale.

### Cross-patch effects

HL in patch 2 can either positively or negatively affect the consumer densities in the neighbouring patch (Fig. 2). The negative effect can be attributed to a loss of cross-foraging options. Potential positive effects occur because of reduced overexploitation through a reduction in the per-capita consumption rate in patch 1 after HL (see SI S4 for details). Overexploitation happens when resources have logistic growth, and the consumers are highly efficient so that they end up pushing the resource density below its optimum growth rate (i.e. below half the carrying capacity *R*\* < *k*/2, see SI S4). In such cases, paradoxically, an apparent increase in consumer density may actually signal a deterioration of the system (Abrams, 2002). Overexploitation is a known property of consumer resource systems and has been shown in a model with two consumer species and two resource types before (Abrams, 2002), as well as in other modelling studies (Prakash & de Roos, 2002; Abrams, 2009, 2019). Unlike Abrams (2002), our study has mating, eco-evolutionary dynamics, HL based on resource loss (not consumer parameters), and a spatial interpretation. Importantly, since many prey populations are below half of their carrying capacity (Shurin et al., 2002; Abrams, 2019), overexploitation is expected to occur frequently in nature assuming prevalence of logistic-like density dependence in the populations (Abrams, 2019). Positive effects in the remaining patches have also been observed through other mechanisms such as edge effects (Ries et al., 2004) or in metapopulations through interaction of competitive ability and colonization-extinction rates (Nee & May, 1992).

The reverse effect of the remaining patch on the patch experiencing HL (patch 2) is mostly detrimental, through cross-patch competition. This effect pushes patch 2 consumers to extinction at even lower intensity of HL than without such competition (Fig. 4). The possibility of negative effects of such cross-patch competition has been researched using patch isolation experiments for example in a coral - coral associated fish system (Bonin et al., 2011), and a leaf litter - decomposing bacterial communities system (Spiesman et al., 2018). However, our results also show a possible rescue of patch 2 consumers in the HL scenario by the presence of a neighbouring patch when cross-mating is common (*β* < 0.5), while cross-foraging is not (*p* = 0.7, default value) (Fig. S4e). Although unlikely, this scenario might occur under strong inbreeding avoidance, with individuals preferring to mate with individuals that are born elsewhere.

### Patch isolation effects

The effects of patch isolation (or fragmentation) independent of habitat amount (or habitat loss) on ecological response variables such as species abundance and richness have been extensively discussed in the literature, where even positive effects of fragmentation have been found to be prevalent (Fahrig, 2003, 2013, 2017). We contribute to this discussion by showing that, under HL, patch isolation can have a positive effect on the local consumer density in the patch experiencing HL (patch 2) through reduced cross-patch competition, and a negative effect on density in the neighbouring patch (patch 1) through reduced cross-foraging opportunities (Fig. 4). On the landscape scale, which is generally where the fragmentation effects are measured (Fahrig, 2017), these opposing effects combined with overexploitation can lead to a positive effect of patch isolation on consumer densities at low HL intensity, a negative effect at intermediate, and no effect at higher HL intensity (Fig. 4). However, this effect can become less prominent when consumers are evolving or completely disappear when the system is outside the overexploitation regime (Fig. 4; Fig. S9). Our study differs from other studies in that HL in our study affects only one of the patches directly, whereas in previous studies, the remaining habitat is typically redistributed, such that all patches lose some habitat (Bonin et al., 2011; Fahrig, 2017).

Current evidence for habitat fragmentation effects in conjunction with HL is inconsistent: theoretical studies predict negative effects at higher levels of HL (i.e. 20-30% habitat remaining) (Bascompte & Solé, 1996; Fahrig, 1998), whereas empirical studies show both positive and negative (but majorly positive) effects at all levels of habitat availability (Fahrig 2017, but see With 2016). Some of the positive effects of fragmentation have been ascribed to release from competition or predation (Fahrig, 2017). Our study supports this reasoning for observations in patch 2, and further adds the release from resource overexploitation as another possible mechanism for positive effects of isolation in patch 1.

### Effects of local HL on the landscape scale

HL is always detrimental for the consumer density at the landscape scale, irrespective of its effects at the local scale (Figs. 2, 4). Thus conserving as much habitat as possible should still be the priority. Furthermore, we also show the possibility of landscape level extinction when only one of the patches is experiencing resource loss, which can happen even when the consumers in the patch experiencing HL are already extinct (Fig. 5). This means that a naive management decision of allowing resources in a patch to be completely harvested because the species of interest (here consumers) does not produce any newborns in that patch can even cause landscape level extinction of the species. Hence, one should carefully consider the impacts of individual patch loss as it can potentially lead to landscape level collapse. Furthermore, other mechanisms of environmental deterioration (*sensu* Abrams, 2002) which can independently affect consumer demographic parameters (such as consumption rate, death rate, and resource conversion efficiency) should also be taken into account. For example, at lower per-capita consumer death rate the consumer population at the landscape level would still survive if one of the patch loses all the resources (Fig. 5c), but if external habitat deteriorating factors increases the death rate of the consumers then the same population would go extinct (Fig. 5g, k).

### Effects of eco-evolutionary dynamics

The eco-evolutionary model (heritable trait variation) showed both quantitative (Fig. 3) and qualitative (Figs. 4, 5) differences compared with the ecological model. Evolution of an increased consumption rate (*b*_0_) even led to a decrease in the equilibrium consumer densities since the system was in an overexploitation regime. Such a reduction in densities is also referred to as “adaptive decline” (Abrams, 2019). The constant variation scenario on the other hand yielded similar results as having no trait variation at all, except when the trait affects the consumer growth rates nonlinearly through Jensen’s inequality (Jensen 1906; Ruel and Ayres 1999; Fig. S8 and Fig. S11).

Heritable variation allowed the patch experiencing HL (patch 2) to survive higher *D* values as compared to other variation scenarios, through “evolutionary rescue” (as reviewed by Bell, 2017) (Fig. 4b; also see Fig. S11b, where the same is observed even if heritable variation leads to lower equilibrium consumer densities). Furthermore, heritable variation allowed complete rescue of patch 1 when other variation scenarios went extinct (Fig. 5, row 2). Evolutionary responses to HL and fragmentation have been studied theoretically, for example for evolution of dispersal distance (North et al., 2011) and coevolution of interacting species (Gawecka et al., 2022), as well as observed empirically. For example, selection of better colonization ability was observed in a large butterfly metapopulation in Finland (Fountain et al., 2016). However, there seems to be a lack of empirical evidence for such evolutionary responses specifically to HL to be rapid enough to save the population from extinction.

### Assumptions, potential improvements, and a suggestion for an empirical test

Our model makes several assumptions. We consider the two-patch consumer-resource system as the simplest scenario to create a local (patch level) and landscape scale. The within-patch foraging (*p*) and mating (*β*) parameters denote the traits that control the propensity to forage and mate in the patch of their birth. The patch boundaries do not have physical barriers so that the consumers can freely move in and out of the patch (similar to the “invisible” boundary in Cantrell et al. (1998)). Thus our model could be relevant to freely moving consumer populations such as fish, birds, and insects with resources confined in their respective habitat patches.

Unlike previous two-patch models with migration between patches (for example Jansen, 1995; Gomulkiewicz et al., 1999; Gyllenberg et al., 1999), our model does not assume direct transfer of population density between the patches. Instead, we have cross-patch mating where half of the progeny is put in the adjacent patch. We also keep identical parameter values for the two patches, except that HL happens only in one of the patches, to not confound the effects of HL with specific patch properties. However, traits in the two patches can evolve to different values in the evolutionary scenario. In principle, the patches could easily be made asymmetric based on other scenarios of interests such as source-sink dynamics (Gomulkiewicz et al., 1999). Another point to be noted is that we have chosen the parameter ranges where the model shows stable equilibrium dynamics. One could indeed find regions with fluctuating dynamics in our model (for example, Fig. S5) and extend the analysis in that direction (see, Jansen, 1995, 2001).

For simplicity, our study considers trait variation and evolution only in the consumer population, and the resource parameters are assumed to be constant. Given that the importance of studying the effects of environmental change under eco-evolutionary dynamics is increasingly being emphasized (Åkesson et al., 2021; Faillace et al., 2021), the above assumption could be relaxed in the future studies to consider consumer-resource coevolution in the current model. A further limitation of the current modeling approach is that only one trait can independently possess trait variation. If two uncoupled traits were to evolve, it could be done by considering a 2-dimensional grid of bins, each representing a trait combination. Such a system could straightforwardly be extended to *n* traits. Although, in principle this is possible, it is outside the scope of this study and might be more easily achieved using an individual-based model.

Our assumed system of inheritance, with offspring getting the mid-parent trait value, also plays an important role in determining the evolutionary outcome. For example, in the scenario where *p* and *β* are coupled, one could expect that, under HL, patch 1 should evolve to *p* = *β* = 1 (high within-patch dependence) and patch 2 should evolve to *p* = *β* = 0 (high cross-patch dependence). However, we do not observe such divergent evolution in this case, or in any other scenario in our model, because the cross-patch mating, mid-parent value assumption, and non-assortative (with respect to traits) mating together puts converging pressure on the evolving traits from the two patches. Dieckmann and Doebeli (1999) have shown such lack of evolutionary branching in an individual based model when there is sexual reproduction but without sufficient assortative mating.

Lastly, we propose a possibility to use the corals and coral-associated fish system used by Bonin et al. (2011) to empirically validate some of our predictions, specifically regarding the effects of HL in one patch on the neighbouring patch, the effects of patch isolation interacting with HL intensity, and the possibility of landscape scale extinction due to HL in a local patch. Bonin et al. (2011) try to disentangle the effects of HL and fragmentation (patch isolation), where live corals are removed to mimic HL and distance between experimental reefs is changed to manipulate patch isolation. In this setup, loss in live coral (resources) still leaves the coral beds which can be accessed by the fish (consumers) as breeding sites, which satisfies one of our important assumption. The amount of time spent in each patch for foraging and/or mating could be used as a proxy for their inherent preferences. Individual differences (variation) in these preferences and/or other consumer parameters (e.g. consumption rate) could further be used to understand the role of trait variation. Such empirical validation, either confirming or contradicting our theoretical predictions, would bring in the much needed dialogue between theory and experiment (Haller, 2014; Otto & Rosales, 2020; Joshi, 2022) as well as provide tangible predictions for future studies directly aimed at policy making.

## Conclusion

Better assessment and prediction of the effects of HL would help in planning well-informed conservation and management strategies. In our study we demonstrate how HL and patch isolation can show opposite effects at the local patches and how they translate to the landscape scale, highlighting that any assessment of HL effects depends on the spatial scale at which the assessment is performed. We also show how trait variation can provide better resistance to HL in the affected patch, and under certain conditions even rescue the populations at the landscape scale. Our study should motivate future studies as well as conservation efforts to simultaneously consider multiple spatial scales and trait variation while assessing and mitigating the effects of habitat loss.

## Supporting information

R code for generating data and figures

## Acknowledgments

We thank all the members of Theoretical Biology Group at Bielefeld University for lively discussions, and valuable inputs. This study was partially funded by research consortium SFB-TRR 212, project numbers 316099922 and 396782288.

