## Supplementary Information for "Multi-scale Effects of Habitat Loss and the Role of Trait Variation"

**Supplementary Information for:**
**Multi-scale effects of habitat loss and the role of trait variation**

Rishabh Bagawade, Koen J. van Benthem, Meike J. Wittmann

### S1 Effect of the shape of resource removal rate

We model habitat loss by removing resources in patch 2. The resources are removed with the resource removal rate $\mathcal{H}_{R_2}$ according to a logistic curve with steepness of the curve $s$, half saturation constant $t_{1/2}$, and maximum resource removal rate $D$ according to equation (3) in the main text. Out of these parameters, we investigate the effects of $D$ in the main text (Fig. 4). In this subsection, we look at the effects of different values of $s$ and $t_{1/2}$ under default parameter settings (table 1).

Fig. S1 shows the $\mathcal{H}_{R_2}$ curves for different values of $t_{1/2}$ and $s$. Fig. S2 shows the corresponding time series plots of consumer densities under HL for the three variation scenarios. Barring differences in transient dynamics, the equilibrium consumer densities are not affected by $s$ and $t_{1/2}$ at least under default parameter values and the considered range of $s$ and $t_{1/2}$. We choose $s = 0.003$ and $t_{1/2} = 3500$ as the default values for these two parameters, and do not change them in our analysis.

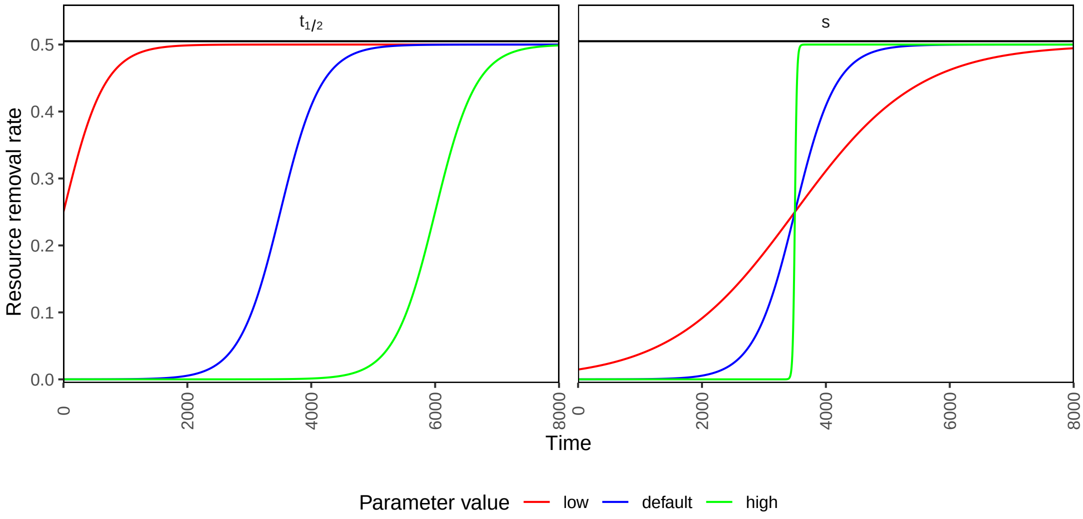


Figure S1: Plot of resource removal rate $\mathcal{H}_{R_2}$ using eq. (3) from the main text, where three example values of $t_{1/2}$ and $s$ are used. The values low, default and high correspond to 0, 3500 and 6000 respectively for $t_{1/2}$, and 0.001, 0.003 and 0.061 respectively for $s$. When $t_{1/2}$ is changed from default then $s$ is kept as default, and vice versa. All other parameter values are default values from table 1. As observed, $t_{1/2}$ denotes time when resource removal rate is half of maximum, and $s$ denotes the steepness of increase in resource removal rate.

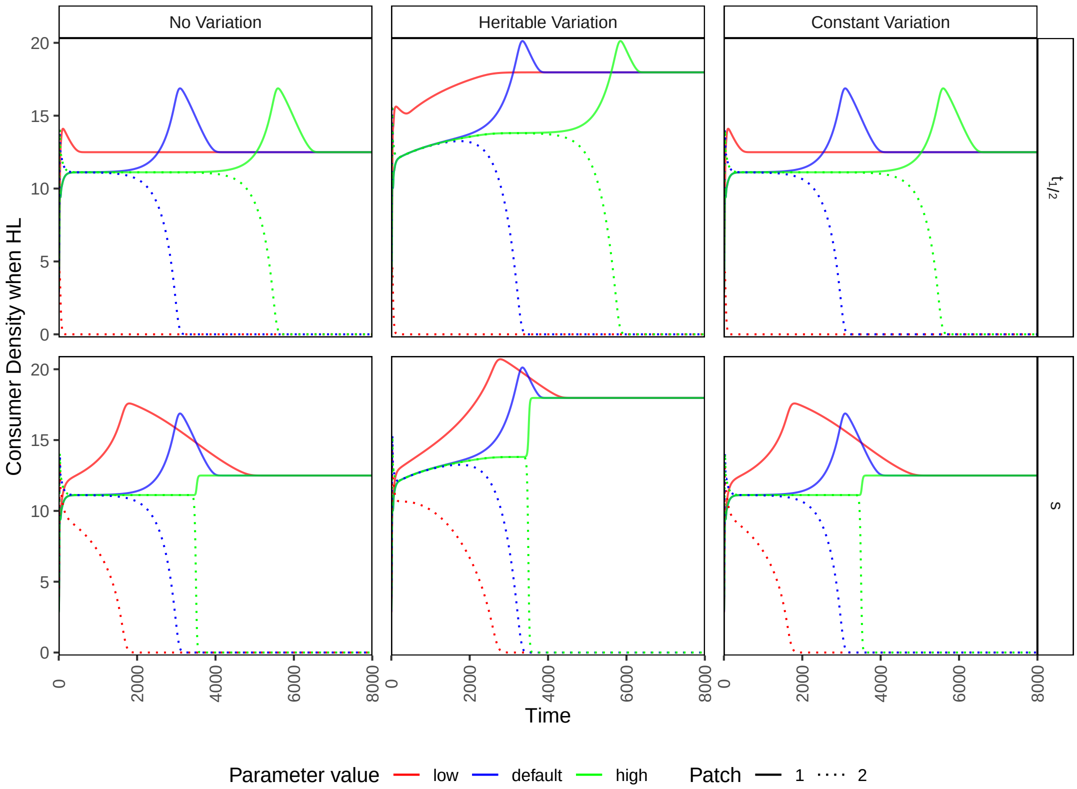


Figure S2: Time series plot for consumer densities of both patch 1 and patch 2 under HL are shown for the three variation scenarios. Three different values used for $t_{1/2}$ and $s$ are as defined in Fig. S1. The evolving trait is resource conversion efficiency $\epsilon$ with initial mean $TT_{mean} = 0.2$, and standard deviation $TT_{sd} = 0.08$ with maximum possible trait value $TT_{max} = 0.4$, and minimum $TT_{min} = 0$. Observe that the equilibrium consumer densities are identical irrespective of the $t_{1/2}$ and $s$ values considered.

### S2 Inheritance Description

To calculate the offspring trait distribution, we start with defining four $B \times B$ matrices: $A^{11}$, $A^{12}$, $A^{22}$, and $A^{21}$. Here, each element $A^{mn}_{ij}$ of the matrix $A^{mn}$ represents the density of newborns produced by females in bin $i$ of patch $m$ by mating with males in bin $j$ of patch $n$. Note that $m, n \in \{1, 2\}$ and $i, j \in \{1, 2, ..., B\}$. We obtain these matrices using an outer product:

$$A^{mn} = \mathcal{Y}^{mn} \bigotimes \mathcal{W}^{n}. \tag{S1}$$

Here $\mathcal{Y}^{mn}$ is the vector denoting newborn density produced by females in each bin in patch $m$ by mating with males in patch $n$. Note that the vectors $\mathcal{Y}^{11}$, $\mathcal{Y}^{12}$, $\mathcal{Y}^{22}$, and $\mathcal{Y}^{21}$ correspond to the terms $q_{11}\mathcal{F}_1$, $q_{12}\mathcal{F}_1$, $q_{22}\mathcal{F}_2$, and $q_{21}\mathcal{F}_2$ respectively, where each element of the vector corresponds to the newborn density produced according the trait values assigned to the index of that element in the vector. $\mathcal{W}^n$ is the vector denoting the population densities of each bin divided by the total population of the patch $n$. Given the even sex ratio and random (trait independent) mating,

$\mathcal{W}^n$ gives us the proportion of males coming from each bin. In other words, $\mathcal{Y}^{mn}$ gives us the trait-dependent offspring contribution by females and $\mathcal{W}^n$ gives us the trait-independent genetic contribution by males i.e. bins with more males contribute more in the inheritance simply due to their high density.

As an example, let us define the vectors $\mathcal{Y}^{11}$, $\mathcal{Y}^{12}$, and $\mathcal{W}^1$. Vectors for patch 2 are then defined analogously. The $b^{th}$ element of $\mathcal{Y}^{11}$ where $b \in \{1, 2, ..., B\}$ is defined as:

$$\mathcal{Y}_b^{11} = q_{11,b} \cdot \epsilon_b \cdot \mathcal{A}_{1,b} \cdot (\mathcal{C}_{11,b} + \epsilon_{c_b} \cdot \mathcal{C}_{12,b}) \cdot n_{1,b}. \tag{S2}$$

Here, the subscript $b$ denotes the value of that parameter or expression calculated using the trait value associated to that bin. Note that identical parameter values are assigned to each bin for a non varying trait, and equidistant values between minimum ($TT_{min}$) and maximum ($TT_{max}$) trait values are assigned to each bin in case of the trait having variation. Further, $q_{11,b} = \beta_b N_1/(\beta_b N_1 + (1 - \beta_b)N_2)$ and $\mathcal{A}_{1,b} = 1 - e^{-\theta_b(\beta_b \frac{N_1}{2} + (1-\beta_b)\frac{N_2}{2})}$. Note that the consumer densities used here are not bin-specific (i.e. $n_{1,b}$ and $n_{2,b}$) but of the whole patches (i.e. $N_1$ and $N_2$) because the mate finding applies to the whole patch and not to the specific bins. $\mathcal{C}_{11,b}$ and $\mathcal{C}_{12,b}$ are defined similar to eq. (2) in the main text with bin specific parameter values (i.e. $b_{0,b}$, $b_{1,b}$, and $p_b$) but full patch resource densities $R_1$ and $R_2$ since we assume resources do not possess trait variation and thereby do not have any bins associated to them. Similarly, $\mathcal{Y}_b^{12}$ is defined as:

$$\mathcal{Y}_b^{12} = q_{12,b} \cdot \epsilon_b \cdot \mathcal{A}_{1,b} \cdot (\mathcal{C}_{11,b} + \epsilon_{c_b} \cdot \mathcal{C}_{12,b}) \cdot n_{1,b}. \tag{S3}$$

Lastly, $b^{th}$ element of vector $\mathcal{W}^1$ is defined as:

$$\mathcal{W}_b^1 = \frac{n_{1,b}}{N_1}, \tag{S4}$$

where $N_1 = \sum_{b=1}^{b=B} n_{1,b}$. The vectors $\mathcal{Y}^{22}$, $\mathcal{Y}^{21}$, and $\mathcal{W}^2$ can be defined analogously. The $A^{mn}$ matrices can then easily be calculated using eq. (S1).

Having defined the four $A$ matrices, we have to assign each element $A_{ij}$ to the bin with the mid-parent trait value. Accordingly, the value goes to bin number $(i + j)/2$ when $i + j$ is even and is split equally between the two bins around $(i + j)/2$ when $i + j$ is odd. In order to do this for all the bins, we define a set of $B \times B$ matrices $\mathcal{M}_b$, where $1 \leq b \leq B$. The elements of matrix $\mathcal{M}_b$, i.e. $\mathcal{M}_{b,ij}$, are defined as follows:

$$\mathcal{M}_{b,ij} = \begin{cases} 1, & \text{if } \frac{i+j}{2} = b \\ 1/2, & \text{if } \frac{i+j}{2} = b \pm 1/2 \\ 0, & \text{Otherwise} \end{cases} \tag{S5}$$

where $1 \leq i, j \leq B$.

We get the population growth rate, for bin $b$, with heritable trait variation as follows:

$$\begin{aligned} sum(A^{11} \circ \mathcal{M}_b) + sum(A^{12} \circ \mathcal{M}_b)/2 + sum(A^{21} \circ \mathcal{M}_b)/2 = f_{1,b}, \\ sum(A^{22} \circ \mathcal{M}_b) + sum(A^{21} \circ \mathcal{M}_b)/2 + sum(A^{12} \circ \mathcal{M}_b)/2 = f_{2,b}. \end{aligned} \tag{S6}$$

Here, $\circ$ denotes element-wise multiplication of the matrices, $sum()$ represents the sum of all the elements in the matrix, and $f_{i,b}$ is the growth rate for bin $b$ in patch $i$.

### S3 Analytical treatment of the simplified model

In this section we find equilibrium points of a simplified version of the ecological model from the main text. There are three important assumptions made during the simplification process. First, $\theta$ is assumed to be very large thus making the mate finding Allee effect terms equal to 1 (i.e. $(1 - e^{-\theta(\beta\frac{N_1}{2}+(1-\beta)\frac{N_2}{2})}) \to 1$ and analogously for the other patch) i.e. there is no Allee effect. This simplified model should be a good approximation to the full model for our default parameter values (where $\theta = 2$) since there is no qualitative effect of consumer densities for $\theta$ values greater than approximately 0.5 (see equilibrium consumer density over the parameter space of $\theta$ in Fig. S7). Second, we focus on scenarios where the habitat loss (HL) case implies $N_2, R_2 \to 0$ i.e. patch 2 goes extinct, and the no habitat loss (No HL) case implies $N_1 = N_2 = N$ and $R_1 = R_2 = R$ i.e. both patches reach same equilibrium points owing to symmetry of the system. Note that for the HL case, patch 2 might not go extinct when $\beta$ is small (Fig. S4e) or $D$ is very small (Fig. S7), and for the No HL case there can also be cases with asymmetric alternative stable states even in symmetric patches (see van Benthem & Wittmann, 2020) or cases where consumers in one of the patch out-compete the other (Fig. 5 and Fig. S12); these then cannot be captured by this analytic approximation. Lastly, the consumption term is linearized from the original type-II functional response, i.e. there is no consumer saturation. For example, the consumption terms $\mathcal{C}_{11} = \frac{b_0 p R_1}{b_1 + pR_1 + (1-p)R_2}$ and $\mathcal{C}_{12} = \frac{b_0(1-p)R_2}{b_1 + pR_1 + (1-p)R_2}$ become $\mathcal{C}_{11} = \frac{b_0 p R_1}{b_1}$ and $\mathcal{C}_{12} = \frac{b_0(1-p)R_2}{b_1}$ respectively. Apart from this, $\epsilon_c \in [0, 1]$ is considered to be 1 for simplicity but all the qualitative claims still hold, at least in the default scenarios, if it is less than 1 (see equilibrium consumer density over parameter space $\epsilon_c$ in Fig. S7, and compare Fig. 2 with Fig. S6 where $\epsilon_c = 1$ in the latter).

For the HL case, according to the above assumptions, $N_2, R_2 \to 0$ hence $\mathcal{F}_2, q_{12} = 0$, $q_{11} = 1$, and $C_{12} = C_{22} = 0$. Then the system (derived from eq. (8) and eq. (4)) looks as follows:

$$\begin{aligned} \frac{dN_1}{dt} &= \epsilon r R_1 \frac{N_1}{2} - a_0 N_1, \\ \frac{dR_1}{dt} &= r_0 R_1 (1 - \frac{R_1}{k}) - r R_1 N_1, \end{aligned} \tag{S7}$$

where $r = \frac{b_0}{b_1} p$, which we call the effective per-capita consumption rate.

For the No HL case, according to the above assumptions, $N_1 = N_2 = N$ and $R_1 = R_2 = R$ hence $\mathcal{F}_1 = \mathcal{F}_2$, $q_{11} = q_{22} = \beta$, $q_{12} = q_{21} = 1 - \beta$, $C_{11} = C_{22}$, and $C_{12} = C_{21}$. This leads to the system for patch 1 to be same as eq. (S7) except $r = \frac{b_0}{b_1}(p + (1-p)) = \frac{b_0}{b_1}$ i.e. put $p = 1$ in the HL case. Henceforth we will use $r$ to denote per-capita effective consumption rate, and it's value will be $\frac{b_0}{b_1} p$ in the HL case and $\frac{b_0}{b_1}$ in the No HL case. Note that since $0 \le p \le 1$, the effective consumption rate is lower in the HL case compared to the No HL case.

The model in eq. (S7) is the same as the consumer-resource model with biotic resource growth in Abrams (2019) (see their system with eq. $1a$ and $2b$) barring the differences in notation. The equilibrium points with positive consumer and resource densities are stable in that model as long as resource growth is density dependent (i.e. $r_0/k \neq 0$ in our model). Consequently, we also get the same dynamics for the equilibrium points. However, we still derive the equilibrium points below since it will be helpful to interpret the effect of habitat loss in our simplified two patch system.

The equilibrium points for habitat loss (HL) case can be calculated as follows:

Putting the rate of change of consumer population to zero we get for the nontrivial (coexistence) equilibrium:

$$\frac{dN_1}{dt} = \epsilon r R_1 \frac{N_1}{2} - a_0 N_1 = 0$$

$$\implies R_1^* = \frac{2a_0}{\epsilon r} \tag{S8}$$

Putting the rate of change of the resource to zero we get:

$$\frac{dR_1}{dt} = r_0 R_1 (1 - \frac{R_1}{k}) - r R_1 N_1 = 0$$

$$\implies N_1^* = \frac{r_0}{r}(1 - \frac{R_1^*}{k})$$

$$\implies N_1^* = \frac{r_0}{r}(1 - \frac{2a_0}{\epsilon r k}) \tag{S9}$$

The other equilibrium points are $N_1^* = 0, R_1^* = 0$ and $N_1^* = 0, R_1^* = k$. The non-trivial equilibrium points for no Habitat Loss (No HL) case are the same as above with $p = 1$ i.e. $r = \frac{b_0}{b_1}$.

Note that $R_1^* = \frac{2a_0}{\epsilon r}$ (eq. (S8)), and for the HL case $r = \frac{b_0}{b_1}p$ which is always less than for the No HL case $r = \frac{b_0}{b_1}$ since $0 \leq p \leq 1$. This implies $R_1^*$ in the habitat loss case is always greater than in the no habitat loss case. This can be observed in Fig. 2, where patch 1 resources are greater in the presence of HL than in the absence of HL for both low and high per-capita consumer death rate ($a_0$), whereas patch 1 consumer density is greater in the presence of HL than in the absence for lower $a_0$ and the other way round for higher $a_0$. The mechanism behind such observation for consumers is explained in the following section.

### S4 Overexploitation and its effect on patch 1 consumers

In a consumer resource system, the naive expectation is that higher per-capita consumption of resources should lead to higher consumer densities. But when the system is in an overexploitation regime, higher consumption leads to lower consumer densities. Abrams (2002, 2009, 2019) have discussed this phenomenon of overexploitation for consumer-resource systems. Recall that our simplified model is conceptually identical to the biotic resource growth model in Abrams (2019). However, it still differs from it in terms of habitat loss and spatial two patch interpretation. In this section we discuss how overexploitation can explain the increase or decrease observed in patch 1 consumer density after habitat loss in patch 2.

Let us consider the simplified model developed in the previous section (SI S3). The equilibrium patch 1 consumer density ($N_1^*$) as a function of $r$ based on eq. (S9) is shown in Fig. S3 (see default case). This function has a peak when $\frac{\mathrm{d}N_1^*}{\mathrm{d}r} = 0$, and hence when:

$$r_0(2\frac{2a_0}{\epsilon r^3 k} - \frac{1}{r^2}) = 0$$

$$\implies 2\frac{2a_0}{\epsilon r k} = 1$$

and thus the function peaks when:

$$r = \frac{4a_0}{\epsilon k}. \quad \text{(S10)}$$

Since $R_1^* = \frac{2a_0}{\epsilon r}$ (eq. (S8)), the equilibrium resource density at the peak of the $N_1^*$ vs $r$ curve is:

$$R_1^* = \frac{k}{2}. \quad \text{(S11)}$$

Now, we realize that the energy flow into the system is solely governed by the resource growth term:

$$r_0 R_1(1 - \frac{R_1}{k}),$$

all other terms in in eq. (S7) concern either outflow terms (death of consumers) or conversion terms (from resource to consumers). The energy influx is maximal with respect to the amount of resources present, when:

$$\frac{\mathrm{d}}{\mathrm{d}R_1}\left(r_0 R_1(1 - \frac{R_1}{k})\right) = 0$$

$$\implies 1 - \frac{2R_1}{k} = 0$$

$$\implies R_1 = \frac{k}{2}. \quad \text{(S12)}$$

Hence we see that the consumer population size at equilibrium is maximized when the effective consumption rate is such that the resources are kept at their maximum growth rate. On either side of the peak, the resource growth rate is not optimal which leads to lower equilibrium consumer densities. On the left side of the peak in Fig. S3, increase in consumption increases the equilibrium consumer density. In contrast, on the right side of the peak, one can observe that the equilibrium consumer density goes down with an increase in effective consumption rate because higher consumption pushes resource density below the values at which it can optimally grow (overexploitation regime).

Furthermore, from eq. (S9), $r > \frac{2a_0}{\epsilon k}$ if $N_1^* > 0$. This implies the effective consumption rate required for consumer density to peak ($r = \frac{4a_0}{\epsilon k}$) is just two times the minimum effective consumer density required for the consumers to exist (also shown in Abrams, 2002). This implies the system is in overexploitation regime over most of the parameter range of $r$.

Now recall from the previous section (SI S3) that the no habitat loss (No HL) case is the same as the habitat loss (HL) case except $p = 1$. Thus the effective consumption rate is $r = \frac{b_0}{b_1}p$ for the HL case and $r = \frac{b_0}{b_1}$ for the No HL case. Since $0 \leq p \leq 1$, $r$ goes down from No HL case to HL case. This is shown in Fig. S3 by red and blue vertical lines. The intersection between the vertical lines and the curves tells us the equilibrium consumer density at that particular $r$ value. Hence one can observe that the equilibrium consumer density of patch 1 goes up after habitat loss when the system is in the overexploitation regime and goes down otherwise. The default case is in the overexploitation regime and it takes its parameter values from the table 1. Changing in some of the parameter values (specifically higher $a_0$, lower $b_0$, and lower $\epsilon$) can shift the curves such that the system is no longer in the overexploitation regime for the given $r$ (see Fig. S3, where $N_1^*$ increases after HL in the default case and decreases in the other cases). This also explains the observations of patch 1 consumer density in Fig. 2a,e, and Fig. S4a,b,c where $N_1^*$ after HL is lower than without HL for higher $a_0$, lower $b_0$, and lower $\epsilon$ respectively.

Note that the explanation provided here is for the simplified model, therefore the Fig. S3 cannot be directly applied to the full model in the main text but can provide qualitative conceptual understanding of the mechanism behind before and after HL equilibrium consumer density patterns in patch 1. As an example, we assume in our approximation that the efficiency of cross-patch foraging $\epsilon_c = 1$. However, $\epsilon_c < 1$ can also potentially contribute in similar direction as the over-exploitation effect where patch 1 consumer density under HL is more than without HL (blue line below red line). Similarly, lower $\epsilon_c$ also has the potential to reduce the cross-patch foraging efficiency thereby reducing the patch 1 consumer density in absence of HL i.e. lowering the blue line (See Fig. S7 for parameter space $\epsilon_c$). However, this effect is very weak in our default scenarios ($\epsilon_c$=0.9).

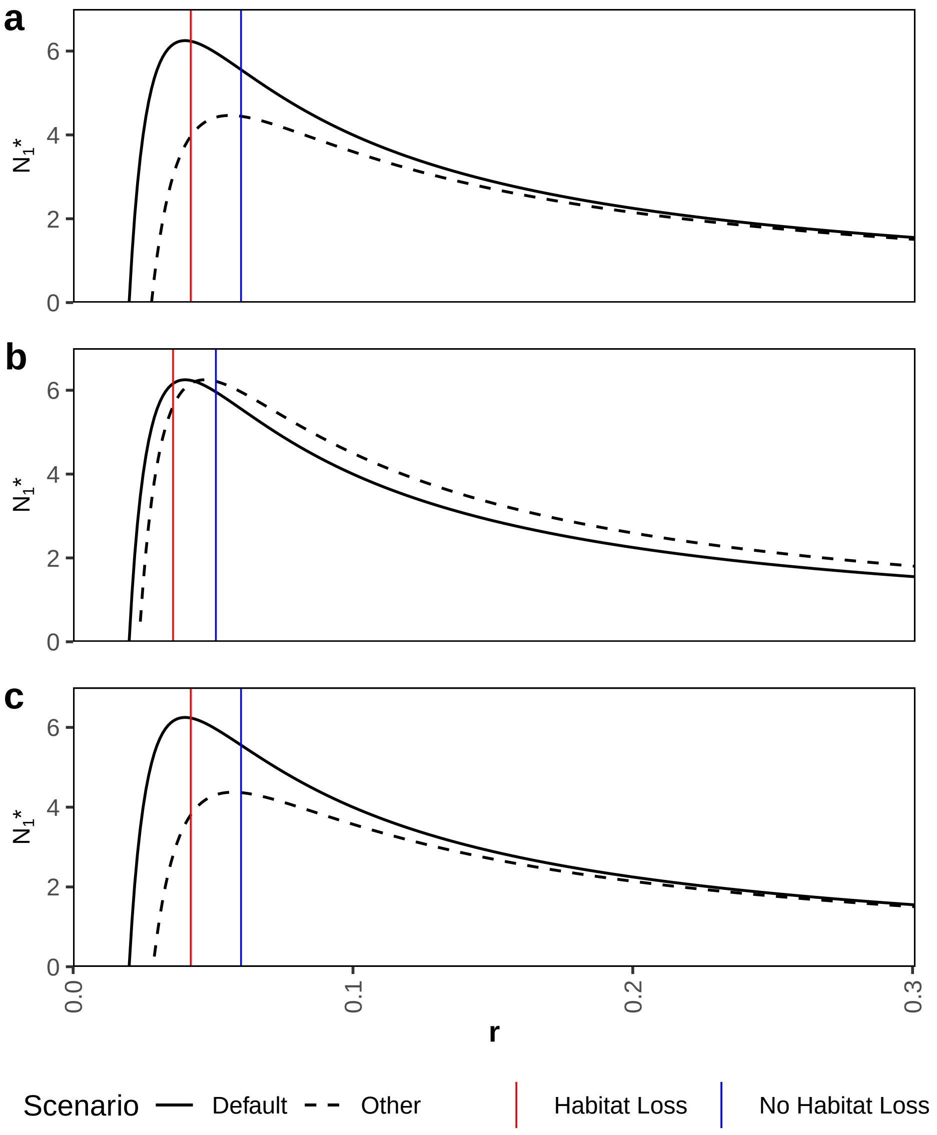


Figure S3: Equilibrium consumer density $N_1^*$ in the simplified model (eq. (S7)) as a function of effective consumption rate $r$, where from eq. (S9) $N_1^* = \frac{r_0}{r}(1 - \frac{2a_0}{\epsilon rk})$. The default curves (solid black lines) are based on the parameter values from table 1. The other curves (dashed black lines) are for $a$. high death rate $a_0 = 0.14$, $b$. low maximum per capita consumption rate $b_0 = 25$, and $c$. low resource conversion efficiency $\epsilon = 0.14$ (while keeping the other parameters the same as in the default case). The vertical lines denote the example $r$ values when there is no habitat loss $r = \frac{b_0}{b_1}$ (blue lines), and when there is habitat loss $r = \frac{b_0}{b_1} \cdot p$ where $p = 0.7$ (red lines).

### S5 Consumer dynamics for ecological model over certain parameter ranges

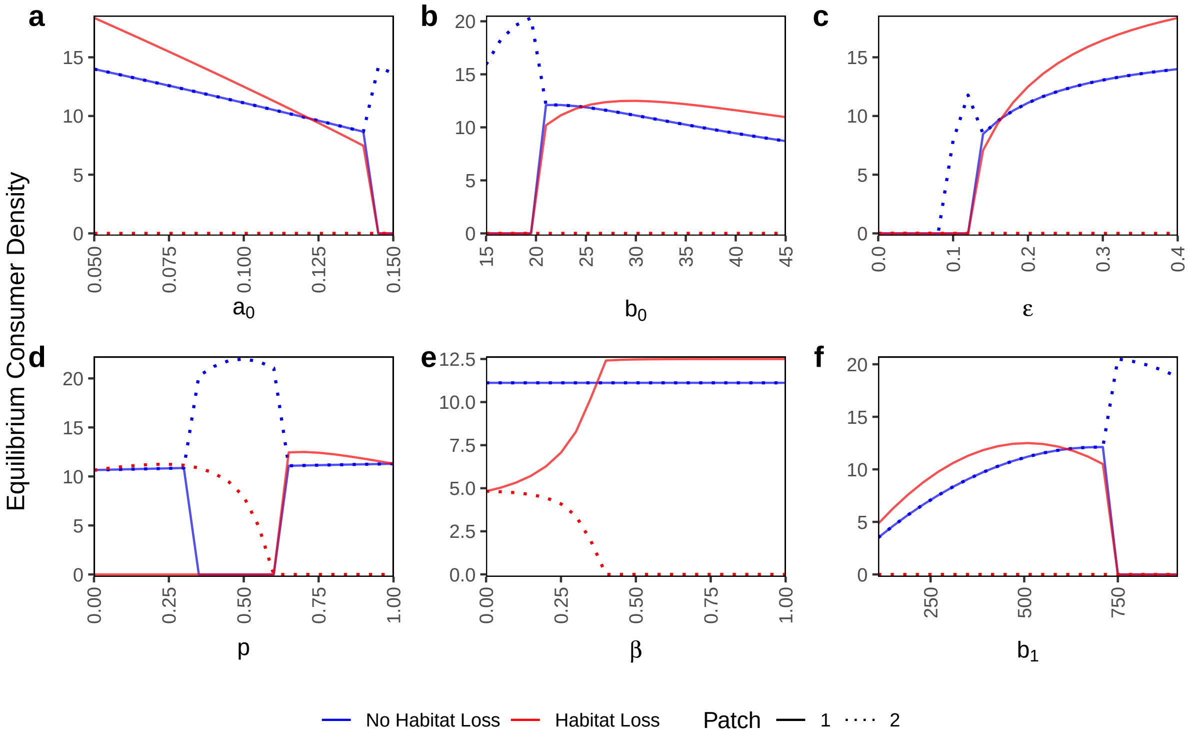


Figure S4: Patch-wise equilibrium consumer densities for the ecological model (no trait variation) over certain parameter ranges of: a. per-capita death rate $a_0$, b. maximum consumption rate $b_0$, c. resource conversion efficiency $\epsilon$, d. within-patch foraging preference $p$, e. within-patch mating preference $\beta$, and f. half-saturation constant for resource consumption $b_1$. Note that the equilibrium densities are calculated for 21 equidistant points in each parameter range but they are depicted by lines for better clarity. All other parameters are at their default values from table 1.

Whether HL in patch 2 increases or decreases the equilibrium density in patch 1 depends on the values of several parameters. This is shown in Fig. S4 by the crossing of the lines for patch 1 equilibrium density for the scenarios with HL (red solid lines) and those without (blue solid lines) for the parameters $a_0$, $b_0$, $\epsilon$, $\beta$, and $b_1$ (Fig. S4a, b, c, e, f). For $a_0$, $b_0$, $b_1$, and $\epsilon$ this effect can be explained by the system coming out of the overexploitation regime for higher $a_0$ (also see Fig. 2e) and $b_1$, and lower $b_0$ and $\epsilon$ (see SI S4). For $\beta$ this happens because, at lower values of $\beta$, consumers in both the patches survive after HL however patch 1 consumers have to suffer in this case since consumers in both patches are now dependent on patch 1 resources. This switching not only happens between HL and no HL scenarios, but can also occur due to the change in parameters themselves. For example in Fig. S4b, under HL, the equilibrium density in patch 1 (red solid line) goes up with increasing $b_0$ at intermediate values ($\sim$20-30) and down with increasing $b_0$ at higher values showing the overexploitation effect in the latter case. A similar observation can be made for $b_1$ (Fig. S4f); however, here the overexploitation happens at lower $b_1$.

In certain parameter regions, we observe outcompetition by the consumers that have higher initial density (i.e. patch 2 consumers). Particularly in the regions where the consumers are less efficient i.e. higher values of $a_0$ and $b_1$ (Fig. S4a and f), and lower values of $b_0$ and $\epsilon$ (Fig. S4b and c). Similar outcompetition also occurs at $p$ values between approximately 0.3 and 0.6 (Fig. S4d). At more extreme values of $p$, the larger initial population size in patch 2 no longer leads to outcompetition. Instead, under HL, the patch that forages least in patch 2 and most in patch 1 survives; this will be patch 1 when $p$ is high and patch 2 when it is low. Apart from outcompeting each other, the consumers in the two patches can also help each other survive by subsidizing the other patch because half of their offspring from cross-patch mating is put in the other patch. This happens at lower values of within-patch mating preference $\beta$, where both patches survive after HL (Fig. S4e).

### S6 Additional Figures

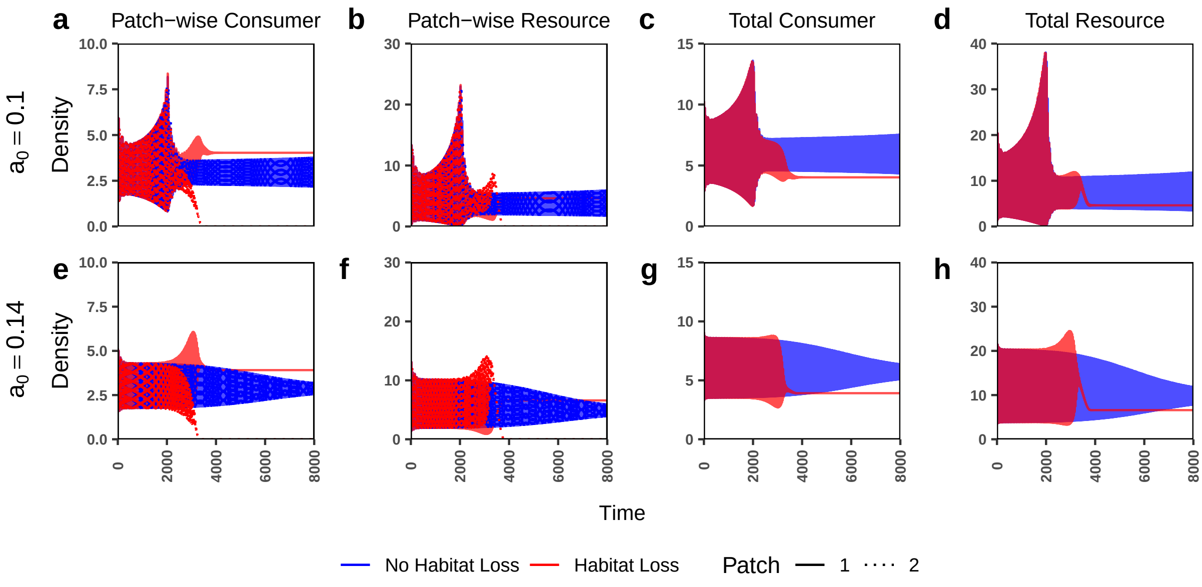


Figure S5: Figure identical to Fig. 2 in the main text, except here the half saturation constant for resource consumption is $b_1 = 90$. Fluctuating dynamics can be observed for the no HL case. These fluctuations are stabilized in this case due to HL and a dampening of the fluctuations is observed for higher per-capita consumer death rate ($a_0 = 0.14$).

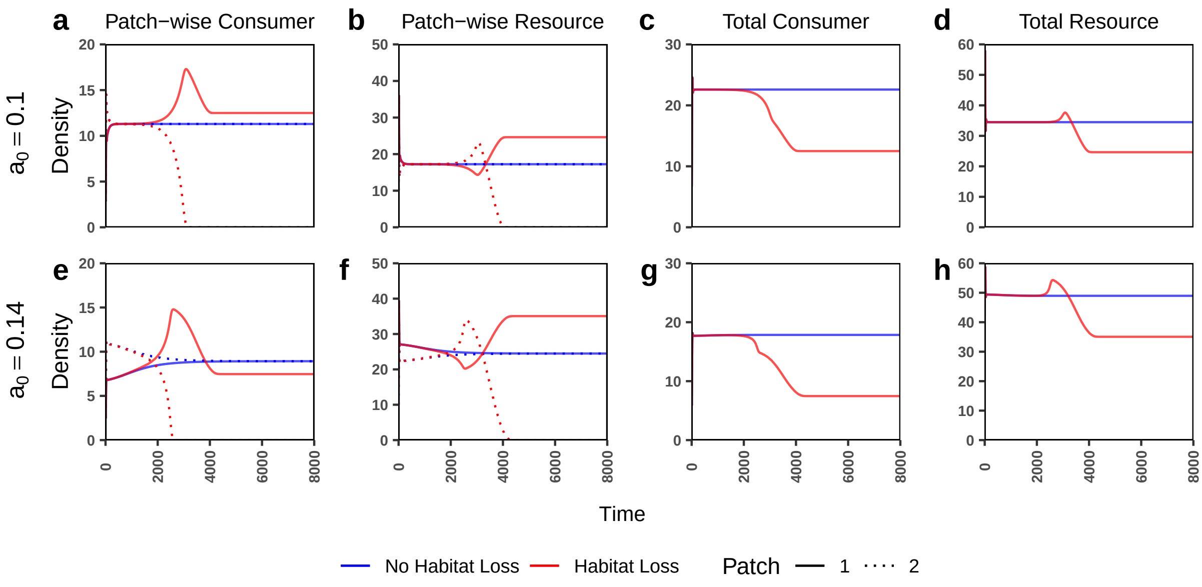


Figure S6: Figure identical to Fig. 2 in the main text, except here the efficiency if cross-foraging is $\epsilon_c = 1$.

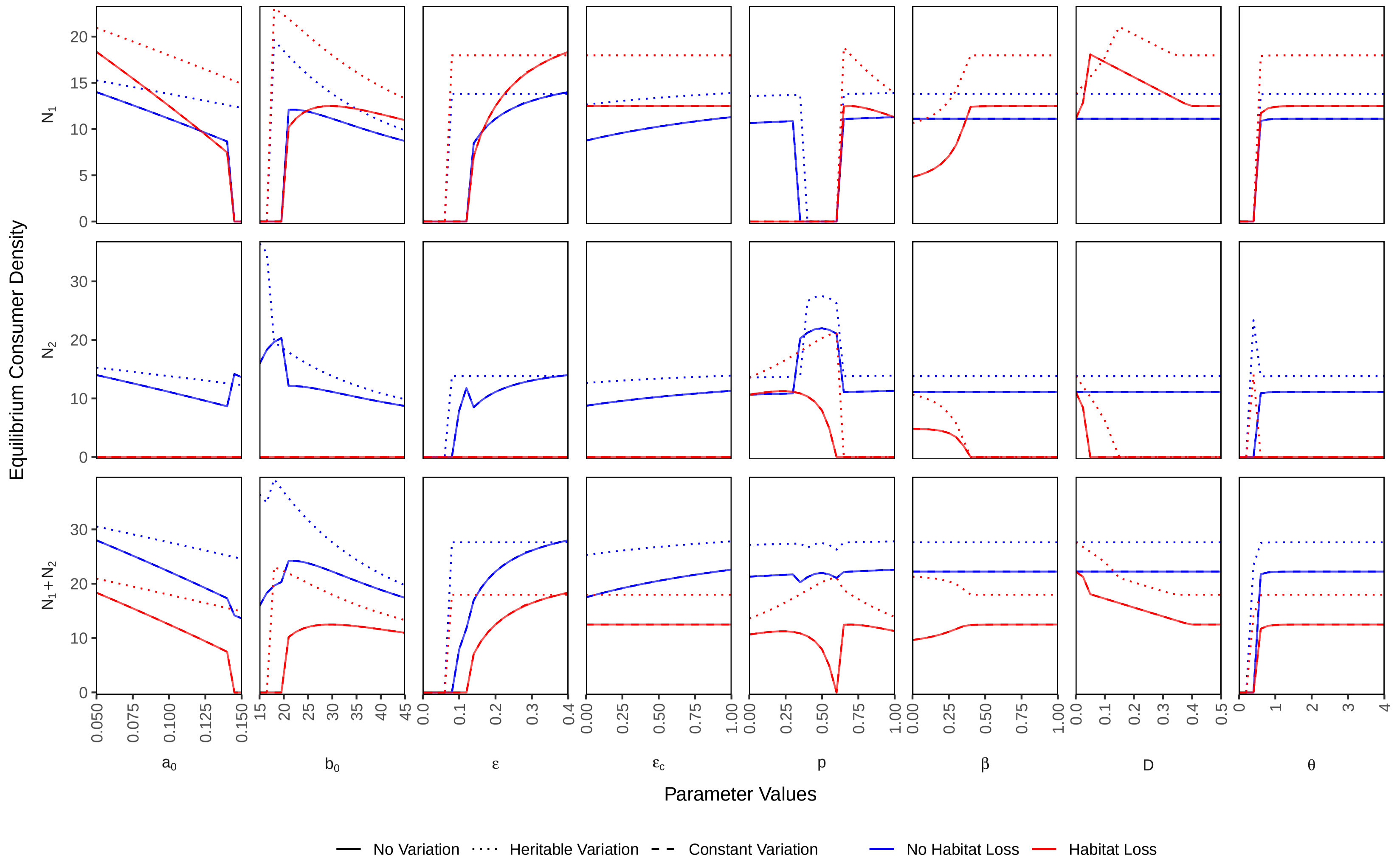


Figure S7: Individual patch and landscape level equilibrium consumer densities for the three trait variation scenarios compared over different parameter ranges. The varying (or evolving) trait is the resource conversion efficiency $\epsilon$ with initial mean $TT_{mean} = 0.2$, and standard deviation $TT_{sd} = 0.08$ with maximum possible trait value $TT_{max} = 0.4$, and minimum $TT_{min} = 0$. No variation and constant variation lines are overlapping. Note that the equilibrium densities are calculated for 21 equidistant points in each parameter range but they are depicted by lines for better clarity. All other parameter values are at their default values from table 1.

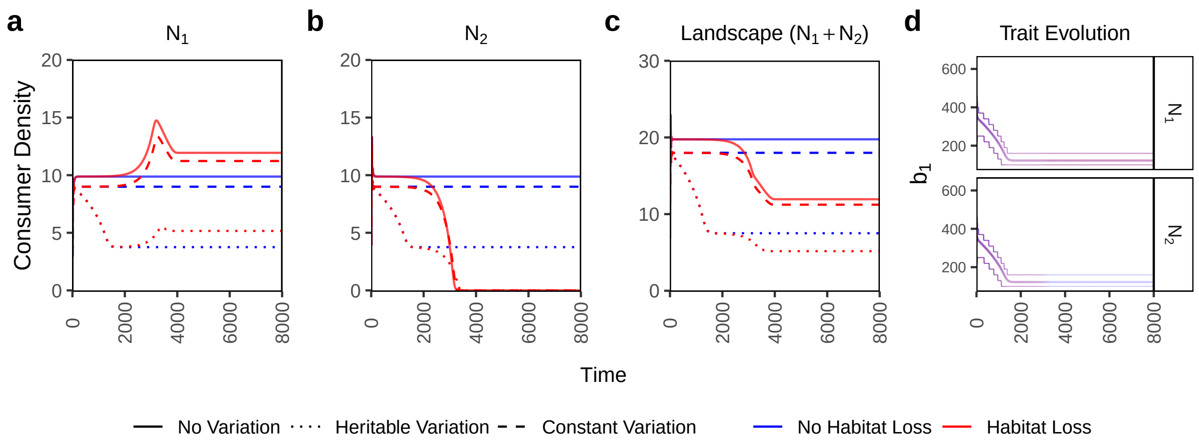


Figure S8: Time series plots for patch and landscape level consumer densities for the three variation scenarios in presence and absence of habitat loss. The evolving trait is the half saturation constant for resource consumption $b_1$ with initial mean $TT_{mean} = 400$, and standard deviation $TT_{sd} = 150$ with maximum possible trait value $TT_{max} = 700$, and minimum $TT_{min} = 100$. The last column shows the trait evolution where the central line denotes the mean trait value and the thinner outer lines denote the range which contains 90% of the density. Line transparency in the trait evolution plots denote consumer density. Note that the red and blue lines are overlapping in the trait evolution plots. All other parameter values are at their default values from table 1.

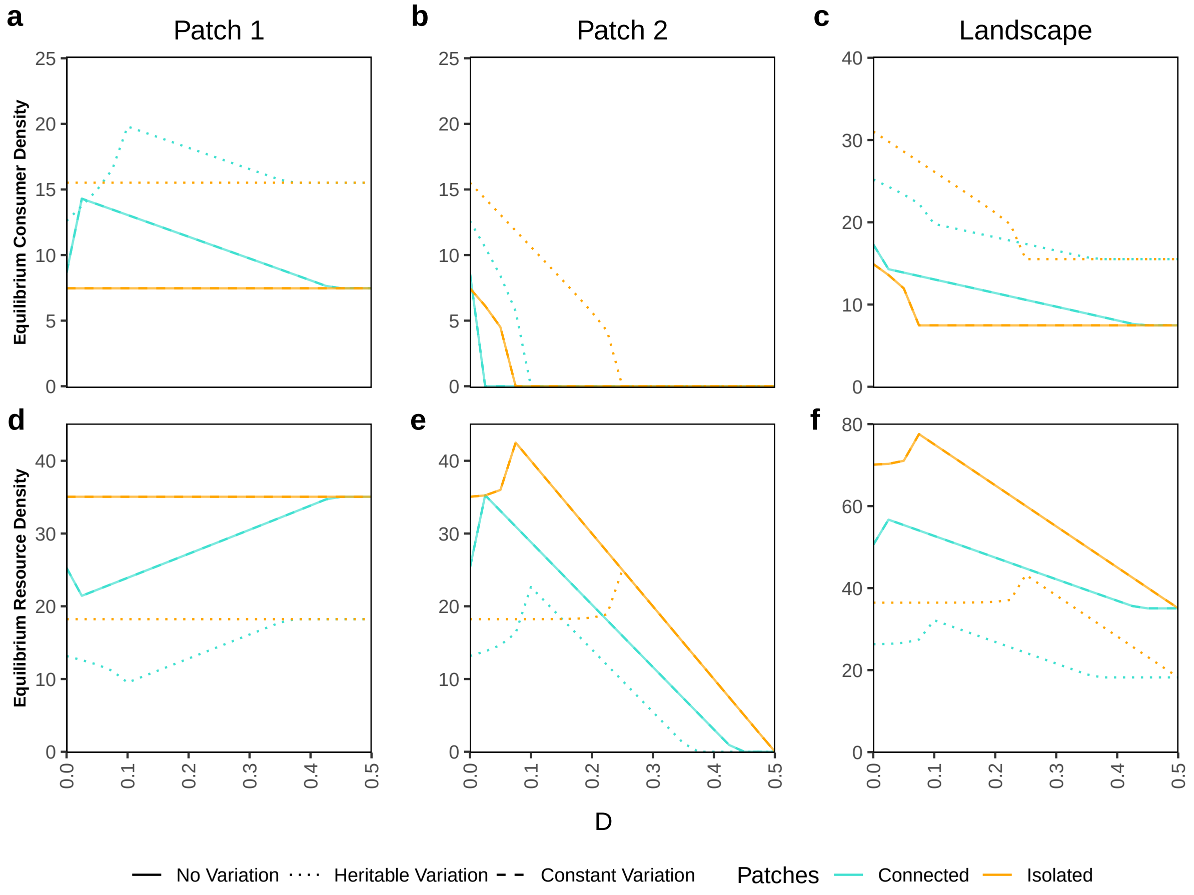


Figure S9: Figure identical to Fig. 4 in the main text, except the consumer death rate is higher ($a_0 = 0.14$) i.e. not in the overexploitation regime. Observe that unlike in Fig. 4c, for no and constant variation cases, the scenario with connected patches is harbouring higher consumer densities at the landscape scale at all values of $D$ (panel c). This happens because patch connectedness is always beneficial for patch 1 consumers (panel a), and patch 2 consumers go extinct at small $D$ values for both connected and isolated patches (panel b) nullifying the benefit of patch isolation at lower $D$ values. However, for the heritable variation case, the evolution of $\epsilon$ to higher values makes the consumers more efficient i.e. it leads to overexploitation. Thus the benefit of patch isolation at lower values of $D$ is amplified (panel a), and we regain the effect where orange and turquoise dotted lines cross each other (panel c).

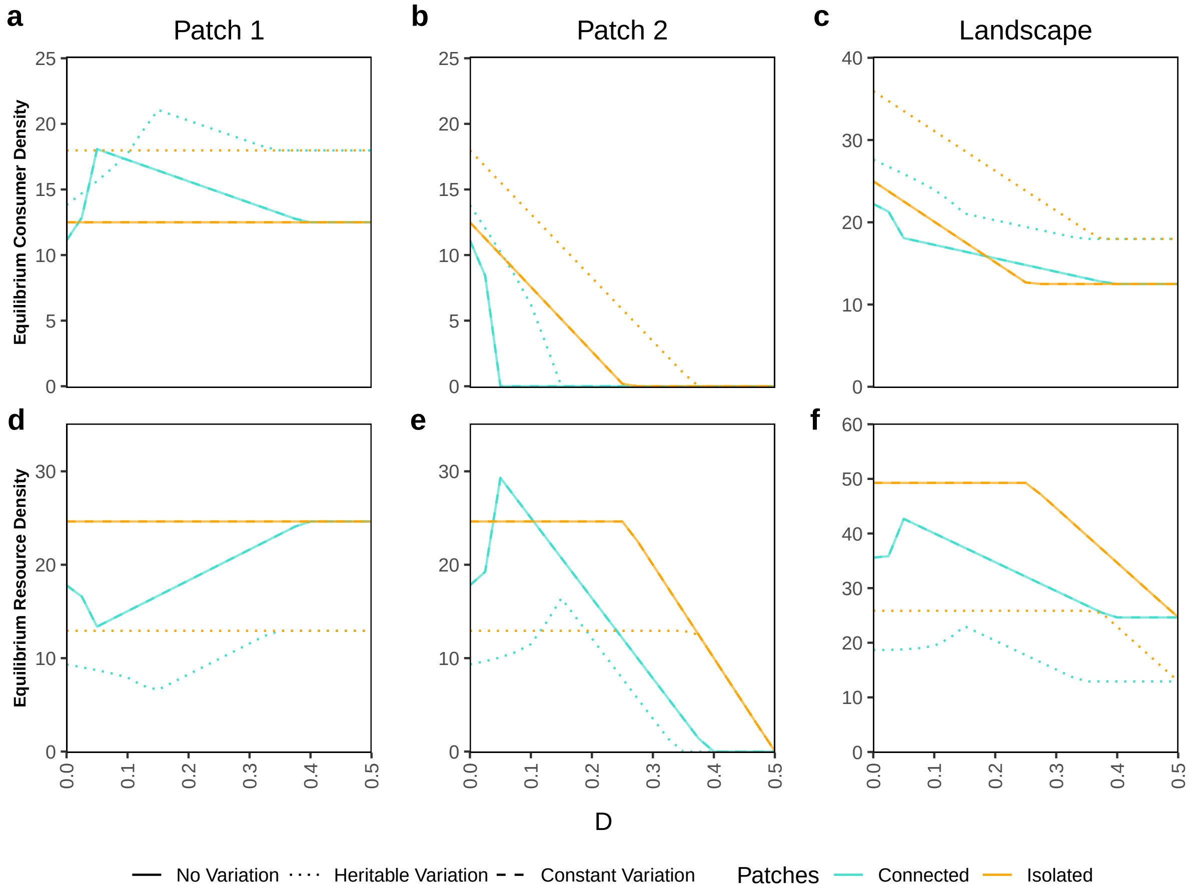


Figure S10: Figure identical to Fig. 4 in the main text, except there is no Allee effect i.e. the Allee effect term is set to 1 while running the numerical simulation. Comparing the orange lines from panel b with Fig. 4b, we observe that the Allee effect does cause extinction at lower HL intensity ($D$ value) for patch 2 consumers.

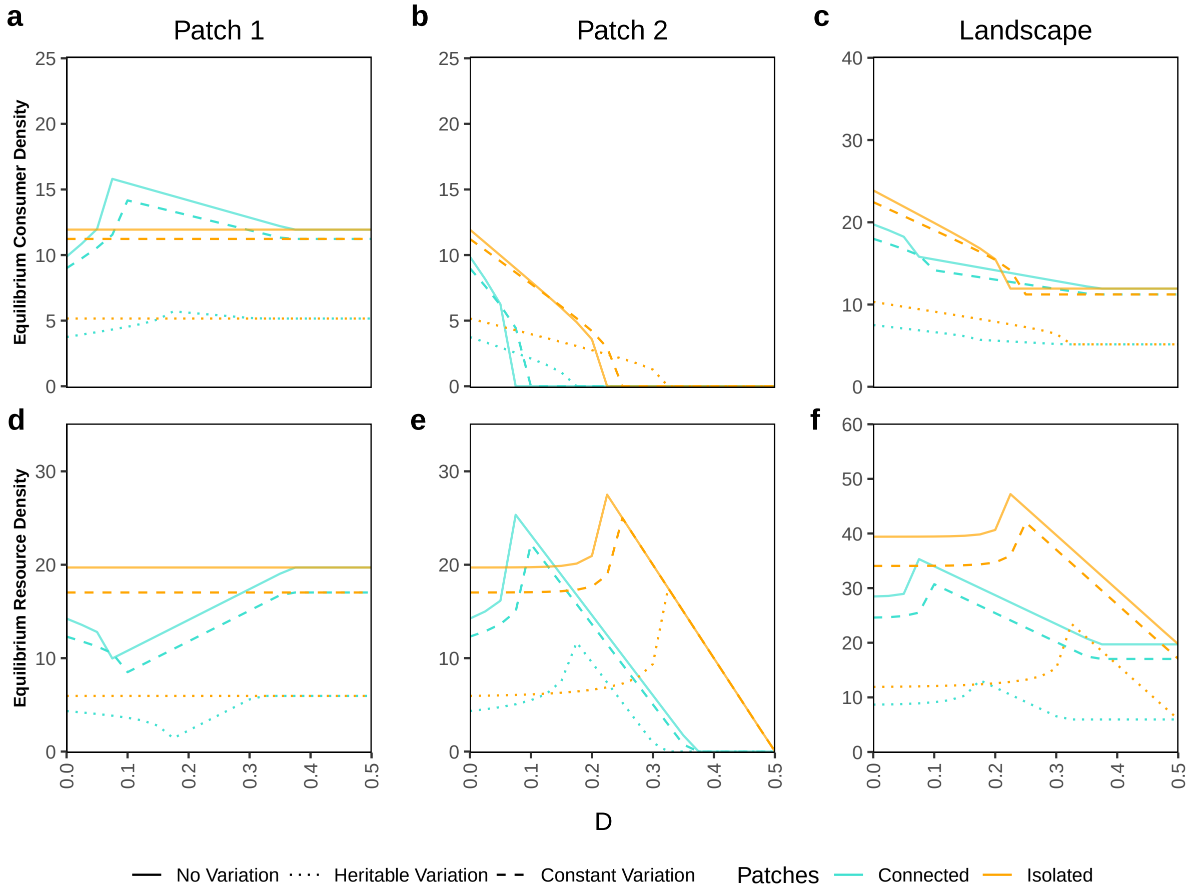


Figure S11: Figure identical to Fig. 4 in the main text, except that the varying (or evolving) trait is the half saturation constant for resource consumption $b_1$ with initial mean $TT_{mean} = 400$, and standard deviation $TT_{sd} = 150$ with maximum possible trait value $TT_{max} = 700$, and minimum $TT_{min} = 100$. Here, heritable variation leads to considerably lower equilibrium consumer densities at the landscape level, although it can still be beneficial for patch 2 consumers as it can help them to sustain higher $D$ values (panel b, dotted lines). The constant variation case deviates from the no variation case in terms of equilibrium density. Furthermore for patch 2 consumers, the constant variation scenario even causes consumers to sustain slightly higher $D$ values than the no variation case (panel b, compare solid and dashed lines).

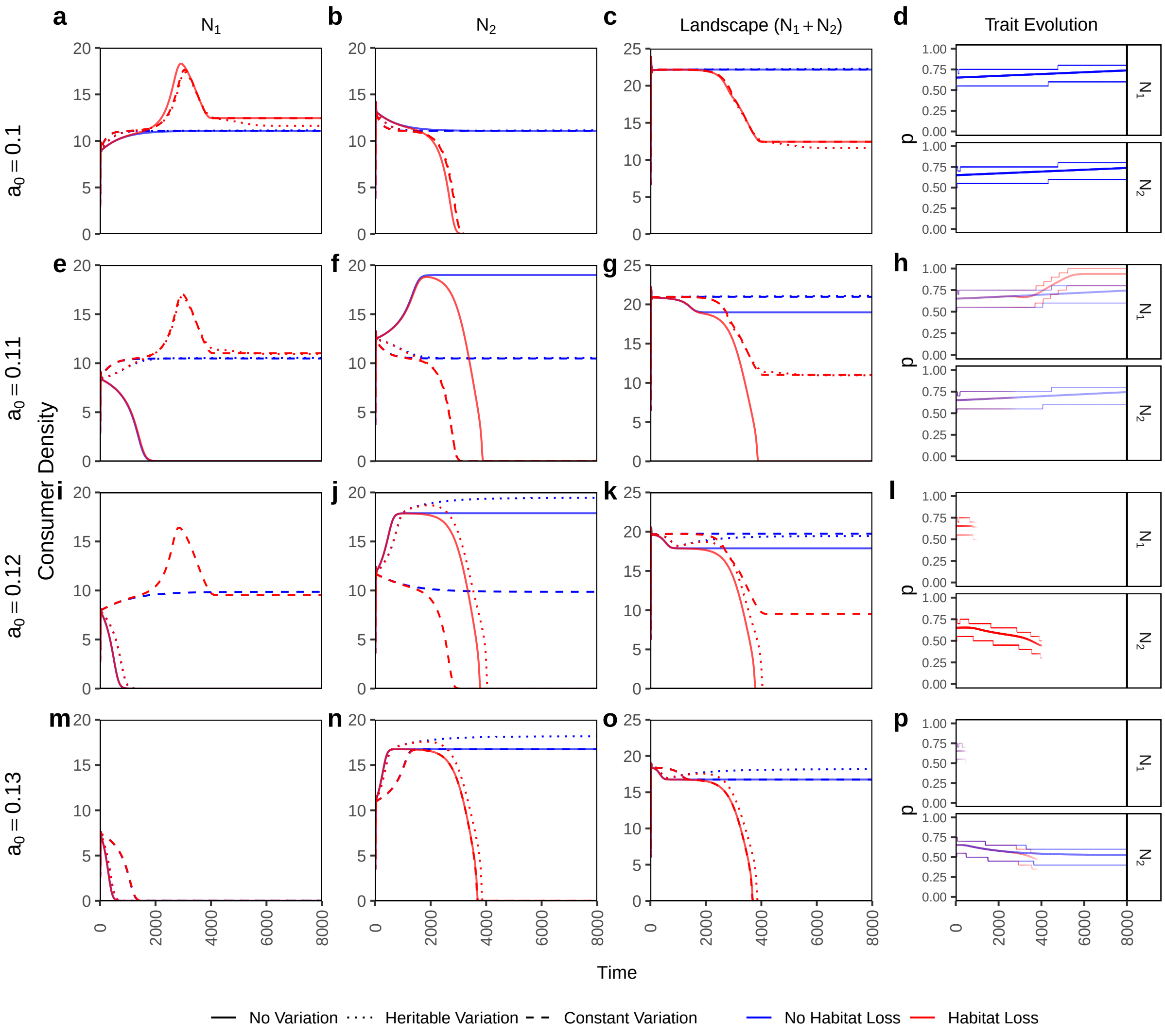


Figure S12: Time series plots of a scenario with coupled within-patch foraging preference $p$ and within-patch mating preference $\beta$ such that $p = \beta$. Patch and landscape level consumer densities are shown for the three variation scenarios in presence and absence of habitat loss. The last column shows the trait evolution of $p = \beta$ where the central line denotes the mean trait value and the thinner outer lines denote the range which contains 90% of the density. Line transparency in the trait evolution plots denote consumer density. $p$ and $\beta$ both are evolving together with initial mean $TT_{mean} = 0.65$, and standard deviation $TT_{sd} = 0.1$ with maximum possible trait value $TT_{max} = 1$, and minimum $TT_{min} = 0$. Default initial consumer densities i.e. $N_{0,1} = 3$ and $N_{0,2} = 4$. All other parameter values are default values from table 1 except the $a_0$ values that are denoted for each row.
