## Supplementary material for "Multi-scale Effects of Habitat Loss and the Role of Trait Variation": R code for generating data and figures: Bagpack-noPD.R

```
# This file is for when the patches are independent
# File storing the important functions for numerically solving the equations
# and creating some raw plots (for quick finding, these plots are not used in the manuscript)

library(deSolve) #Load the library deSolve (solves ODEs numerically)
library(ggplot2) #Load the library ggplot (for plotting)
library(ggpubr) #for ggarrange (for plotting)
library(matrixStats) #for row sds 
source("MCfun_multcases.R") # Loading the functions related to running all HL and variation combinations in one go

#defining the function for numerical ode solving (consistent to deSolve's requirements)
MC_TT_mut <- function(times, s, p) { #times, states, parameters
  with(as.list(c(s, p)), {
    len <- length(p[,1]) #number of bins
    N1 <- sum(s[1:len]); N2 <- sum(s[(1:len)+len]) #consumer densities
    R1 <- s[(2*len)+1]; R2 <- s[(2*len)+2] #resource densities
    f1 <- p[,"a0"]*s[1:len] #deaths N1
    f2 <- p[,"a0"]*s[(1:len)+len] #deaths N2
    g1   <- ((p[,"b0"]*p[,"p1"]*R1)/(p[,"b1"]+(p[,"p1"]*R1)))*(s[1:len]/2) #N1 own patch foraging
    g1cf <- 0 #N1 cross patch foraging
    g2   <- ((p[,"b0"]*p[,"p1"]*R2)/(p[,"b1"]+(p[,"p1"]*R2)))*(s[(1:len)+len]/2) #N2 own patch foraging
    g2cf <- 0 #N2 cross patch foraging
    h1 <- (p[1,"C01"]*R1)*(1 - (R1/p[1,"C11"])) #R1 resource growth 
    if (p[1,"HL"]==F){ #R2 resource growth without HL
      h2 <- (p[1,"C02"]*R2)*(1 - (R2/p[1,"C12"]))
    } else if (p[1,"HL"]==T & p[1,"HLEQ"]==T) { #R2 resource growth when HL after 10000 time steps
      if (times < 10000) {
        h2 <- (p[1,"C02"]*R2)*(1 - (R2/p[1,"C12"]))
      } else { 
        h2 <- (p[1,"C02"]*R2*(1 - (R2/p[1,"C12"]))) - ((p[1,"d"]/(1+exp(-p[1,"k"]*(times-p[1,"t0"]))))*R2)
      }
    } else if ((p[1,"HL"]==T & p[1,"HLEQ"]==F)) { #R2 resource growth when HL
      h2 <- (p[1,"C02"]*R2*(1 - (R2/p[1,"C12"]))) - ((p[1,"d"]/(1+exp(-p[1,"k"]*(times-p[1,"t0"]))))*R2)
    }
    
    # mating, inheritance, and offspring distribution part (genetic mixing (GM))
    if (p[1,"GM"] == TRUE) { #when GM is allowed
      dN <- GMFun(p, s, N1, N2, g1, g1cf, g2, g2cf, len) #See GMFun function below
      premut.dN1 <- dN[[1]]; premut.dN2 <- dN[[2]]
    } else if (p[1,"GM"] == FALSE) { #when GM is not allowed
      if (p[1,"Allee"]){ #when Allee effect is allowed
        premut.dN1 <- p[,"e"]*(1 - exp(-p[1,"theta"]*(p[,"beta"]*N1)/2))*(g1+(p[,"ec"]*g1cf))
        premut.dN2 <- p[,"e"]*(1 - exp(-p[1,"theta"]*(p[,"beta"]*N2)/2))*(g2+(p[,"ec"]*g2cf))
      } else { #when Allee effect is not allowed
        premut.dN1 <- p[,"e"]*(g1+(p[,"ec"]*g1cf))
        premut.dN2 <- p[,"e"]*(g2+(p[,"ec"]*g2cf))
      }
    }
    
    #Mutations part
    mut_away <- p[,"mu"] #creating mutation rate vector
    mut_away[c(1,len)] <- p[1,"mu"]/2 # first and last element has mu/2 mutation rate
    if (p[1,"mut"]==TRUE & len>1) { #when mutations are allowed and there is trait variation i.e. more than 1 bin
      #Mutation only affects to newborn (i.e. only the growth increment is redistributed)
      # The deaths are added after mutation
      dN1 <- premut.dN1 - (mut_away*premut.dN1) + (p[1,"mu"]*c(premut.dN1[2:len],0)/2) + (p[1,"mu"]*c(0,premut.dN1[1:(len-1)])/2) - f1
      dN2 <- premut.dN2 - (mut_away*premut.dN2) + (p[1,"mu"]*c(premut.dN2[2:len],0)/2) + (p[1,"mu"]*c(0,premut.dN2[1:(len-1)])/2) - f2
    } else { #when mutations are not allowed or there is no trait variation
      dN1 <- premut.dN1 - f1
      dN2 <- premut.dN2 - f2
    }
    dR1 <- h1 - sum(g1) - sum(g2cf)
    dR2 <- h2 - sum(g2) - sum(g1cf)
    
    #when variance in maintained
    if (p[1,"VM"]==TRUE) {
      # Maintaining constant variation. (Redistribute the surviving density into bins according to mean and sd)
      s.wt <- numeric(len)
      if ((p[1,"ttmax"]-p[1,"ttmean"]) <= (p[1,"ttmean"]-p[1,"ttmin"])) {
        s.wt[seq(p[1,"ttmin"],p[1,"ttmax"],length.out=len)>=(p[1,"ttmean"]-(p[1,"ttmax"]-p[1,"ttmean"]))] <- 
          dnorm(seq(p[1,"ttmin"],p[1,"ttmax"],length.out=len),p[1,"ttmean"],p[1,"ttsd"])[seq(p[1,"ttmin"],p[1,"ttmax"],length.out=len)>=(p[1,"ttmean"]-(p[1,"ttmax"]-p[1,"ttmean"]))]
        s.wt <- s.wt/sum(s.wt)
      } else {
        s.wt[seq(p[1,"ttmin"],p[1,"ttmax"],length.out=len)<=(p[1,"ttmean"]+(p[1,"ttmean"]-p[1,"ttmin"]))] <- 
          dnorm(seq(p[1,"ttmin"],p[1,"ttmax"],length.out=len),p[1,"ttmean"],p[1,"ttsd"])[seq(p[1,"ttmin"],p[1,"ttmax"],length.out=len)<=(p[1,"ttmean"]+(p[1,"ttmean"]-p[1,"ttmin"]))]
        s.wt <- s.wt/sum(s.wt)
      }
      dN1 <- s.wt*sum(dN1)
      dN2 <- s.wt*sum(dN2)
    }
    
    return(list(c(dN1, dN2, dR1, dR2)))
  })
}

#Traits trade-off with an underlying trait (UT) (use only linear i.e. strength=1 since non linear tradeoffs create interpretation issues)
traitrelations_UT <- function(xmin, xmax, ymin, ymax, strength=1, bins, positive=TRUE, typeII=F){
  ut <- seq(0,1,length.out = bins) # underlying trait
  if (typeII==F) {
    cd <- strength
  } else if (typeII==T) {
    cd <- 1/strength
  }
  cd1 <- sqrt(cd)
  cd2 <- cd1/cd
  if (positive == TRUE) {
    x <- xmin + (xmax-xmin)*(ut^cd1)
    y <- ymin + (ymax-ymin)*(ut^cd2)
  } else if (positive == FALSE) {
    x <- xmin + (xmax-xmin)*(ut^cd1)
    y <- ymax - (ymax-ymin)*(ut^cd2)
  }
  return(list(x=x,y=y,strength=cd))
}

# Function that creates a matrices used in GM calculations (Mb matrices in the manuscript)
GMmats <- function(len) {
  m1 <- matrix(1, nrow = len, ncol = len)
  m1b2 <- matrix(1/2, nrow = len, ncol = len)
  m.outadd <- outer(1:len,1:len,"+")
  outlist <- lapply(1:len, matrix, data= NA, nrow=len, ncol=len)
  for (i in 1:len) {
    filter_1 <- (m.outadd/2 == i)
    filter_1b2 <- (m.outadd/2 < (i+1) & m.outadd/2 > (i-1) & m.outadd/2 != i)
    outlist[[i]] <- m1*filter_1 + m1b2*filter_1b2
  }
  return(outlist)
}

#Genetic mixing: A function that facilitates mating, inheritance and offspring distribution
#Newborns with cross patch mating are split equally in own and opposite patch
GMFun <- function (p, s, N1, N2, g1, g1cf, g2, g2cf, len) {
  #Growth increment that we are going to re-distribute
  if (p[1,"Allee"]){ #with Allee effect
    dN1_temp <- p[,"e"]*(1 - exp(-p[1,"theta"]*(p[,"beta"]*N1)/2))*(g1+(p[,"ec"]*g1cf))
    dN2_temp <- p[,"e"]*(1 - exp(-p[1,"theta"]*(p[,"beta"]*N2)/2))*(g2+(p[,"ec"]*g2cf))
  } else { #without Allee effect
    dN1_temp <- p[,"e"]*(g1+(p[,"ec"]*g1cf))
    dN2_temp <- p[,"e"]*(g2+(p[,"ec"]*g2cf))
  }
  
  #Portion distributed within patch
  dN1_intra <- dN1_temp
  dN2_intra <- dN2_temp
  
  #Portion distributed across patch
  #dN1_inter <- dN1_temp - dN1_intra
  #dN2_inter <- dN2_temp - dN2_intra
  
  #Weight of each bin i.e. paternal weights
  N1_wts <- s[1:len]/sum(s[1:len]); N2_wts <- s[(1:len)+len]/sum(s[(1:len)+len])
  
  #Distribution matrices, within or between patch, correspond to A^{mn} in the manuscript
  DistMat.intra1 <- outer(dN1_intra,N1_wts,"*"); DistMat.intra2 <- outer(dN2_intra,N2_wts,"*")
  #DistMat.inter1 <- outer(dN1_inter,N2_wts,"*"); DistMat.inter2 <- outer(dN2_inter,N1_wts,"*")
  
  #initializing final growth increment vector in this time step
  dN1_SG <- numeric(length = len); dN2_SG <- numeric(length = len)
  
  for (i in 1:len) {
    #Inter-patch mating with progeny split equally in own and other patch
    dN1_SG[i] <- sum(DistMat.intra1*GMmatrix[[i]]) #+ 0.5*sum(DistMat.inter1*GMmatrix[[i]]) + 0.5*sum(DistMat.inter2*GMmatrix[[i]])
    dN2_SG[i] <- sum(DistMat.intra2*GMmatrix[[i]]) #+ 0.5*sum(DistMat.inter2*GMmatrix[[i]]) + 0.5*sum(DistMat.inter1*GMmatrix[[i]])
  }
  
  return(list(dN1_SG, dN2_SG))
}

############### functions for some raw plots #################
#Function to plot the raw time series plots (not used for manuscript)
timseriesplot <- function(foldername,savedData=FALSE,plotsave=FALSE){
  if (savedData==TRUE) {
    load(paste0(foldername,"/rawdata/file.RData"))
  }
  out <- obj$out
  len <- length(obj$params[,1])
  if (plotsave == F) {
    matplot(out[ ,1], cbind(rowSums(out[, 2:(len+1), drop=F]),
                            rowSums(out[,(len+2):(2*len+1), drop=F])), type = "l",
            xlab = "time", ylab = "Density", main = "Density time series", lwd = 2,
            xaxs="i", yaxs="i")
    legend("topright", c("N1", "N2"), col = 1:2, lty = 1:2)
    matplot(out[, 1], out[, (2*len+2):(2*len+3)], type = "l", xlab = "time", ylab = "Resource",
            main = "Respource time series", lwd = 2, xaxs="i", yaxs="i")
    legend("topright", c("R1", "R2"), col = 1:2, lty = 1:2)
  } else if (plotsave == T) {
    if (dir.exists(paste0(foldername,"/plots"))==F) {
      dir.create(paste0(foldername,"/plots"))
    }
    png(filename = paste0(foldername,"/plots/population_timeseries.png"), width = 640, height = 480)
    matplot(out[ ,1], cbind(rowSums(out[, 2:(len+1), drop=F]),
                            rowSums(out[,(len+2):(2*len+1), drop=F])), type = "l",
            xlab = "time", ylab = "Density", main = "Density time series", lwd = 2,
            xaxs="i", yaxs="i")
    legend("topright", c("N1", "N2"), col = 1:2, lty = 1:2)
    dev.off()
    png(filename = paste0(foldername,"/plots/resource_timeseries.png"), width = 640, height = 480)
    matplot(out[, 1], out[, (2*len+2):(2*len+3)], type = "l", xlab = "time", ylab = "Resource",
            main = "Respource time series", lwd = 2, xaxs="i", yaxs="i")
    legend("topright", c("R1", "R2"), col = 1:2, lty = 1:2)
    dev.off()
  }
}

#Function to plot trait distribution (not used for manuscript)
traitdistplot <- function(foldername, traits, savedData=FALSE, plotsave=FALSE){
  if (savedData==TRUE) {
    load(paste0(foldername,"/rawdata/file.RData"))
  }
  out <- obj$out
  p <- obj$params
  len <- length(p[,1])
  bardata <- rbind(N1 = out[nrow(out),2:(len+1)], N2 = out[nrow(out),(len+2):(2*len+1)])
  for (i in traits){
    if (plotsave == F) {
      barplot(bardata, beside = T, names = as.character(round(p[,i],2)), xlab = paste(i,"values"), ylab = "Density", legend.text = T, col=c("black", "red"), border = F)
    } else if (plotsave == T) {
      if (dir.exists(paste0(foldername,"/traitdist"))==F) {
        dir.create(paste0(foldername,"/traitdist"))
      }
      png(filename = paste0(foldername,"/traitdist/",i,"_traitdistribution.png"), width = 640, height = 480)
      barplot(bardata, beside = T, names = as.character(round(p[,i],2)), xlab = paste(i,"values"), ylab = "Density", legend.text = T, col=c("black", "red"), border = F)
      dev.off()
    }
  }
}

#Function to plot changes in trait distributions over time
#i.e. Heatmap for the trait values at each time step (not used for manuscript)
traitsovertime <- function(foldername, traits, times, scaled = TRUE, savedData=FALSE, plotsave=FALSE) {
  if (savedData==TRUE) {
    load(paste0(foldername,"/rawdata/file.RData"))
  }
  out <- obj$out
  p <- obj$params
  len <- length(p[,1])
  outdf <- expand.grid(tm.i=1:length(times),zvals.i=1:len)
  for (i in traits){
    if (scaled==T) {
      outdf1 <- outdf
      outdf1$patch <- "N1"
      outdf1$freq <- as.numeric(paste0(t(scale(t(out[,2:(len+1)]),center=F,scale=colSums(t(out[,2:(len+1)]))))))
      #outdf1$freq <- as.numeric(t(out[,2:(len+1)]))
      outdf2 <- outdf
      outdf2$patch <- "N2"
      outdf2$freq <- as.numeric(paste0(t(scale(t(out[,(len+2):(2*len+1)]),center=F,scale=colSums(t(out[,(len+2):(2*len+1)]))))))
      #outdf2$freq <- as.numeric(t(out[,(len+2):(2*len+1)]))
      totout <- rbind(outdf1,outdf2)  
      totout$z <- as.numeric(p[,i])[totout$zvals.i]
      totout$z <- round(totout$z,2)
      totout$tm <- as.numeric(times)[totout$tm.i]
      # generate plotting object
      p1 <- ggplot(totout,aes(x=tm,y=z,fill=freq)) + geom_tile(stat="identity") +
        facet_grid(.~patch) + scale_fill_gradient(low="white",high="black") + 
        xlab("Time") + ylab(paste("Trait",i)) + labs(fill = "Scaled density") +
        theme_bw() + theme(panel.grid.major = element_blank(),
                           panel.grid.minor = element_blank(),
                           strip.background = element_rect(colour="black", fill="white"),
                           panel.spacing.x = unit(c(0.7), "lines"),
                           axis.text.x = element_text(angle=90, vjust=0.5)) +
        scale_x_continuous(expand = c(0,0), guide = guide_axis(check.overlap = T)) + 
        scale_y_continuous(expand = c(0,0))
      #print(p1)
    } else if (scaled==F){
      outdf1 <- outdf
      outdf1$patch <- "N1"
      #outdf1$freq <- as.numeric(paste0(t(scale(t(out[,2:(len+1)]),center=F,scale=colSums(t(out[,2:(len+1)]))))))
      outdf1$freq <- as.numeric((out[,2:(len+1)]))
      outdf2 <- outdf
      outdf2$patch <- "N2"
      #outdf2$freq <- as.numeric(paste0(t(scale(t(out[,(len+2):(2*len+1)]),center=F,scale=colSums(t(out[,(len+2):(2*len+1)]))))))
      outdf2$freq <- as.numeric((out[,(len+2):(2*len+1)]))
      totout <- rbind(outdf1,outdf2)  
      totout$z <- as.numeric(p[,i])[totout$zvals.i]
      totout$z <- round(totout$z,2)
      totout$tm <- as.numeric(times)[totout$tm.i]
      # generate plotting object
      p1 <- ggplot(totout,aes(x=tm,y=z,fill=freq)) + geom_tile(stat="identity") +
        facet_grid(.~patch) + scale_fill_gradient(low="white",high="black") + 
        xlab("Time") + ylab(paste("Trait",i)) + labs(fill = "Absolute density") +
        theme_bw() + theme(panel.grid.major = element_blank(),
                           panel.grid.minor = element_blank(),
                           strip.background = element_rect(colour="black", fill="white"),
                           panel.spacing.x = unit(c(0.7), "lines"),
                           axis.text.x = element_text(angle=90, vjust=0.5)) +
        scale_x_continuous(expand = c(0,0), guide = guide_axis(check.overlap = T)) + 
        scale_y_continuous(expand = c(0,0))
      #print(p1)
    } 
    if (plotsave == F) {
      print(p1)
    } else if (plotsave == T) {
      if (dir.exists(paste0(foldername,"/traitsovertime"))==F) {
        dir.create(paste0(foldername,"/traitsovertime"))
      }
      if (scaled == F) {
        png(filename = paste0(foldername,"/traitsovertime/",i,"_abs_traitdistribution_overtime.png"), width = 640, height = 480)
        print(p1); dev.off()
      } else if (scaled == T) {
        png(filename = paste0(foldername,"/traitsovertime/",i,"_scaled_traitdistribution.png"), width = 640, height = 480)
        print(p1); dev.off() 
      }
    }
  }
}

#Plotting the traits tradeoff values (not used for manuscript)
traitrelplot <- function(foldername, traitrel, traits, plotsave=F) {
  if (plotsave == F) {
    plot(traitrel[["x"]], traitrel[["y"]], xlab = traits[1], ylab = traits[2], main = "Tradeoff")
  } else if (plotsave == T) {
    if (dir.exists(paste0(foldername,"/tradeoff"))==F) {
      dir.create(paste0(foldername,"/tradeoff"))
    }
    png(filename = paste0(foldername,"/tradeoff/",traits[2],"Vs",traits[1], "_tradeoff_strength_",
                          traitrel[["strength"]],".png"), width = 640, height = 480)
    plot(traitrel[["x"]], traitrel[["y"]], xlab = traits[1], ylab = traits[2], main = "Tradeoff")
    dev.off()
  }
}
```
