## Supplementary material for "Multi-scale Effects of Habitat Loss and the Role of Trait Variation": R code for generating data and figures: Figure_4.R

```
# Code to generate data for parameter space plots over d
# patch dependence and independence data as well

# Parameter scan over d while epsilon is evolving [~19-20 mins]
rm(list = ls())
source("Bagpack.R")

foldername = "Data/e-var-mean_0p2-sd_0p08-parsweep_d"
trait = c("e")
parsweep_trait = c("d")
obj <- MCfun_multcases(a0 = 0.1, b0 = 30, b1 = 500, bins=21,
                       p1 = 0.7, beta = 0.8, vartrait = trait, ToffMinMaxPos = c(0,1,TRUE),
                       ttmin=0, ttmax=0.4, ttmean=0.2, ttsd=0.08,
                       e = 0.2, ec = 0.9, C01 = 0.5, C02 = 0.5, C11 = 50, C12 =50,
                       N01 = 3, N02 = 4, R01 = 10, R02 = 10, foldername = foldername,
                       mu = 0.3, d = seq(0,0.5,0.025), k = 0.003, t0 = 3500, theta = 2, times = seq(0, 8000, 1),
                       mutations = T, parasweep=parsweep_trait, GeneticMixing=T, Allee = T, HLatEQ = F, savedata = T)


# Parameter scan over d while epsilon is evolving and no patch dependence [~19-20 mins]
rm(list = ls())
source("Bagpack-noPD.R")

foldername = "Data/e-var-mean_0p2-sd_0p08-parsweep_d-noPD"
trait = c("e")
parsweep_trait = c("d")
obj <- MCfun_multcases(a0 = 0.1, b0 = 30, b1 = 500, bins=21,
                       p1 = 0.7, beta = 0.8, vartrait = trait, ToffMinMaxPos = c(0,1,TRUE),
                       ttmin=0, ttmax=0.4, ttmean=0.2, ttsd=0.08,
                       e = 0.2, ec = 0.9, C01 = 0.5, C02 = 0.5, C11 = 50, C12 =50,
                       N01 = 3, N02 = 4, R01 = 10, R02 = 10, foldername = foldername,
                       mu = 0.3, d = seq(0,0.5,0.025), k = 0.003, t0 = 3500, theta = 2, times = seq(0, 8000, 1),
                       mutations = T, parasweep=parsweep_trait, GeneticMixing=T, Allee = T, HLatEQ = F, savedata = T)

#Plot the figure by sourcing the file which plots it
source("plotcode/Figure_4_plotfile.R")
```
