## Supplementary material for "Multi-scale Effects of Habitat Loss and the Role of Trait Variation": R code for generating data and figures: MCfun_multcases.R

```
# This file contains the function to run all combination of HL and trait variation
# and functions to plot the respective comparison plots (plots for quick findings, plots not directly used in the manuscript)

#A function to run all combinations of HL (presence, absence) and trait variation (no, constant, heritable)
# can sweep one parameter at a time
# can contain variation in 1 or 2 (tradeoff) parameters at a time
MCfun_multcases <- function(a0,b0,b1,p1,beta,e,ec,C01,C02,C11,C12,times,N01,N02,bins,
                            ttmin, ttmax, ttmean, ttsd, ToffMinMaxPos,
                           R01,R02,foldername,mu,d,k,t0,theta,mutations=TRUE,
                           GeneticMixing=TRUE,HLatEQ=FALSE,Allee=TRUE,savedata=FALSE,
                           vartrait=c("e"),parasweep=c()){
  
  grain <- as.integer((length(times)-1)/tail(times,n=1)) #used to store data only at integer times (reduces saved .Rdata file size)
  
  #creating directories to save data
  if (savedata==TRUE & !dir.exists("Data")) {
    dir.create("Data") #Create a new folder
  }
  
  dir.create(foldername) #Create a new folder, will give error if folder already exists
  
  cases <- expand.grid(Va=c(T,F),HL=c(T,F),VM=c(T,F)) #All combinations of HL and variation 
  cases <- cases[!(cases[,"Va"]==F & cases[,"VM"]==T),] #Remove meaningless cases of variation
  
  if (length(parasweep)==0) { #when there is no parameter sweeping
    obj <- list() #initializing big output object
    for (i in 1:nrow(cases)){ #iterating through the combinations
      
      p <- cbind(sr.no=1:bins,a0=a0,b0=b0,b1=b1,p1=p1,beta=beta,N01=N01,N02=N02,R01=R01,R02=R02,
                 e=e,ec=ec,C01=C01,C02=C02,C11=C11,C12=C12,mu=mu,d=d,k=k,t0=t0,theta=theta,
                 ttmin=ttmin,ttmax=ttmax,ttmean=ttmean,ttsd=ttsd,Allee=Allee,bins=bins,
                 HL=cases[i,"HL"],Va=cases[i,"Va"],VM=cases[i,"VM"],
                 GM=GeneticMixing,HLEQ=HLatEQ,mut=mutations) #Parameters matrix for deSolve, nrow=bins
      
      p[,vartrait[1]] <- seq(ttmin,ttmax,length.out=bins) #assiging equidistant trait values to the bins
      if (cases[i,"Va"] == F & length(vartrait)==1) { 
        p <- t(as.matrix(colMeans(p))) #When there is no variation, take mean value of the range provided
        p[,vartrait[1]] <- ttmean
      }
      if (length(vartrait)==2) { #i.e. when tradeoff between the two traits
        traitrel <- traitrelations_UT(ttmin,ttmax,ToffMinMaxPos[1],ToffMinMaxPos[2],1,bins,ToffMinMaxPos[3])
        # values assigned to each bin according to tradeoff
        p[,vartrait[1]] <- traitrel$x
        p[,vartrait[2]] <- traitrel$y
        if (cases[i,"Va"] == F) {
          p <- t(as.matrix(colMeans(p))) #When there is no variation, take mean value of the range provided
          p[,vartrait[1]] <- ttmean
          p[,vartrait[2]] <- traitrel$y[which(traitrel$x>=ttmean)[1]]
        }
      } else if (length(vartrait)>=3) {stop("Cannot vary more than 2 traits for now")}
      
      #setting initial distribution
      len <- length(p[,1])
      if (len==1) {
        s.wt <- 1
      } else {
        s.wt <- numeric(len)
        if ((ttmax-ttmean) <= (ttmean-ttmin)) {
          s.wt[seq(ttmin,ttmax,length.out=len)>=(ttmean-(ttmax-ttmean))] <- 
            dnorm(seq(ttmin,ttmax,length.out=len),ttmean,ttsd)[seq(ttmin,ttmax,length.out=len)>=(ttmean-(ttmax-ttmean))]
          s.wt <- s.wt/sum(s.wt)
        } else {
          s.wt[seq(ttmin,ttmax,length.out=len)<=(ttmean+(ttmean-ttmin))] <- 
            dnorm(seq(ttmin,ttmax,length.out=len),ttmean,ttsd)[seq(ttmin,ttmax,length.out=len)<=(ttmean+(ttmean-ttmin))]
          s.wt <- s.wt/sum(s.wt)
        }
      }
      s <- c(s.wt*p[1,"N01"], s.wt*p[1,"N02"], p[1,"R01"], p[1,"R02"]) #initial distribution of consumers in the bins

      GMmatrix <<- GMmats(len) ###### GLOBAL VARIABLE ###### creating the matrices corresponding to Mb in the manuscript
      
      if (cases[i,"HL"]==T & HLatEQ==T) { #habitat loss will start after 10000 steps if HLatEQ==TRUE
        times <- c(0:9999,(times+10000))
      }
      out <- ode(s, times, MC_TT_mut, p, rtol=1e-4, atol=1e-4) #Solving the deSolve function ode(state,times,function,parameters)
      if (cases[i,"HL"]==F) { # appending the output into the big output object
        obj <- append(obj, list(list(params=p,states=s,out=out[seq(1,(length(times)),grain),]))) #big object to be saved as raw file
      } else if (cases[i,"HL"]==T & HLatEQ==T) { #For habitat loss after equilibrium, only the data after habitat loss starts is stored
        obj <- append(obj, list(list(params=p,states=s,out=out[10001:nrow(out),][seq(1,(length(times)),grain),]))) #big object to be saved as raw file
      } else if (cases[i,"HL"]==T & HLatEQ==F) {
        obj <- append(obj, list(list(params=p,states=s,out=out[seq(1,(length(times)),grain),]))) #big object to be saved as raw file
      }
    }
    
    if (savedata==TRUE) { #save the R.Data object if savedata=TRUE
      sink(paste0(foldername,"/functioncall.txt"))
      print(as.list(match.call()))
      sink()
      dir.create(paste0(foldername,"/rawdata")) #Create folder to save raw data
      save(obj,file = paste0(foldername,"/rawdata","/file.RData")) #Save raw data in inside the folder 
    }
    return(obj)
  
    
  # Sweeping the parameter range
  } else if (length(parasweep) > 0) { #Note: Write parameter name as a character in the parasweep attribute

    for (paraname in parasweep){
      if (savedata==TRUE & !dir.exists(paste0(foldername,"/",paraname))) { #create new folder
        dir.create(paste0(foldername,"/",paraname))
      }
      obj <- list() #initial big output object
      parvals <- eval(as.name(paraname)) #vector input is given to parameter which is to be evolved, here the values are extected in parvals
      for (j in parvals) { #iteration over each parameter value of the parameter that is swept
        assign(paraname,j) #assign the current parameter value in the loop to the parameter variable
        for (i in 1:nrow(cases)){ 
          p <- cbind(sr.no=1:bins,a0=a0,b0=b0,b1=b1,p1=p1,beta=beta,N01=N01,N02=N02,R01=R01,R02=R02,
                     e=e,ec=ec,C01=C01,C02=C02,C11=C11,C12=C12,mu=mu,d=d,k=k,t0=t0,theta=theta,
                     ttmin=ttmin,ttmax=ttmax,ttmean=ttmean,ttsd=ttsd,Allee=Allee,bins=bins,
                     HL=cases[i,"HL"],Va=cases[i,"Va"],VM=cases[i,"VM"],
                     GM=GeneticMixing,HLEQ=HLatEQ,mut=mutations) #Parameters matrix for deSolve, nrow=bins
          p[,vartrait[1]] <- seq(ttmin,ttmax,length.out=bins) #assiging equidistant trait values to the bins
          if (cases[i,"Va"] == F & length(vartrait)==1) { 
            p <- t(as.matrix(colMeans(p))) #When there is no variation, take mean value of the range provided
            p[,vartrait[1]] <- ttmean
          }
          if (length(vartrait)==2) { #i.e. when tradeoff between the two traits
            traitrel <- traitrelations_UT(ttmin,ttmax,ToffMinMaxPos[1],ToffMinMaxPos[2],1,bins,ToffMinMaxPos[3])
            # values assigned to each bin according to tradeoff
            p[,vartrait[1]] <- traitrel$x
            p[,vartrait[2]] <- traitrel$y
            if (cases[i,"Va"] == F) {
              p <- t(as.matrix(colMeans(p))) #When there is no variation, take mean value of the range provided
              p[,vartrait[1]] <- ttmean
              p[,vartrait[2]] <- traitrel$y[which(traitrel$x>=ttmean)[1]]
            }
          } else if (length(vartrait)>=3) {stop("Cannot vary more than 2 traits for now")}
          
          #setting initial distribution
          len <- length(p[,1])
          if (len==1) {
            s.wt <- 1
          } else {
            s.wt <- numeric(len)
            if ((ttmax-ttmean) <= (ttmean-ttmin)) {
              s.wt[seq(ttmin,ttmax,length.out=len)>=(ttmean-(ttmax-ttmean))] <- 
                dnorm(seq(ttmin,ttmax,length.out=len),ttmean,ttsd)[seq(ttmin,ttmax,length.out=len)>=(ttmean-(ttmax-ttmean))]
              s.wt <- s.wt/sum(s.wt)
            } else {
              s.wt[seq(ttmin,ttmax,length.out=len)<=(ttmean+(ttmean-ttmin))] <- 
                dnorm(seq(ttmin,ttmax,length.out=len),ttmean,ttsd)[seq(ttmin,ttmax,length.out=len)<=(ttmean+(ttmean-ttmin))]
              s.wt <- s.wt/sum(s.wt)
            }
          }
          s <- c(s.wt*p[1,"N01"], s.wt*p[1,"N02"], p[1,"R01"], p[1,"R02"]) #initial distribution of consumers in the bins
          
          GMmatrix <<- GMmats(len) ###### GLOBAL VARIABLE ###### creating the matrices corresponding to Mb in the manuscript
          
          if (cases[i,"HL"]==T & HLatEQ==T) { #habitat loss will start after 10000 steps when HLateq=TRUE
            times <- c(0:9999,(times+10000))
          }
          out <- ode(s, times, MC_TT_mut, p, rtol=1e-4, atol=1e-4) #Solving the deSolve function ode(states,times,function,parameters)
          if (cases[i,"HL"]==F) { # appending the output into the big output object
            obj[[as.name(j)]] <- append(obj[[as.name(j)]], list(list(params=p,states=s,out=out[seq(1,(length(times)),grain),]))) #big object to be saved as raw file
          } else if (cases[i,"HL"]==T & HLatEQ==T) { #For habitat loss after, only the data after habitat loss starts is stored
            obj[[as.name(j)]] <- append(obj[[as.name(j)]], list(list(params=p,states=s,out=out[10001:nrow(out),][seq(1,(length(times)),grain),]))) #big object to be saved as raw file
          } else if (cases[i,"HL"]==T & HLatEQ==F) {
            obj[[as.name(j)]] <- append(obj[[as.name(j)]], list(list(params=p,states=s,out=out[seq(1,(length(times)),grain),]))) #big object to be saved as raw file
          }
        }
        print(j)
      }
      if (savedata==TRUE) { #save the big output object when savedata=TRUE
        sink(paste0(foldername,"/",paraname,"/functioncall.txt"))
        print(as.list(match.call()))
        sink()
        dir.create(paste0(foldername,"/",paraname,"/rawdata")) #Create folder to save raw data
        save(obj,file = paste0(foldername,"/",paraname,"/rawdata","/file.RData")) #Save raw data in inside the folder 
      }
      return(obj)
    }
  }
}

#Function that plots raw plots for individual parameter combinations (plots not directly used in manuscript)
plot_multcases <- function(foldername, traits, parasweep=c()) {
  if (length(parasweep)==0) {
    load(paste0(foldername,"/rawdata/file.RData"))
    if (length(traits)==2) { #Plottig the tradeoff
      traitrelplot_multcases(foldername, traits, obj)
    }
    HLshape_multcases(foldername, obj) #Plot shape of the habitat loss (resource reduction term with time)
    cases <- expand.grid(Va=c(T,F),HL=c(T,F),VM=c(T,F))
    cases <- cases[!(cases[,"Va"]==F & cases[,"VM"]==T),]
    for (i in 1:nrow(cases)) {
      if (dir.exists(paste0(foldername,"/","Va_",cases[i,"Va"],"_HL_",cases[i,"HL"],"_VM_",cases[i,"VM"]))==F) {
        dir.create(paste0(foldername,"/","Va_",cases[i,"Va"],"_HL_",cases[i,"HL"],"_VM_",cases[i,"VM"])) 
      }
      foldernameTS<-paste0(foldername,"/","Va_",cases[i,"Va"],"_HL_",cases[i,"HL"],"_VM_",cases[i,"VM"])
      timeseriesplot_multcases(foldernameTS,i,obj)
      traitsovertime_multcases(foldernameTS,i,traits,obj)
    }
  }
  else if (length(parasweep)==1) {
    load(paste0(foldername,"/",parasweep[1],"/rawdata/file.RData"))
    cases <- expand.grid(Va=c(T,F),HL=c(T,F),VM=c(T,F))
    cases <- cases[!(cases[,"Va"]==F & cases[,"VM"]==T),]
    
    for (j in names(obj)) {
      foldername.para <- paste0(foldername,"/",parasweep[1],"/",parasweep[1],"_value_",j)
      if (!dir.exists(foldername.para)) {
        dir.create(foldername.para)
      }
      obj1 <- obj[[j]]
      if (length(traits)==2) { #Plottig the tradeoff
        traitrelplot_multcases(foldername.para, traits, obj1)
      }
      HLshape_multcases(foldername.para, obj1) #Plot shape of the habitat loss (resource reduction term with time)
      
      for (i in 1:nrow(cases)) {
        foldernameTS<-paste0(foldername.para,"/","Va_",cases[i,"Va"],"_HL_",cases[i,"HL"],"_VM_",cases[i,"VM"])
        if (!dir.exists(foldernameTS)) {
          dir.create(foldernameTS)
        }
        times <- obj1[[i]]$out[,1]
        
        timeseriesplot_multcases(foldernameTS,i,obj1)
        traitsovertime_multcases(foldernameTS,i,traits,obj1)
      }
    }
  }
  else {
    stop("Currently this works when only one parameter is varied")
  }
}

#function to plot variance comparison cases (raw plots, not directly used in manuscript)
plot_varcomp <- function(foldername,parasweep=c()) {
  if (length(parasweep)==0) {
    load(paste0(foldername,"/rawdata/file.RData"))
    if (!dir.exists(paste0(foldername,"/plot_varcomp"))){
      dir.create(paste0(foldername,"/plot_varcomp"))
    }
    cases <- expand.grid(Va=c(T,F),HL=c(T,F),VM=c(T,F))
    cases <- cases[!(cases[,"Va"]==F & cases[,"VM"]==T),]
    
    # Habitat loss case
    HLData <- data.frame()
    for (i in c(1:nrow(cases))[cases[,"HL"]]) { #comparison in presence of HL
      len <- nrow(obj[[i]]$params)
      tempData <- data.frame(cbind(t=obj[[i]]$out[,1], N1=rowSums(obj[[i]]$out[, 2:(len+1), drop=F]),
                                   N2=rowSums(obj[[i]]$out[,(len+2):(2*len+1), drop=F]),
                                   R1=obj[[i]]$out[, (2*len+2)],R2=obj[[i]]$out[, (2*len+3)]))
      tempData$type <- if (cases[i,"VM"]==TRUE) "Constant Variation" else if (cases[i,"Va"]==FALSE) "No Variation" else "Heritable Variation"
      HLData <- rbind(HLData,tempData)
    }
    HLData$type <- factor(HLData$type,levels=c("No Variation","Heritable Variation","Constant Variation"))
    #Basic ggplot object
    p <- ggplot(data=HLData,aes(x=t)) + xlab("Time") + 
      scale_color_manual(values = c("No Variation"="#56B4E9", "Heritable Variation"="#F0E442",
                                    "Constant Variation"="#D55E00")) +
      scale_linetype_manual(values = c("1"="solid","2"="dotted")) +
      theme_bw() + theme(plot.title = element_text(hjust = 0.5, size = 20),
                         axis.title.x = element_text(size = 15),
                         axis.title.y = element_text(size = 15),
                         axis.text = element_text(size = 10, face = "bold"),
                         legend.title = element_text(size = 15),
                         legend.text = element_text(size = 10),
                         strip.text = element_text(size = 12),
                         panel.grid.major = element_blank(),
                         panel.grid.minor = element_blank(),
                         strip.background = element_rect(colour="black", fill="white"),
                         panel.border = element_rect(colour = "black"),
                         panel.spacing = unit(0, "lines")) +
      scale_x_continuous(expand = c(0,0)) + scale_y_continuous(expand = c(0,0))
    
    pNHL <- p + geom_line(aes(y=N1, color=type, linetype="1"), linewidth=1.2) +
      geom_line(aes(y=N2, color=type, linetype="2"), linewidth=1.2) + 
      ylab("Population Density") + labs(color="Trait Variation",linetype="patch") +
      ggtitle(paste("Population Density Time-Series","HL"))
    png(filename = paste0(foldername,"/plot_varcomp/population_HL.png"), width = 852, height = 480)
    print(pNHL); dev.off()
    
    pRHL <- p + geom_line(aes(y=R1, color=type, linetype="1"), linewidth=1.2) +
      geom_line(aes(y=R2, color=type, linetype="2"), linewidth=1.2) + 
      ylab("Resource Density") + labs(color="Trait Variation",linetype="patch") +
      ggtitle(paste("Resource Density Time-Series", "HL"))
    png(filename = paste0(foldername,"/plot_varcomp/resource_HL.png"), width = 852, height = 480)
    print(pRHL); dev.off()
    
    # No Habitat loss case
    Data <- data.frame()
    for (i in c(1:nrow(cases))[!cases[,"HL"]]) { #comparison in absence of HL
      len <- nrow(obj[[i]]$params)
      tempData <- data.frame(cbind(t=obj[[i]]$out[,1], N1=rowSums(obj[[i]]$out[, 2:(len+1), drop=F]),
                                   N2=rowSums(obj[[i]]$out[,(len+2):(2*len+1), drop=F]),
                                   R1=obj[[i]]$out[, (2*len+2)],R2=obj[[i]]$out[, (2*len+3)]))
      tempData$type <- if (cases[i,"VM"]==TRUE) "Constant Variation" else if (cases[i,"Va"]==FALSE) "No Variation" else "Heritable Variation"
      Data <- rbind(Data,tempData)
    }
    Data$type <- factor(Data$type,levels=c("No Variation","Heritable Variation","Constant Variation"))
    #Basic ggplot object
    p <- ggplot(data=Data,aes(x=t)) + xlab("Time") + 
      scale_color_manual(values = c("No Variation"="#56B4E9", "Heritable Variation"="#F0E442",
                                    "Constant Variation"="#D55E00")) +
      scale_linetype_manual(values = c("1"="solid","2"="dotted")) +
      theme_bw() + theme(plot.title = element_text(hjust = 0.5, size = 20),
                         axis.title.x = element_text(size = 15),
                         axis.title.y = element_text(size = 15),
                         axis.text = element_text(size = 10, face = "bold"),
                         legend.title = element_text(size = 15),
                         legend.text = element_text(size = 10),
                         strip.text = element_text(size = 12),
                         panel.grid.major = element_blank(),
                         panel.grid.minor = element_blank(),
                         strip.background = element_rect(colour="black", fill="white"),
                         panel.border = element_rect(colour = "black"),
                         panel.spacing = unit(0, "lines")) +
      scale_x_continuous(expand = c(0,0)) + scale_y_continuous(expand = c(0,0))
    
    pN <- p + geom_line(aes(y=N1, color=type, linetype="1"), linewidth=1.2) +
      geom_line(aes(y=N2, color=type, linetype="2"), linewidth=1.2) + 
      ylab("Population Density") + labs(color="Trait Variation",linetype="patch") +
      ggtitle("Population Density Time-Series")
    png(filename = paste0(foldername,"/plot_varcomp/population.png"), width = 852, height = 480)
    print(pN); dev.off()
    
    pR <- p + geom_line(aes(y=R1, color=type, linetype="1"), linewidth=1.2) +
      geom_line(aes(y=R2, color=type, linetype="2"), linewidth=1.2) + 
      ylab("Resource Density") + labs(color="Trait Variation",linetype="patch") +
      ggtitle("Resource Density Time-Series")
    png(filename = paste0(foldername,"/plot_varcomp/resource.png"), width = 852, height = 480)
    print(pR); dev.off()
  } 
  else if (length(parasweep==1)) {
    load(paste0(foldername,"/",parasweep[1],"/rawdata/file.RData"))
    if (!dir.exists(paste0(foldername,"/",parasweep[1],"/plot_varcomp"))) {
      dir.create(paste0(foldername,"/",parasweep[1],"/plot_varcomp"))
    }
    cases <- expand.grid(Va=c(T,F),HL=c(T,F),VM=c(T,F))
    cases <- cases[!(cases[,"Va"]==F & cases[,"VM"]==T),]
    
    Data <- data.frame()
    for (j in 1:length(obj)) {
      for (i in 1:nrow(cases)) {
        obj1<-obj[[j]]
        len <- nrow(obj1[[i]]$params)
        tempData <- data.frame(cbind(t=obj1[[i]]$out[,1], N1=rowSums(obj1[[i]]$out[, 2:(len+1), drop=F]),
                                     N2=rowSums(obj1[[i]]$out[,(len+2):(2*len+1), drop=F]),
                                     R1=obj1[[i]]$out[, (2*len+2)],R2=obj1[[i]]$out[, (2*len+3)]))
        tempData$type <- if (cases[i,"VM"]==TRUE) "Constant Variation" else if (cases[i,"Va"]==FALSE) "No Variation" else "Heritable Variation"
          #paste0("Va_",cases[i,"Va"],"_VM_",cases[i,"VM"])
        tempData$HL <- if (cases[i,"HL"]==TRUE) "HL patch 2" else "No HL"
        tempData$param_val <- names(obj)[j]
        Data <- rbind(Data,tempData)
      }
    }
    Data$type <- factor(Data$type,levels=c("No Variation","Heritable Variation","Constant Variation"))
    Data$param_val <- factor(Data$param_val,levels=c(names(obj)))
    #Habitat loss case
    #Basic ggplot object
    p <- ggplot(data=subset(Data,HL=="HL patch 2"),aes(x=t)) + xlab("Time") + 
      facet_grid(.~type) +
      scale_linetype_manual(values = c("1"="solid","2"="dotted")) +
      theme_bw() + theme(plot.title = element_text(hjust = 0.5, size = 20),
                         axis.title.x = element_text(size = 15),
                         axis.title.y = element_text(size = 15),
                         axis.text = element_text(size = 10, face = "bold"),
                         legend.title = element_text(size = 15),
                         legend.text = element_text(size = 10),
                         legend.position="bottom",
                         strip.text = element_text(size = 12),
                         panel.grid.major = element_blank(),
                         panel.grid.minor = element_blank(),
                         strip.background = element_rect(colour="black", fill="white"),
                         panel.border = element_rect(colour = "black"),
                         panel.spacing = unit(1, "lines")) +
      scale_x_continuous(expand = c(0,0),guide = guide_axis(check.overlap = TRUE)) + 
      scale_y_continuous(expand = c(0,0))
    
    pNHL <- p + geom_line(aes(y=N1, color=param_val, linetype="1"), linewidth=1.2) +
      geom_line(aes(y=N2, color=param_val, linetype="2"), linewidth=1.2) + 
      ylab("Population Density") + labs(color=paste0(parasweep[1],"_value"),linetype="patch") +
      ggtitle(paste("Changing parameter",parasweep[1],"HL"))
    png(filename = paste0(foldername,"/",parasweep[1],"/plot_varcomp/population_HL.png"), width = 1280, height = 720)
    print(pNHL); dev.off()
    
    pRHL <- p + geom_line(aes(y=R1, color=param_val, linetype="1"), linewidth=1.2) +
      geom_line(aes(y=R2, color=param_val, linetype="2"), linewidth=1.2) + 
      ylab("Resource Density") + labs(color=paste0(parasweep[1],"_value"),linetype="patch") +
      ggtitle(paste("Changing parameter",parasweep[1], "HL"))
    png(filename = paste0(foldername,"/",parasweep[1],"/plot_varcomp/resource_HL.png"), width = 1280, height = 720)
    print(pRHL); dev.off()
    
    #N0 Habitat loss case
    #Basic ggplot object
    p <- ggplot(data=subset(Data,HL=="No HL"),aes(x=t)) + xlab("Time") + 
      facet_grid(.~type) + 
      scale_linetype_manual(values = c("1"="solid","2"="dotted")) +
      theme_bw() + theme(plot.title = element_text(hjust = 0.5, size = 20),
                         axis.title.x = element_text(size = 15),
                         axis.title.y = element_text(size = 15),
                         axis.text = element_text(size = 10, face = "bold"),
                         legend.title = element_text(size = 15),
                         legend.text = element_text(size = 10),
                         legend.position="bottom",
                         strip.text = element_text(size = 12),
                         panel.grid.major = element_blank(),
                         panel.grid.minor = element_blank(),
                         strip.background = element_rect(colour="black", fill="white"),
                         panel.border = element_rect(colour = "black"),
                         panel.spacing = unit(1, "lines")) +
      scale_x_continuous(expand = c(0,0),guide = guide_axis(check.overlap = TRUE)) + 
      scale_y_continuous(expand = c(0,0))
    
    pNHL <- p + geom_line(aes(y=N1, color=param_val, linetype="1"), linewidth=1.2) +
      geom_line(aes(y=N2, color=param_val, linetype="2"), linewidth=1.2) + 
      ylab("Population Density") + labs(color=paste0(parasweep[1],"_value"),linetype="patch") +
      ggtitle(paste("Changing parameter",parasweep[1],"No HL"))
    png(filename = paste0(foldername,"/",parasweep[1],"/plot_varcomp/population_NoHL.png"), width = 1280, height = 720)
    print(pNHL); dev.off()
    
    pRHL <- p + geom_line(aes(y=R1, color=param_val, linetype="1"), linewidth=1.2) +
      geom_line(aes(y=R2, color=param_val, linetype="2"), linewidth=1.2) + 
      ylab("Resource Density") + labs(color=paste0(parasweep[1],"_value"),linetype="patch") +
      ggtitle(paste("Changing parameter",parasweep[1], "No HL"))
    png(filename = paste0(foldername,"/",parasweep[1],"/plot_varcomp/resource_NoHL.png"), width = 1280, height = 720)
    print(pRHL); dev.off()
  } 
  else {
    stop("Currently this works when only one parameter is varied")
  }
}

#Function to plot HL comparison cases (raw plots, not directly used in manuscript)
plot_HLcomp <- function(foldername,parasweep=c()) {
  if (length(parasweep)==0) {
    load(paste0(foldername,"/rawdata/file.RData"))
    if (!dir.exists(paste0(foldername,"/plot_HLcomp"))){
      dir.create(paste0(foldername,"/plot_HLcomp"))
    }
    cases <- expand.grid(Va=c(T,F),HL=c(T,F),VM=c(T,F))
    cases <- cases[!(cases[,"Va"]==F & cases[,"VM"]==T),]
    # Habitat loss data
    HLData <- data.frame()
    for (i in c(1:nrow(cases))[cases[,"HL"]]) { #comparison in presence of HL
      len <- nrow(obj[[i]]$params)
      tempData <- data.frame(cbind(t=obj[[i]]$out[,1], N1=rowSums(obj[[i]]$out[, 2:(len+1), drop=F]),
                                   N2=rowSums(obj[[i]]$out[,(len+2):(2*len+1), drop=F]),
                                   R1=obj[[i]]$out[, (2*len+2)],R2=obj[[i]]$out[, (2*len+3)]))
      tempData$type <- if (cases[i,"VM"]==TRUE) "Constant Variation" else if (cases[i,"Va"]==FALSE) "No Variation" else "Heritable Variation"
      HLData <- rbind(HLData,tempData)
    }
    HLData$type <- factor(HLData$type,levels=c("No Variation","Heritable Variation","Constant Variation"))
    HLData$HL <- "HL patch 2"
    
    # No Habitat loss data
    NHLData <- data.frame()
    for (i in c(1:nrow(cases))[!cases[,"HL"]]) { #comparison in absence of HL
      len <- nrow(obj[[i]]$params)
      tempData <- data.frame(cbind(t=obj[[i]]$out[,1], N1=rowSums(obj[[i]]$out[, 2:(len+1), drop=F]),
                                   N2=rowSums(obj[[i]]$out[,(len+2):(2*len+1), drop=F]),
                                   R1=obj[[i]]$out[, (2*len+2)],R2=obj[[i]]$out[, (2*len+3)]))
      tempData$type <- if (cases[i,"VM"]==TRUE) "Constant Variation" else if (cases[i,"Va"]==FALSE) "No Variation" else "Heritable Variation"
      NHLData <- rbind(NHLData,tempData)
    }
    NHLData$type <- factor(NHLData$type,levels=c("No Variation","Heritable Variation","Constant Variation"))
    NHLData$HL <- "No HL"
    
    Data <- rbind(HLData,NHLData)
    
    #The basic ggplot object
    p <- ggplot(data=Data,aes(x=t)) + xlab("Time") + 
      scale_color_manual(values = c("No Variation"="#56B4E9", "Heritable Variation"="#F0E442",
                                    "Constant Variation"="#D55E00")) +
      scale_linetype_manual(breaks = c("No HL", "HL patch 2"),
                            values = c("No HL"="solid", "HL patch 2"="dotted")) +
      theme_bw() + theme(plot.title = element_text(hjust = 0.5, size = 20),
                         axis.title.x = element_text(size = 15),
                         axis.title.y = element_text(size = 15),
                         axis.text = element_text(size = 10, face = "bold"),
                         legend.title = element_text(size = 15),
                         legend.text = element_text(size = 10),
                         strip.text = element_text(size = 12),
                         panel.grid.major = element_blank(),
                         panel.grid.minor = element_blank(),
                         strip.background = element_rect(colour="black", fill="white"),
                         panel.border = element_rect(colour = "black"),
                         panel.spacing = unit(0, "lines")) +
      scale_x_continuous(expand = c(0,0)) + scale_y_continuous(expand = c(0,0))
    
    #Plot population density comparison for HL and NHL
    pN1 <- p + geom_line(aes(y=N1, color=type, linetype=HL), linewidth=1.2) + 
      labs(color="Trait Variation",linetype="Habitat Loss") + 
      ylab("Population Density N1") + ggtitle("Population Density HL vs NHL")
    png(filename = paste0(foldername,"/plot_HLcomp/N1_HLvsNHL.png"), width = 852, height = 480)
    print(pN1); dev.off()
    pN2 <- p + geom_line(aes(y=N2, color=type, linetype=HL), linewidth=1.2) + 
      labs(color="Trait Variation",linetype="Habitat Loss") + 
      ylab("Population Density N2") + ggtitle("Population Density HL vs NHL")
    png(filename = paste0(foldername,"/plot_HLcomp/N2_HLvsNHL.png"), width = 852, height = 480)
    print(pN2); dev.off()
    
    #Plot resource comparison for HL and NHL
    pR1 <- p + geom_line(aes(y=R1, color=type, linetype=HL), linewidth=1.2) + 
      labs(color="Trait Variation",linetype="Habitat Loss") + 
      ylab("Resource Density R1") + ggtitle("Resource Density HL vs NHL")
    png(filename = paste0(foldername,"/plot_HLcomp/R1_HLvsNHL.png"), width = 852, height = 480)
    print(pR1); dev.off()
    pR2 <- p + geom_line(aes(y=R2, color=type, linetype=HL), linewidth=1.2) + 
      labs(color="Trait Variation",linetype="Habitat Loss") + 
      ylab("Resource Density R2") + ggtitle("Resource Density HL vs NHL")
    png(filename = paste0(foldername,"/plot_HLcomp/R2_HLvsNHL.png"), width = 852, height = 480)
    print(pR2); dev.off()
  }
  else if (length(parasweep)==1) {
    load(paste0(foldername,"/",parasweep[1],"/rawdata/file.RData"))
    if (!dir.exists(paste0(foldername,"/",parasweep[1],"/plot_HLcomp"))) {
      dir.create(paste0(foldername,"/",parasweep[1],"/plot_HLcomp"))
    }
    cases <- expand.grid(Va=c(T,F),HL=c(T,F),VM=c(T,F))
    cases <- cases[!(cases[,"Va"]==F & cases[,"VM"]==T),]
    
    Data <- data.frame()
    for (j in 1:length(obj)) {
      for (i in 1:nrow(cases)) {
        obj1<-obj[[j]]
        len <- nrow(obj1[[i]]$params)
        tempData <- data.frame(cbind(t=obj1[[i]]$out[,1], N1=rowSums(obj1[[i]]$out[, 2:(len+1), drop=F]),
                                     N2=rowSums(obj1[[i]]$out[,(len+2):(2*len+1), drop=F]),
                                     R1=obj1[[i]]$out[, (2*len+2)],R2=obj1[[i]]$out[, (2*len+3)]))
        tempData$type <- if (cases[i,"VM"]==TRUE) "Constant Variation" else if (cases[i,"Va"]==FALSE) "No Variation" else "Heritable Variation"
        #paste0("Va_",cases[i,"Va"],"_VM_",cases[i,"VM"])
        tempData$HL <- if (cases[i,"HL"]==TRUE) "HL patch 2" else "No HL"
        tempData$param_val <- names(obj)[j]
        Data <- rbind(Data,tempData)
      }
    }
    Data$type <- factor(Data$type,levels=c("No Variation","Heritable Variation","Constant Variation"))
    Data$param_val <- factor(Data$param_val,levels=c(names(obj)))
    
    #The basic ggplot object
    p <- ggplot(data=Data,aes(x=t)) + xlab("Time") + 
      scale_linetype_manual(breaks = c("No HL", "HL patch 2"),
                            values = c("No HL"="solid", "HL patch 2"="dotted")) +
      ggtitle(paste("Changing parameter",parasweep[1],"HL vs NHL")) + 
      theme_bw() + theme(plot.title = element_text(hjust = 0.5, size = 20),
                         axis.title.x = element_text(size = 15),
                         axis.title.y = element_text(size = 15),
                         axis.text = element_text(size = 10, face = "bold"),
                         legend.title = element_text(size = 15),
                         legend.text = element_text(size = 10),
                         legend.position = "bottom",
                         strip.text = element_text(size = 12),
                         panel.grid.major = element_blank(),
                         panel.grid.minor = element_blank(),
                         strip.background = element_rect(colour="black", fill="white"),
                         panel.border = element_rect(colour = "black"),
                         panel.spacing = unit(1, "lines")) +
      scale_x_continuous(expand = c(0,0)) + scale_y_continuous(expand = c(0,0))
    
    #Plot population density comparison for HL and NHL
    pN1 <- p + geom_line(aes(y=N1, color=param_val, linetype=HL), linewidth=1.2) + 
      facet_grid(.~type) + labs(color=paste0(parasweep[1],"_value"),linetype="Habitat Loss") + 
      ylab("Population Density N1")
    png(filename = paste0(foldername,"/",parasweep[1],"/plot_HLcomp/N1_HLvsNHL.png"), width = 1280, height = 720)
    print(pN1); dev.off()
    pN2 <- p + geom_line(aes(y=N2, color=param_val, linetype=HL), linewidth=1.2) + 
      facet_grid(.~type) + labs(color=paste0(parasweep[1],"_value"),linetype="Habitat Loss") + 
      ylab("Population Density N2")
    png(filename = paste0(foldername,"/",parasweep[1],"/plot_HLcomp/N2_HLvsNHL.png"), width = 1280, height = 720)
    print(pN2); dev.off()
    
    #Plot resource comparison for HL and NHL
    pR1 <- p + geom_line(aes(y=R1, color=param_val, linetype=HL), linewidth=1.2) + 
      facet_grid(.~type) + labs(color=paste0(parasweep[1],"_value"),linetype="Habitat Loss") + 
      ylab("Resource Density R1")
    png(filename = paste0(foldername,"/",parasweep[1],"/plot_HLcomp/R1_HLvsNHL.png"), width = 1280, height = 720)
    print(pR1); dev.off()
    pR2 <- p + geom_line(aes(y=R2, color=param_val, linetype=HL), linewidth=1.2) + 
      facet_grid(.~type) + labs(color=paste0(parasweep[1],"_value"),linetype="Habitat Loss") + 
      ylab("Resource Density R2")
    png(filename = paste0(foldername,"/",parasweep[1],"/plot_HLcomp/R2_HLvsNHL.png"), width = 1280, height = 720)
    print(pR2); dev.off()
  }
  else {
    stop("Currently this works when only one parameter is varied")
  }
}

#Function to plot HL-NHL comparison for N1+N2 and R1+R2 (raw plots, not directly used in manuscript)
plot_LandscapeHLcomp <- function(foldername,parasweep=c()) {
  if (length(parasweep)==0) {
    load(paste0(foldername,"/rawdata/file.RData"))
    if (!dir.exists(paste0(foldername,"/plot_LandscapeHLcomp"))){
      dir.create(paste0(foldername,"/plot_LandscapeHLcomp"))
    }
    cases <- expand.grid(Va=c(T,F),HL=c(T,F),VM=c(T,F))
    cases <- cases[!(cases[,"Va"]==F & cases[,"VM"]==T),]
    # Habitat loss data
    HLData <- data.frame()
    for (i in c(1:nrow(cases))[cases[,"HL"]]) { #comparison in presence of HL
      len <- nrow(obj[[i]]$params)
      tempData <- data.frame(cbind(t=obj[[i]]$out[,1],
                                   N=rowSums(obj[[i]]$out[, 2:(len+1), drop=F])+rowSums(obj[[i]]$out[,(len+2):(2*len+1), drop=F]),
                                   R=obj[[i]]$out[, (2*len+2)]+obj[[i]]$out[, (2*len+3)]))
      tempData$type <- if (cases[i,"VM"]==TRUE) "Constant Variation" else if (cases[i,"Va"]==FALSE) "No Variation" else "Heritable Variation"
      HLData <- rbind(HLData,tempData)
    }
    HLData$type <- factor(HLData$type,levels=c("No Variation","Heritable Variation","Constant Variation"))
    HLData$HL <- "HL patch 2"
    
    # No Habitat loss data
    NHLData <- data.frame()
    for (i in c(1:nrow(cases))[!cases[,"HL"]]) { #comparison in absence of HL
      len <- nrow(obj[[i]]$params)
      tempData <- data.frame(cbind(t=obj[[i]]$out[,1],
                                   N=rowSums(obj[[i]]$out[, 2:(len+1), drop=F])+rowSums(obj[[i]]$out[,(len+2):(2*len+1), drop=F]),
                                   R=obj[[i]]$out[, (2*len+2)]+obj[[i]]$out[, (2*len+3)]))
      tempData$type <- if (cases[i,"VM"]==TRUE) "Constant Variation" else if (cases[i,"Va"]==FALSE) "No Variation" else "Heritable Variation"
      NHLData <- rbind(NHLData,tempData)
    }
    NHLData$type <- factor(NHLData$type,levels=c("No Variation","Heritable Variation","Constant Variation"))
    NHLData$HL <- "No HL"
    
    Data <- rbind(HLData,NHLData)
    
    #The basic ggplot object
    p <- ggplot(data=Data,aes(x=t)) + xlab("Time") + 
      scale_color_manual(values = c("No Variation"="#56B4E9", "Heritable Variation"="#F0E442",
                                    "Constant Variation"="#D55E00")) +
      scale_linetype_manual(breaks = c("No HL", "HL patch 2"),
                            values = c("No HL"="solid", "HL patch 2"="dotted")) +
      theme_bw() + theme(plot.title = element_text(hjust = 0.5, size = 20),
                         axis.title.x = element_text(size = 15),
                         axis.title.y = element_text(size = 15),
                         axis.text = element_text(size = 10, face = "bold"),
                         legend.title = element_text(size = 15),
                         legend.text = element_text(size = 10),
                         strip.text = element_text(size = 12),
                         panel.grid.major = element_blank(),
                         panel.grid.minor = element_blank(),
                         strip.background = element_rect(colour="black", fill="white"),
                         panel.border = element_rect(colour = "black"),
                         panel.spacing = unit(0, "lines")) +
      scale_x_continuous(expand = c(0,0)) + scale_y_continuous(expand = c(0,0))
    
    #Plot landscape population density comparison for HL and NHL
    pN <- p + geom_line(aes(y=N, color=type, linetype=HL), linewidth=1.2) + 
      labs(color="Trait Variation",linetype="Habitat Loss") + 
      ylab("Population Density N1+N2") + ggtitle("Landscape Population Density HL vs NHL")
    png(filename = paste0(foldername,"/plot_LandscapeHLcomp/N_HLvsNHL.png"), width = 852, height = 480)
    print(pN); dev.off()
    
    #Plot resource comparison for HL and NHL
    pR <- p + geom_line(aes(y=R, color=type, linetype=HL), linewidth=1.2) + 
      labs(color="Trait Variation",linetype="Habitat Loss") + 
      ylab("Resource Density R1+R2") + ggtitle("Landscape Resource Density HL vs NHL")
    png(filename = paste0(foldername,"/plot_LandscapeHLcomp/R_HLvsNHL.png"), width = 852, height = 480)
    print(pR); dev.off()
  }
  else if (length(parasweep)==1) {
    load(paste0(foldername,"/",parasweep[1],"/rawdata/file.RData"))
    if (!dir.exists(paste0(foldername,"/",parasweep[1],"/plot_LandscapeHLcomp"))) {
      dir.create(paste0(foldername,"/",parasweep[1],"/plot_LandscapeHLcomp"))
    }
    cases <- expand.grid(Va=c(T,F),HL=c(T,F),VM=c(T,F))
    cases <- cases[!(cases[,"Va"]==F & cases[,"VM"]==T),]
    
    Data <- data.frame()
    for (j in 1:length(obj)) {
      for (i in 1:nrow(cases)) {
        obj1<-obj[[j]]
        len <- nrow(obj1[[i]]$params)
        tempData <- data.frame(cbind(t=obj1[[i]]$out[,1],
                                     N=rowSums(obj1[[i]]$out[, 2:(len+1), drop=F])+rowSums(obj1[[i]]$out[,(len+2):(2*len+1), drop=F]),
                                     R=obj1[[i]]$out[, (2*len+2)]+obj1[[i]]$out[, (2*len+3)]))
        tempData$type <- if (cases[i,"VM"]==TRUE) "Constant Variation" else if (cases[i,"Va"]==FALSE) "No Variation" else "Heritable Variation"
        #paste0("Va_",cases[i,"Va"],"_VM_",cases[i,"VM"])
        tempData$HL <- if (cases[i,"HL"]==TRUE) "HL patch 2" else "No HL"
        tempData$param_val <- names(obj)[j]
        Data <- rbind(Data,tempData)
      }
    }
    Data$type <- factor(Data$type,levels=c("No Variation","Heritable Variation","Constant Variation"))
    Data$param_val <- factor(Data$param_val,levels=c(names(obj)))
    
    #The basic ggplot object
    p <- ggplot(data=Data,aes(x=t)) + xlab("Time") + 
      scale_linetype_manual(breaks = c("No HL", "HL patch 2"),
                            values = c("No HL"="solid", "HL patch 2"="dotted")) +
      ggtitle(paste("Changing parameter",parasweep[1],"Landscape HL vs NHL")) + 
      theme_bw() + theme(plot.title = element_text(hjust = 0.5, size = 20),
                         axis.title.x = element_text(size = 15),
                         axis.title.y = element_text(size = 15),
                         axis.text = element_text(size = 10, face = "bold"),
                         legend.title = element_text(size = 15),
                         legend.text = element_text(size = 10),
                         legend.position = "bottom",
                         strip.text = element_text(size = 12),
                         panel.grid.major = element_blank(),
                         panel.grid.minor = element_blank(),
                         strip.background = element_rect(colour="black", fill="white"),
                         panel.border = element_rect(colour = "black"),
                         panel.spacing = unit(1, "lines")) +
      scale_x_continuous(expand = c(0,0)) + scale_y_continuous(expand = c(0,0))
    
    #Plot landscape population density comparison for HL and NHL
    pN <- p + geom_line(aes(y=N, color=param_val, linetype=HL), linewidth=1.2) + 
      facet_grid(.~type) + labs(color=paste0(parasweep[1],"_value"),linetype="Habitat Loss") + 
      ylab("Population Density N1+N2")
    png(filename = paste0(foldername,"/",parasweep[1],"/plot_LandscapeHLcomp/N1_HLvsNHL.png"), width = 1280, height = 720)
    print(pN); dev.off()
    
    #Plot resource comparison for HL and NHL
    pR <- p + geom_line(aes(y=R, color=param_val, linetype=HL), linewidth=1.2) + 
      facet_grid(.~type) + labs(color=paste0(parasweep[1],"_value"),linetype="Habitat Loss") + 
      ylab("Resource Density R1+R2")
    png(filename = paste0(foldername,"/",parasweep[1],"/plot_LandscapeHLcomp/R1_HLvsNHL.png"), width = 1280, height = 720)
    print(pR); dev.off()
  }
  else {
    stop("Currently this works when only one parameter is varied")
  }
}

#timeseries plot for multcases (raw plots, not directly used in manuscript)
timeseriesplot_multcases <- function(foldername,i,obj){
  out <- obj[[i]]$out
  len <- length(obj[[i]]$params[,1])
  if (dir.exists(paste0(foldername,"/plots"))==F) {
    dir.create(paste0(foldername,"/plots"))
  }
  png(filename = paste0(foldername,"/plots/population_timeseries.png"), width = 852, height = 480)
  matplot(out[ ,1], cbind(rowSums(out[, 2:(len+1), drop=F]),
                          rowSums(out[,(len+2):(2*len+1), drop=F])), type = "l",
          xlab = "time", ylab = "Density", main = "Density time series", lwd = 2,
          xaxs="i", yaxs="i")
  legend("topright", c("N1", "N2"), col = 1:2, lty = 1:2)
  dev.off()
  png(filename = paste0(foldername,"/plots/resource_timeseries.png"), width = 852, height = 480)
  matplot(out[, 1], out[, (2*len+2):(2*len+3)], type = "l", xlab = "time", ylab = "Resource",
          main = "Respource time series", lwd = 2, xaxs="i", yaxs="i")
  legend("topright", c("R1", "R2"), col = 1:2, lty = 1:2)
  dev.off()
}

#Traits distribution over time for multiple cases (raw plots, not directly used in manuscript)
traitsovertime_multcases <- function(foldername,j,traits,obj) {
  out <- obj[[j]]$out
  p <- obj[[j]]$params
  len <- length(p[,1])
  times <- out[,1]
  if (dir.exists(paste0(foldername,"/traitsovertime"))==F) {
    dir.create(paste0(foldername,"/traitsovertime"))
  }
  outdf <- expand.grid(tm.i=1:length(times),zvals.i=1:len)
  for (i in traits){
    outdf1 <- outdf
    outdf1$patch <- "N1"
    outdf1$freqST <- as.numeric(paste0(t(scale(t(out[,2:(len+1)]),center=F,scale=colSums(t(out[,2:(len+1)]))))))
    outdf1$freqSF <- as.numeric(out[,2:(len+1)]) #Scaled Falls (SF), Scaled True (ST)
    outdf2 <- outdf
    outdf2$patch <- "N2"
    outdf2$freqST <- as.numeric(paste0(t(scale(t(out[,(len+2):(2*len+1)]),center=F,scale=colSums(t(out[,(len+2):(2*len+1)]))))))
    outdf2$freqSF <- as.numeric(out[,(len+2):(2*len+1)]) #Scaled Falls (SF), Scaled True (ST)
    totout <- rbind(outdf1,outdf2)  
    totout$z <- as.numeric(p[,i])[totout$zvals.i]
    totout$z <- round(totout$z,2)
    totout$tm <- as.numeric(times)[totout$tm.i]
    
    # yvals <- round(as.numeric(p[,i]),2)
    # yvals2 <- c(0,yvals,max(yvals))
    # yheight <- diff((yvals2[-1]+yvals2[-length(yvals2)]))
    # totout$ht <- as.numeric(sapply(totout$z, FUN = function(x) yheight[which(x==yvals)]))
    
    # generate plotting object ST
    pST <- ggplot(totout,aes(x=tm,y=z,fill=freqST)) + geom_tile(stat="identity") +
      facet_grid(.~patch) + scale_fill_gradient(low="white",high="black") + 
      xlab("Time") + ylab(paste("Trait",i)) + labs(fill = "Scaled density") +
      theme_bw() + theme(panel.grid.major = element_blank(),
                         panel.grid.minor = element_blank(),
                         strip.background = element_rect(colour="black", fill="white"),
                         panel.spacing.x = unit(c(0.7), "lines"),
                         axis.text.x = element_text(angle=90, vjust=0.5)) +
      scale_x_continuous(expand = c(0,0), guide = guide_axis(check.overlap = T)) + 
      scale_y_continuous(expand = c(0,0))
    
    # generate plotting object SF
    pSF <- ggplot(totout,aes(x=tm,y=z,fill=freqSF)) + geom_tile(stat="identity") +
      facet_grid(.~patch) + scale_fill_gradient(low="white",high="black") + 
      xlab("Time") + ylab(paste("Trait",i)) + labs(fill = "Absolute density") +
      theme_bw() + theme(panel.grid.major = element_blank(),
                         panel.grid.minor = element_blank(),
                         strip.background = element_rect(colour="black", fill="white"),
                         panel.spacing.x = unit(c(0.7), "lines"),
                         axis.text.x = element_text(angle=90, vjust=0.5)) +
      scale_x_continuous(expand = c(0,0), guide = guide_axis(check.overlap = T)) + 
      scale_y_continuous(expand = c(0,0))
    
    #Save the plot 
    png(filename = paste0(foldername,"/traitsovertime/",i,"_abs_traitdistribution_overtime.png"), width = 852, height = 480)
    print(pSF); dev.off()
    png(filename = paste0(foldername,"/traitsovertime/",i,"_scaled_traitdistribution.png"), width = 852, height = 480)
    print(pST); dev.off() 
  }
}

#Trade-off plot (raw plots, not directly used in manuscript)
traitrelplot_multcases <- function(foldername, traits, obj) {
  p <- obj[[1]]$params
  len <- length(p[,1])
  if (dir.exists(paste0(foldername,"/tradeoff"))==F) {
    dir.create(paste0(foldername,"/tradeoff"))
  }
  png(filename = paste0(foldername,"/tradeoff/",traits[2],"Vs",traits[1],".png"), width = 852, height = 480)
  plot(p[,traits[1]], p[,traits[2]], xlab = traits[1], ylab = traits[2],
       main = "Trade-off")
  dev.off()
}

#Habita Loss shape (sigmoid, dependent on d, t0, and k) (raw plots, not directly used in manuscript)
HLshape_multcases <- function(foldername, obj) {
  p <- obj[[1]]$params
  times <- obj[[1]]$out[,1]
  Res.red.rate <- p[1,"d"]/(1+exp(-p[1,"k"]*(times-p[1,"t0"]))) #Rate of resource removal over time
  if (dir.exists(paste0(foldername,"/HL_Shape"))==F) {
    dir.create(paste0(foldername,"/HL_Shape"))
  }
  png(filename = paste0(foldername,"/HL_Shape/","HL_Shape_","d_",p[1,"d"],"_t0_",p[1,"t0"],
                        "_k_",p[1,"k"],".png"), width = 852, height = 480)
  plot(times, Res.red.rate, xlab = "time", ylab = "Resource removal rate term",
       main = paste0("HL_Shape_","d_",p[1,"d"],"_t0_",p[1,"t0"],"_k_",p[1,"k"]))
  dev.off()
}
    
```
