## Supplementary material for "Multi-scale Effects of Habitat Loss and the Role of Trait Variation": R code for generating data and figures: Figure_2_plotfile.R

```
# Here we compare the two patches with and without habitat loss for no variation cases
rm(list = ls())
library(ggplot2)
library(ggpubr)
library(latex2exp) #useful to use latex symbols
loadRData <- function(fileName){ #loads an RData file, and returns it
  load(fileName)
  get(ls()[ls() != "fileName"])
}

###################### Preppeing Data ########################
OE.Data <- loadRData("Data/e-var-mean_0p2-sd_0p08-a0_0p1/rawdata/file.RData") #Over Exploitation Data
NE.Data <- loadRData("Data/e-var-mean_0p2-sd_0p08-a0_0p14/rawdata/file.RData") #Normal Exploitation Data
filname <- "Figure_2.pdf"

cases <- expand.grid(Va=c(T,F),HL=c(T,F),VM=c(T,F))
cases <- cases[!(cases[,"Va"]==F & cases[,"VM"]==T),]

# Over Exploitation data
DataOE <- data.frame()
for (i in c(1:nrow(cases))) { #comparison in presence of HL
  len <- nrow(OE.Data[[i]]$params)
  tempData <- data.frame(cbind(t=OE.Data[[i]]$out[,1], N1=rowSums(OE.Data[[i]]$out[, 2:(len+1), drop=F]),
                               N2=rowSums(OE.Data[[i]]$out[,(len+2):(2*len+1), drop=F]),
                               R1=OE.Data[[i]]$out[, (2*len+2)],R2=OE.Data[[i]]$out[, (2*len+3)]))
  tempData$type <- if (cases[i,"VM"]==TRUE) "Constant Variation" else if (cases[i,"Va"]==FALSE) "No Variation" else "Heritable Variation"
  tempData$HL <- if (cases[i,"HL"]==TRUE) "Habitat Loss" else "No Habitat Loss"
  DataOE <- rbind(DataOE,tempData)
}
DataOE$type <- factor(DataOE$type,levels=c("No Variation","Heritable Variation","Constant Variation"))
DataOE$HL <- factor(DataOE$HL, levels = c("No Habitat Loss", "Habitat Loss"))

# Normal Exploitation data
DataNE <- data.frame()
for (i in c(1:nrow(cases))) { #comparison in presence of HL
  len <- nrow(NE.Data[[i]]$params)
  tempData <- data.frame(cbind(t=NE.Data[[i]]$out[,1], N1=rowSums(NE.Data[[i]]$out[, 2:(len+1), drop=F]),
                               N2=rowSums(NE.Data[[i]]$out[,(len+2):(2*len+1), drop=F]),
                               R1=NE.Data[[i]]$out[, (2*len+2)],R2=NE.Data[[i]]$out[, (2*len+3)]))
  tempData$type <- if (cases[i,"VM"]==TRUE) "Constant Variation" else if (cases[i,"Va"]==FALSE) "No Variation" else "Heritable Variation"
  tempData$HL <- if (cases[i,"HL"]==TRUE) "Habitat Loss" else "No Habitat Loss"
  DataNE <- rbind(DataNE,tempData)
}
DataNE$type <- factor(DataNE$type,levels=c("No Variation","Heritable Variation","Constant Variation"))
DataNE$HL <- factor(DataNE$HL, levels = c("No Habitat Loss", "Habitat Loss"))

rm("NE.Data","OE.Data","tempData")

################# OE Plots ######################
p.OE <- ggplot(data=DataOE[DataOE$type=="No Variation",],aes(x=t)) + xlab("Time") + 
  scale_color_manual("",values = c("No Habitat Loss"="blue", "Habitat Loss"="red")) +
  scale_linetype_manual("Patch",values=c("1"="solid","2"="dotted")) +
  theme_bw() + theme(#plot.title = element_text(hjust = 0.5, size = 10),
    # axis.title.x = element_text(size = 15),
    # axis.title.y = element_text(size = 8),
    axis.text = element_text(size = 7, face = "bold"),
    # legend.title = element_text(size = 15),
    # legend.text = element_text(size = 10),
    legend.position="bottom",
    # strip.text = element_text(size = 12),
    plot.margin = margin(0.5, 0.5, 0.5, 0.5, "lines"),
    panel.grid.major = element_blank(),panel.grid.minor = element_blank(),
    strip.background = element_rect(colour="black", fill="white"),
    panel.border = element_rect(colour = "black"),
    panel.spacing = unit(0, "lines")) +
  scale_x_continuous(expand = c(0,0))# + scale_y_continuous(expand = c(0,0),limits = c(0,10))

pN.OE <- p.OE + geom_line(aes(y=N1, color=HL, linetype="1"), linewidth=0.5, alpha=0.7) + 
  geom_line(aes(y=N2, color=HL, linetype="2"), linewidth=0.5) +
  ylab("Consumer Density") + scale_y_continuous(expand = c(0,0),limits = c(0,20))
#pN.OE

pR.OE <- p.OE + geom_line(aes(y=R1, color=HL, linetype="1"), linewidth=0.5, alpha=0.7) + 
  geom_line(aes(y=R2, color=HL, linetype="2"), linewidth=0.5) +
  ylab("Resource Density") + scale_y_continuous(expand = c(0,0),limits = c(0,50))
#pR.OE

####################### NE Plots ##########################
p.NE <- ggplot(data=DataNE[DataNE$type=="No Variation",],aes(x=t)) + xlab("Time") + 
  scale_color_manual("HL Type",values = c("No Habitat Loss"="blue", "Habitat Loss"="red")) +
  scale_linetype_manual("Patch",values=c("1"="solid","2"="dotted")) +
  theme_bw() + theme(#plot.title = element_text(hjust = 0.5, size = 10),
    # axis.title.x = element_text(size = 15),
    # axis.title.y = element_text(size = 8),
    axis.text = element_text(size = 7, face = "bold"),
    # legend.title = element_text(size = 15),
    # legend.text = element_text(size = 10),
    legend.position="bottom",
    # strip.text = element_text(size = 12),
    plot.margin = margin(0.5, 0.5, 0.5, 0.5, "lines"),
    panel.grid.major = element_blank(),panel.grid.minor = element_blank(),
    strip.background = element_rect(colour="black", fill="white"),
    panel.border = element_rect(colour = "black"),
    panel.spacing = unit(0, "lines")) +
  scale_x_continuous(expand = c(0,0))# + scale_y_continuous(expand = c(0,0),limits = c(0,10))

pN.NE <- p.NE + geom_line(aes(y=N1, color=HL, linetype="1"), linewidth=0.5, alpha=0.7) + 
  geom_line(aes(y=N2, color=HL, linetype="2"), linewidth=0.5) +
  ylab("Consumer Density") + scale_y_continuous(expand = c(0,0),limits = c(0,20))
#pN.NE

pR.NE <- p.NE + geom_line(aes(y=R1, color=HL, linetype="1"), linewidth=0.5, alpha=0.7) + 
  geom_line(aes(y=R2, color=HL, linetype="2"), linewidth=0.5) +
  ylab("Resource Density") + scale_y_continuous(expand = c(0,0),limits = c(0,50))
#pR.NE

############## Landscape level density plots ###########################
pNL.OE <- p.OE + geom_line(aes(y=N1+N2, color=HL), linewidth=0.5, alpha=0.7) +
  ylab("Total Consumer Density") + scale_y_continuous(expand = c(0,0),limits = c(0,30))
#pNL.OE

pRL.OE <- p.OE + geom_line(aes(y=R1+R2, color=HL), linewidth=0.5, alpha=0.7) +
  ylab("Total Resource Density") + scale_y_continuous(expand = c(0,0),limits = c(0,60))
#pRL.OE

pNL.NE <- p.NE + geom_line(aes(y=N1+N2, color=HL), linewidth=0.5, alpha=0.7) +
  ylab("Total Consumer Density") + scale_y_continuous(expand = c(0,0),limits = c(0,30))
#pNL.NE

pRL.NE <- p.NE + geom_line(aes(y=R1+R2, color=HL), linewidth=0.5, alpha=0.7) +
  ylab("Total Resource Density") + scale_y_continuous(expand = c(0,0),limits = c(0,60))
#pRL.NE


################### Making Final Plot ##########################
p <- ggarrange(pN.OE+theme(plot.title = element_text(hjust = 0.5,size = 10),axis.title.x = element_blank(),axis.ticks.x = element_blank(),axis.text.x = element_blank())+ggtitle("Patch-wise Consumer")+ylab("Density"),
               pR.OE+theme(plot.title = element_text(hjust = 0.5,size = 10),axis.title.x = element_blank(),axis.ticks.x = element_blank(),axis.text.x = element_blank(), axis.title.y = element_blank())+ggtitle("Patch-wise Resource"),
               pNL.OE+theme(plot.title = element_text(hjust = 0.5,size = 10),axis.title.x = element_blank(),axis.ticks.x = element_blank(),axis.text.x = element_blank(), axis.title.y = element_blank())+ggtitle("Total Consumer"),
               pRL.OE+theme(plot.title = element_text(hjust = 0.5,size = 10),axis.title.x = element_blank(),axis.ticks.x = element_blank(),axis.text.x = element_blank(), axis.title.y = element_blank())+ggtitle("Total Resource"),
               pN.NE+theme(plot.title = element_blank(),axis.text.x = element_text(angle = 90, vjust = 0.5),axis.title.x = element_blank())+ylab("Density"),
               pR.NE+theme(plot.title = element_blank(),axis.text.x = element_text(angle = 90, vjust = 0.5),axis.title.x = element_blank(), axis.title.y = element_blank()),
               pNL.NE+theme(plot.title = element_blank(),axis.text.x = element_text(angle = 90, vjust = 0.5),axis.title.x = element_blank(), axis.title.y = element_blank()),
               pRL.NE+theme(plot.title = element_blank(),axis.text.x = element_text(angle = 90, vjust = 0.5),axis.title.x = element_blank(), axis.title.y = element_blank()),
               labels = "auto",
               ncol = 4, nrow = 2, align = "v", vjust = 1.5, hjust = -1,
               common.legend = TRUE, legend = "none")
p <- annotate_figure(p, bottom = text_grob("Time", size = 10))
p <- annotate_figure(p, bottom = get_legend(pN.OE))
p <- annotate_figure(p, left = text_grob(TeX("$a_0 = 0.14$                  $a_0 = 0.1$"), rot = 90, hjust=0.32, face = "bold"))
ggsave(filename = filname, device = "pdf", plot = p, width = 8, height = 4)
#p
```
