## Supplementary material for "Multi-scale Effects of Habitat Loss and the Role of Trait Variation": R code for generating data and figures: Figure_4_plotfile.R

```
# N* and R* at different values of d when there is habitat loss,
# compared when patches are dependent or independent

rm(list = ls())
library(ggplot2)
library(ggpubr)
library(latex2exp) #useful to use latex symbols
library(reshape2) #for reshaping dataframes
loadRData <- function(fileName){ #loads an RData file, and returns it
  load(fileName)
  get(ls()[ls() != "fileName"])
}

###################### Preppeing Data ########################
filname <- "Figure_4.pdf"

parsweep.curator <- function(filepath, trait) {
  obj <- loadRData(filepath)
  
  cases <- expand.grid(Va=c(T,F),HL=c(T,F),VM=c(T,F))
  cases <- cases[!(cases[,"Va"]==F & cases[,"VM"]==T),]
  
  Data <- data.frame()
  for (j in 1:length(obj)) {
    for (i in 1:nrow(cases)) {
      obj1<-obj[[j]]
      len <- nrow(obj1[[i]]$params)
      tempData <- data.frame(cbind(N1=mean(tail(rowSums(obj1[[i]]$out[, 2:(len+1), drop=F]),n=1)),
                                   N2=mean(tail(rowSums(obj1[[i]]$out[,(len+2):(2*len+1), drop=F]),n=1)),
                                   R1=mean(tail(obj1[[i]]$out[, (2*len+2)],n=1)),
                                   R2=mean(tail(obj1[[i]]$out[, (2*len+3)],n=1))))
      tempData$type <- if (cases[i,"VM"]==TRUE) "Constant Variation" else if (cases[i,"Va"]==FALSE) "No Variation" else "Heritable Variation"
      tempData$HL <- if (cases[i,"HL"]==TRUE) "Habitat Loss" else "No Habitat Loss"
      tempData$param_val <- as.numeric(names(obj)[j])
      Data <- rbind(Data,tempData)
    }
  }
  Data$Trait <- trait
  Data$Trait <- factor(Data$Trait)
  Data$type <- factor(Data$type,levels=c("No Variation","Heritable Variation","Constant Variation"))
  Data$HL <- factor(Data$HL, levels = c("No Habitat Loss", "Habitat Loss"))
  return(Data)
}

Data.PD <- parsweep.curator("Data/e-var-mean_0p2-sd_0p08-parsweep_d/d/rawdata/file.RData","PD")
Data.noPD <- parsweep.curator("Data/e-var-mean_0p2-sd_0p08-parsweep_d-noPD/d/rawdata/file.RData","noPD")
Data <- rbind(Data.PD,Data.noPD)
levels(Data$Trait) <- list(Connected="PD", Isolated="noPD")
Data <- Data[Data$HL=="Habitat Loss",]
rm(Data.PD,Data.noPD)

pobj <- ggplot(data=Data,aes(x=param_val)) + xlab("D") +
  scale_color_manual("Patches",values=c("Connected"="turquoise","Isolated"="orange")) +
  scale_linetype_manual("",values = c("No Variation"="solid", "Heritable Variation"="dotted",
                                      "Constant Variation"="dashed")) +
  scale_alpha_manual(values = c("No Variation"=0.7, "Heritable Variation"=1,
                                "Constant Variation"=1),guide = "none") +
  theme_bw() + theme(plot.title = element_text(hjust = 0.5),
                     axis.title.x = element_text(face = "bold"),
                     axis.title.y = element_text(size=8,face = "bold"),
                     axis.text.x = element_text(angle=90, vjust=0.5),
                     #legend.title = element_text(size = 10),
                     #legend.text = element_text(size = 7),
                     legend.position="bottom",
                     # strip.text = element_text(size = 12),
                     plot.margin = margin(0.5, 1, 0.5, 0.5, "lines"),
                     panel.grid.major = element_blank(),panel.grid.minor = element_blank(),
                     strip.background = element_blank(),
                     strip.placement = "outside",
                     panel.border = element_rect(colour = "black"),
                     panel.spacing = unit(0.7, "lines")) +
  scale_x_continuous(expand = c(0,0.0001))# + scale_y_continuous(expand = c(0,0),limits = c(0,10))

pN1.OE <- pobj + geom_line(aes(y=N1, color=Trait, linetype=type, alpha=type), linewidth=0.5) +
  ylab("Equilibrium Consumer Density") + scale_y_continuous(expand = c(0,0.1),limits = c(-0.01,25))
#pN1.OE

pN2.OE <- pobj + geom_line(aes(y=N2, color=Trait, linetype=type, alpha=type), linewidth=0.5) +
  ylab("Equilibrium Consumer Density") + scale_y_continuous(expand = c(0,0.1),limits = c(-0.01,25))
#pN2.OE

pN.OE <- pobj + geom_line(aes(y=N1+N2, color=Trait, linetype=type, alpha=type), linewidth=0.5) +
  ylab("Equilibrium Consumer Density") + scale_y_continuous(expand = c(0,0.1),limits = c(-0.01,40))
#pN.OE

pR1.OE <- pobj + geom_line(aes(y=R1, color=Trait, linetype=type, alpha=type), linewidth=0.5) +
  ylab("Equilibrium Resource Density") + scale_y_continuous(expand = c(0,0.1),limits = c(-0.01,35))
#pR1.OE

pR2.OE <- pobj + geom_line(aes(y=R2, color=Trait, linetype=type, alpha=type), linewidth=0.5) +
  ylab("Equilibrium Resource Density") + scale_y_continuous(expand = c(0,0.1),limits = c(-0.01,35))
#pR2.OE

pR.OE <- pobj + geom_line(aes(y=R1+R2, color=Trait, linetype=type, alpha=type), linewidth=0.5) +
  ylab("Equilibrium Resource Density") + scale_y_continuous(expand = c(0,0.1),limits = c(-0.01,60))
#pR.OE

################### Making Final Plot ##########################
figure <- ggarrange(pN1.OE+theme(axis.title.x = element_blank(),axis.ticks.x=element_blank(),axis.text.x = element_blank())+ggtitle("Patch 1"),
                    pN2.OE+theme(axis.title.x = element_blank(),axis.ticks.x=element_blank(),axis.text.x = element_blank(),axis.title.y = element_blank())+ggtitle("Patch 2"),
                    pN.OE+theme(axis.title.x = element_blank(),axis.ticks.x=element_blank(),axis.text.x = element_blank(),axis.title.y = element_blank())+ggtitle("Landscape"),
                    pR1.OE+theme(axis.title.y = element_text(),axis.title.x = element_blank(),plot.title = element_blank()),
                    pR2.OE+theme(axis.title.x = element_blank(),axis.title.y = element_blank(),plot.title = element_blank()),
                    pR.OE+theme(axis.title.x = element_blank(),axis.title.y = element_blank(),plot.title = element_blank()),
                    labels = "auto", heights = c(1,0.98),
                    ncol = 3, nrow = 2, align = "v", vjust = 1,# hjust = -1,
                    common.legend = TRUE, legend = "none")

figure <- annotate_figure(figure, bottom = text_grob("D", size = 12))
figure <- annotate_figure(figure, bottom = get_legend(pN1.OE))

ggsave(filename = filname, device = "pdf", plot = figure, width = 8, height = 6)


# # Code for the above plot with only no variation case + resource plot
# filname <- "Fig-parsweep_d-HL-PD_NR_comp.pdf"
# pobj <- ggplot(data=Data[Data$type=="No Variation",],aes(x=param_val)) + xlab("D") +
#   scale_linetype_manual("Patches",values=c("Dependent"="solid","Independent"="dotted")) +
#   scale_color_manual("",values=c("Consumer"="orange","Resource"="green")) +
#   theme_bw() + theme(plot.title = element_text(hjust = 0.5),
#                      axis.title.x = element_text(face = "bold"),
#                      axis.title.y = element_text(size=9,face = "bold"),
#                      axis.text.x = element_text(angle=90, vjust=0.5),
#                      #legend.title = element_text(size = 10),
#                      #legend.text = element_text(size = 7),
#                      legend.position="bottom",
#                      # strip.text = element_text(size = 12),
#                      plot.margin = margin(0.5, 1, 0.5, 0.5, "lines"),
#                      panel.grid.major = element_blank(),panel.grid.minor = element_blank(),
#                      strip.background = element_blank(),
#                      strip.placement = "outside",
#                      panel.border = element_rect(colour = "black"),
#                      panel.spacing = unit(0.7, "lines")) +
#   scale_x_continuous(expand = c(0,0.0001))# + scale_y_continuous(expand = c(0,0),limits = c(0,10))
# 
# pN1.OE <- pobj + geom_line(aes(y=N1, linetype=Trait, color="Consumer"), linewidth=0.5) +
#   geom_line(aes(y=R1, linetype=Trait, color="Resource"), linewidth=0.5) +
#   ylab("Equilibrium Density (HL case)") + scale_y_continuous(expand = c(0,0),limits = c(-0.01,35))
# #pN1.OE
# 
# pN2.OE <- pobj + geom_line(aes(y=N2, linetype=Trait, color="Consumer"), linewidth=0.5) +
#   geom_line(aes(y=R2, linetype=Trait, color="Resource"), linewidth=0.5) +
#   ylab("Equilibrium Density (HL case)") + scale_y_continuous(expand = c(0,0),limits = c(-0.01,35))
# #pN2.OE
# 
# pN.OE <- pobj + geom_line(aes(y=N1+N2, linetype=Trait, color="Consumer"), linewidth=0.5) +
#   geom_line(aes(y=R1+R2, linetype=Trait, color="Resource"), linewidth=0.5) +
#   ylab("Equilibrium Density (HL case)") + scale_y_continuous(expand = c(0,0),limits = c(-0.01,60))
# #pN.OE
# 
# ################### Making Final Plot ##########################
# figure <- ggarrange(pN1.OE+theme(axis.title.x = element_blank())+ggtitle(TeX("Patch 1")),
#                     pN2.OE+theme(axis.title.y = element_blank())+ggtitle(TeX("Patch 2")),
#                     pN.OE+theme(axis.title.x = element_blank(),axis.title.y = element_blank())+ggtitle(TeX("Landscape")),
#                     labels = "auto",
#                     ncol = 3, nrow = 1, align = "hv", vjust = 1,# hjust = -1,
#                     common.legend = TRUE, legend = "bottom")
# 
# ggsave(filename = filname, device = "pdf", plot = figure, width = 8, height = 3.5)
```
