## Supplementary material for "Multi-scale Effects of Habitat Loss and the Role of Trait Variation": R code for generating data and figures: Figure_5_plotfile.R

```
# Here we compare the two patches with and without habitat loss for different variation cases
# and when the traits p and beta are traded off positively
rm(list = ls())
library(ggplot2)
library(ggpubr)
library(latex2exp) #useful to use latex symbols
loadRData <- function(fileName){ #loads an RData file, and returns it
  load(fileName)
  get(ls()[ls() != "fileName"])
}

###################### Preppeing Data ########################
OE.Data <- loadRData("Data/p-beta-pos-tradeoff-mean_0p65-N0_switched-a0_0p1/rawdata/file.RData") #Over Exploitation Data
ONE.Data <- loadRData("Data/p-beta-pos-tradeoff-mean_0p65-N0_switched-a0_0p15/rawdata/file.RData")
NE.Data <- loadRData("Data/p-beta-pos-tradeoff-mean_0p65-N0_switched-a0_0p18/rawdata/file.RData") #Normal Exploitation Data
filname <- "Figure_5.pdf"
trait <- "p1"

cases <- expand.grid(Va=c(T,F),HL=c(T,F),VM=c(T,F))
cases <- cases[!(cases[,"Va"]==F & cases[,"VM"]==T),]

# low a0 value
DataOE <- data.frame()
for (i in c(1:nrow(cases))) { #comparison in presence of HL
  len <- nrow(OE.Data[[i]]$params)
  tempData <- data.frame(cbind(t=OE.Data[[i]]$out[,1], N1=rowSums(OE.Data[[i]]$out[, 2:(len+1), drop=F]),
                               N2=rowSums(OE.Data[[i]]$out[,(len+2):(2*len+1), drop=F]),
                               R1=OE.Data[[i]]$out[, (2*len+2)],R2=OE.Data[[i]]$out[, (2*len+3)]))
  tempData$type <- if (cases[i,"VM"]==TRUE) "Constant Variation" else if (cases[i,"Va"]==FALSE) "No Variation" else "Heritable Variation"
  tempData$HL <- if (cases[i,"HL"]==TRUE) "Habitat Loss" else "No Habitat Loss"
  DataOE <- rbind(DataOE,tempData)
}
DataOE$type <- factor(DataOE$type,levels=c("No Variation","Heritable Variation","Constant Variation"))
DataOE$HL <- factor(DataOE$HL, levels = c("No Habitat Loss", "Habitat Loss"))

# Middle a0 value
DataONE <- data.frame()
for (i in c(1:nrow(cases))) { #comparison in presence of HL
  len <- nrow(ONE.Data[[i]]$params)
  tempData <- data.frame(cbind(t=ONE.Data[[i]]$out[,1], N1=rowSums(ONE.Data[[i]]$out[, 2:(len+1), drop=F]),
                               N2=rowSums(ONE.Data[[i]]$out[,(len+2):(2*len+1), drop=F]),
                               R1=ONE.Data[[i]]$out[, (2*len+2)],R2=ONE.Data[[i]]$out[, (2*len+3)]))
  tempData$type <- if (cases[i,"VM"]==TRUE) "Constant Variation" else if (cases[i,"Va"]==FALSE) "No Variation" else "Heritable Variation"
  tempData$HL <- if (cases[i,"HL"]==TRUE) "Habitat Loss" else "No Habitat Loss"
  DataONE <- rbind(DataONE,tempData)
}
DataONE$type <- factor(DataONE$type,levels=c("No Variation","Heritable Variation","Constant Variation"))
DataONE$HL <- factor(DataONE$HL, levels = c("No Habitat Loss", "Habitat Loss"))

# high a0 value
DataNE <- data.frame()
for (i in c(1:nrow(cases))) { #comparison in presence of HL
  len <- nrow(NE.Data[[i]]$params)
  tempData <- data.frame(cbind(t=NE.Data[[i]]$out[,1], N1=rowSums(NE.Data[[i]]$out[, 2:(len+1), drop=F]),
                               N2=rowSums(NE.Data[[i]]$out[,(len+2):(2*len+1), drop=F]),
                               R1=NE.Data[[i]]$out[, (2*len+2)],R2=NE.Data[[i]]$out[, (2*len+3)]))
  tempData$type <- if (cases[i,"VM"]==TRUE) "Constant Variation" else if (cases[i,"Va"]==FALSE) "No Variation" else "Heritable Variation"
  tempData$HL <- if (cases[i,"HL"]==TRUE) "Habitat Loss" else "No Habitat Loss"
  DataNE <- rbind(DataNE,tempData)
}
DataNE$type <- factor(DataNE$type,levels=c("No Variation","Heritable Variation","Constant Variation"))
DataNE$HL <- factor(DataNE$HL, levels = c("No Habitat Loss", "Habitat Loss"))

Trait.Data.NHL.NE <- NE.Data[[5]]
Trait.Data.HL.NE <- NE.Data[[3]]
Trait.Data.NHL.ONE <- ONE.Data[[5]]
Trait.Data.HL.ONE <- ONE.Data[[3]]
Trait.Data.NHL.OE <- OE.Data[[5]]
Trait.Data.HL.OE <- OE.Data[[3]]

rm("NE.Data","ONE.Data","OE.Data","tempData")

################# OE Plots variation comparison ######################
p.OE <- ggplot(data=DataOE,aes(x=t)) + xlab("Time") + 
  scale_linetype_manual("",values = c("No Variation"="solid", "Heritable Variation"="dotted",
                                      "Constant Variation"="dashed")) +
  scale_alpha_manual(values = c("No Variation"=0.7, "Heritable Variation"=1,
                                "Constant Variation"=1),guide = "none") +
  scale_color_manual("",values=c("No Habitat Loss"="blue","Habitat Loss"="red")) +
  theme_bw() + theme(plot.title = element_text(hjust = 0.5, size = 20),
                     # axis.title.x = element_text(size = 15),
                     # axis.title.y = element_text(size = 8),
                     # axis.text = element_text(size = 7, face = "bold"),
                     # legend.title = element_text(size = 10),
                     # legend.text = element_text(size = 7),
                     legend.position="bottom",
                     # strip.text = element_text(size = 12),
                     plot.margin = margin(0.5, 0.5, 0.5, 0.5, "lines"),
                     panel.grid.major = element_blank(),panel.grid.minor = element_blank(),
                     strip.background = element_rect(colour="black", fill="white"),
                     panel.border = element_rect(colour = "black"),
                     panel.spacing = unit(0, "lines")) +
  scale_x_continuous(expand = c(0,0))# + scale_y_continuous(expand = c(0,0),limits = c(0,10))

pN1.OE <- p.OE + geom_line(data=DataOE,aes(y=N1, color=HL, linetype=type, alpha=type), linewidth=0.5) + 
  ylab("Consumer Density N1") + scale_y_continuous(expand = c(0,0),limits = c(-0.01,20))
#pN1.OE

pN2.OE <- p.OE + geom_line(data=DataOE,aes(y=N2, color=HL, linetype=type, alpha=type), linewidth=0.5) + 
  ylab("Consumer Density N2") + scale_y_continuous(expand = c(0,0),limits = c(-0.01,20))
#pN2.OE

################# ONE Plots variation comparison ######################
pN1.ONE <- p.OE + geom_line(data=DataONE,aes(y=N1, color=HL, linetype=type, alpha=type), linewidth=0.5) + 
  ylab("Consumer Density N1") + scale_y_continuous(expand = c(0,0),limits = c(-0.01,20))
#pN1.ONE

pN2.ONE <- p.OE + geom_line(data=DataONE,aes(y=N2, color=HL, linetype=type, alpha=type), linewidth=0.5) + 
  ylab("Consumer Density N2") + scale_y_continuous(expand = c(0,0),limits = c(-0.01,20))
#pN2.ONE

################# NE Plots variation comparison ######################
pN1.NE <- p.OE + geom_line(data=DataNE,aes(y=N1, color=HL, linetype=type, alpha=type), linewidth=0.5) + 
  ylab("Consumer Density N1") + scale_y_continuous(expand = c(0,0),limits = c(-0.01,20))
#pN1.NE

pN2.NE <- p.OE + geom_line(data=DataNE,aes(y=N2, color=HL, linetype=type, alpha=type), linewidth=0.5) + 
  ylab("Consumer Density N2") + scale_y_continuous(expand = c(0,0),limits = c(-0.01,20))
#pN2.NE

############## Landscape level density plots ###########################
pNL.OE <- p.OE + geom_line(data=DataOE,aes(y=N1+N2, color=HL, linetype=type, alpha=type), linewidth=0.5) +
  ylab("Total Consumer Density") + scale_y_continuous(expand = c(0,0),limits = c(0,25))
#pNL.OE

pNL.ONE <- p.OE + geom_line(data=DataONE,aes(y=N1+N2, color=HL, linetype=type, alpha=type), linewidth=0.5) +
  ylab("Total Consumer Density") + scale_y_continuous(expand = c(0,0),limits = c(0,25))
#pNL.ONE

pNL.NE <- p.OE + geom_line(data=DataNE,aes(y=N1+N2, color=HL, linetype=type, alpha=type), linewidth=0.5) +
  ylab("Total Consumer Density") + scale_y_continuous(expand = c(0,0),limits = c(0,25))
#pNL.NE

#################### Trait evolution plot ########################
times <- Trait.Data.NHL.NE$out[,1]
grain <- as.integer((length(times)-1)/tail(times,n=1))*5
out.NHL.NE <- Trait.Data.NHL.NE$out[seq(1,(length(times)),grain),]
out.HL.NE <- Trait.Data.HL.NE$out[seq(1,(length(times)),grain),]
out.NHL.ONE <- Trait.Data.NHL.ONE$out[seq(1,(length(times)),grain),]
out.HL.ONE <- Trait.Data.HL.ONE$out[seq(1,(length(times)),grain),]
out.NHL.OE <- Trait.Data.NHL.OE$out[seq(1,(length(times)),grain),]
out.HL.OE <- Trait.Data.HL.OE$out[seq(1,(length(times)),grain),]
p <- Trait.Data.NHL.NE$params
len <- length(p[,1])
times <- seq(0,tail(times,n=1),5)
rm("Trait.Data.NHL.NE","Trait.Data.HL.NE","Trait.Data.NHL.OE","Trait.Data.HL.OE","Trait.Data.NHL.ONE","Trait.Data.HL.ONE")

trait.dist.fun <- function(data,trait,p,min.quant,max.quant,HLtype,a0val) {
  trait.dist.data <- data.frame(Time=data[,1])
  
  out.df.1 <- trait.dist.data
  out.df.1$trait.mean <- rowSums(t(t((data[,2:(len+1)])/rowSums(data[,2:(len+1)]))*p[,trait]))
  out.df.1$trait.min <- as.numeric(apply(data[,2:(len+1)]/rowSums(data[,2:(len+1)]), 1, function(x) p[,trait][max(which(cumsum(x)<=min.quant))]))
  out.df.1$trait.max <- as.numeric(apply(data[,2:(len+1)]/rowSums(data[,2:(len+1)]), 1, function(x) p[,trait][min(which(cumsum(x)>=max.quant))]))
  out.df.1$density <- rowSums(data[, 2:(len+1), drop=F])
  out.df.1$consumer <- "N1"
  
  out.df.2 <- trait.dist.data
  out.df.2$trait.mean <- rowSums(t(t((data[,(len+2):(2*len+1)])/rowSums(data[,(len+2):(2*len+1)]))*p[,trait]))
  out.df.2$trait.min <- as.numeric(apply(data[,(len+2):(2*len+1)]/rowSums(data[,(len+2):(2*len+1)]), 1, function(x) p[,trait][max(which(cumsum(x)<=min.quant))]))
  out.df.2$trait.max <- as.numeric(apply(data[,(len+2):(2*len+1)]/rowSums(data[,(len+2):(2*len+1)]), 1, function(x) p[,trait][min(which(cumsum(x)>=max.quant))]))
  out.df.2$density <- rowSums(data[,(len+2):(2*len+1), drop=F])
  out.df.2$consumer <- "N2"
  
  trait.dist.data <- rbind(out.df.1,out.df.2)
  trait.dist.data[,"HL"] <- HLtype
  trait.dist.data[,"a0"] <- a0val
  return(trait.dist.data)
}

trait.NHL.NE <- trait.dist.fun(out.NHL.NE,trait,p,0.05,0.95,"No Habitat Loss","NE")
trait.HL.NE <- trait.dist.fun(out.HL.NE,trait,p,0.05,0.95,"Habitat Loss","NE")
trait.NHL.ONE <- trait.dist.fun(out.NHL.ONE,trait,p,0.05,0.95,"No Habitat Loss","ONE")
trait.HL.ONE <- trait.dist.fun(out.HL.ONE,trait,p,0.05,0.95,"Habitat Loss","ONE")
trait.NHL.OE <- trait.dist.fun(out.NHL.OE,trait,p,0.05,0.95,"No Habitat Loss","OE")
trait.HL.OE <- trait.dist.fun(out.HL.OE,trait,p,0.05,0.95,"Habitat Loss","OE")

trait.dist.data <- rbind(trait.NHL.NE,trait.HL.NE,trait.NHL.ONE,trait.HL.ONE,
                         trait.NHL.OE,trait.HL.OE)
#levels(trait.dist.data$consumer) <- c(N1="$N_1$", N2="$N_2$")
rm("out.NHL.NE","out.HL.NE","out.NHL.OE","out.HL.OE","out.NHL.ONE","out.HL.ONE",
   "trait.NHL.NE","trait.HL.NE","trait.NHL.ONE","trait.HL.ONE","trait.NHL.OE","trait.HL.OE")

# generate trait distributions over time
consumer_names <- list(N1=TeX("$N_1$"),N2=TeX("$N_2$")) #For latex in facet strip text
consumer_labeller <- function(variable,value){
  return(consumer_names[value])
}
#Common ggplot object
p.trait <- ggplot(trait.dist.data,aes(x=Time, alpha=density, col=HL)) +
  facet_grid(consumer~.,labeller = consumer_labeller) + xlab("Time") + ylab(trait) +
  scale_color_manual("",values=c("No Habitat Loss"="blue","Habitat Loss"="red")) +
  #scale_fill_manual("",values=c("No Habitat Loss"="blue","Habitat Loss"="red")) +
  #scale_linetype_manual("",values=c("N1"="solid","N2"="dotdash")) +
  #scale_color_gradient(low="white",high="black") +
  scale_alpha_continuous(range=c(0,0.7)) + ylim(c(0,1)) +
  theme_bw() + theme(panel.grid.major = element_blank(),
                     panel.grid.minor = element_blank(),
                     strip.background = element_rect(colour="black", fill="white"),
                     panel.spacing.y = unit(c(0.25), "lines"),
                     axis.text.x = element_text(angle=90, vjust=0.5),
                     plot.title = element_text(hjust = 0.5, size = 20),
                     # axis.title.x = element_text(size = 15),
                     # axis.title.y = element_text(size = 8),
                     # axis.text = element_text(size = 7, face = "bold"),
                     # legend.title = element_text(size = 10),
                     # legend.text = element_text(size = 7),
                     # strip.text = element_text(size = 12),
                     plot.margin = margin(0.5, 0.5, 0.5, 0.5, "lines"),
                     panel.border = element_rect(colour = "black")) +
  scale_x_continuous(expand = c(0,0), guide = guide_axis(check.overlap = T))

p.NE.trait <- p.trait + geom_line(data=trait.dist.data[trait.dist.data$a0=="NE",],aes(y=trait.mean), linewidth=0.5) +
  geom_line(data=trait.dist.data[trait.dist.data$a0=="NE",],aes(y=trait.min), linewidth=0.25) +
  geom_line(data=trait.dist.data[trait.dist.data$a0=="NE",],aes(y=trait.max), linewidth=0.25)
#p.NE.trait
p.ONE.trait <- p.trait + geom_line(data=trait.dist.data[trait.dist.data$a0=="ONE",],aes(y=trait.mean), linewidth=0.5) +
  geom_line(data=trait.dist.data[trait.dist.data$a0=="ONE",],aes(y=trait.min), linewidth=0.25) +
  geom_line(data=trait.dist.data[trait.dist.data$a0=="ONE",],aes(y=trait.max), linewidth=0.25)
#p.NE.trait
p.OE.trait <- p.trait + geom_line(data=trait.dist.data[trait.dist.data$a0=="OE",],aes(y=trait.mean), linewidth=0.5) +
  geom_line(data=trait.dist.data[trait.dist.data$a0=="OE",],aes(y=trait.min), linewidth=0.25) +
  geom_line(data=trait.dist.data[trait.dist.data$a0=="OE",],aes(y=trait.max), linewidth=0.25)
#p.OE.trait

################### Making Final Plot ##########################
pobj <- ggarrange(ggarrange(pN1.OE+theme(plot.title = element_text(hjust = 0.5,size = 10),axis.title.x = element_blank(),axis.ticks.x = element_blank(),axis.text.x = element_blank(), axis.title.y = element_blank())+ggtitle(TeX("$N_1$"))+ylab("Consumer Density"),
                      pN2.OE+theme(plot.title = element_text(hjust = 0.5,size = 10),axis.title.x = element_blank(),axis.ticks.x = element_blank(),axis.text.x = element_blank(), axis.title.y = element_blank())+ggtitle(TeX("$N_2$")),
                      pNL.OE+theme(plot.title = element_text(hjust = 0.5,size = 10),axis.title.x = element_blank(),axis.ticks.x = element_blank(),axis.text.x = element_blank(), axis.title.y = element_blank())+ggtitle(TeX("Landscape ($N_1 + N_2$)")),
                      pN1.ONE+theme(plot.title = element_blank(),axis.title.x = element_blank(),axis.ticks.x = element_blank(),axis.text.x = element_blank(), axis.title.y = element_blank())+ylab("Consumer Density"),
                      pN2.ONE+theme(plot.title = element_blank(),axis.title.x = element_blank(),axis.ticks.x = element_blank(),axis.text.x = element_blank(), axis.title.y = element_blank()),
                      pNL.ONE+theme(plot.title = element_blank(),axis.title.x = element_blank(),axis.ticks.x = element_blank(),axis.text.x = element_blank(), axis.title.y = element_blank()),
                      pN1.NE+theme(plot.title = element_blank(),axis.text.x = element_text(angle = 90, vjust = 0.5),axis.title.x = element_blank(), axis.title.y = element_blank())+ylab("Consumer Density"),
                      pN2.NE+theme(plot.title = element_blank(),axis.text.x = element_text(angle = 90, vjust = 0.5),axis.title.x = element_blank(), axis.title.y = element_blank()),
                      pNL.NE+theme(plot.title = element_blank(),axis.text.x = element_text(angle = 90, vjust = 0.5),axis.title.x = element_blank(), axis.title.y = element_blank()),
                      labels = c("a","b","c","e","f","g","i","j","k"),
                      heights = c(0.96,0.85,1),
                      ncol = 3, nrow = 3, align = "v", vjust = 1.1, hjust = 0.3,
                      common.legend = TRUE, legend = "none"),
                  ggarrange(p.OE.trait+theme(plot.title = element_text(hjust = 0.5,size = 10),axis.title.x = element_blank(),axis.ticks.x = element_blank(),axis.text.x = element_blank(),axis.text.y = element_text(size = 7))+
                              ylab("p")+ggtitle("Trait Evolution"),
                            p.ONE.trait+theme(plot.title = element_blank(),axis.title.x = element_blank(),axis.ticks.x = element_blank(),axis.text.x = element_blank(),axis.text.y = element_text(size = 7))+
                              ylab("p"),
                            p.NE.trait+theme(plot.title = element_blank(),axis.title.x = element_blank(),axis.text.y = element_text(size = 7))+
                              ylab("p"),
                            nrow=3, ncol = 1, labels = c("d","h","l"),
                            align = "v", vjust = 1.1, heights = c(0.96,0.85,1),
                            common.legend = TRUE, legend = "none"),
                  ncol=2,widths = c(1,0.35))

pobj <- annotate_figure(pobj, bottom = text_grob("Time", size = 10))
pobj <- annotate_figure(pobj, bottom = get_legend(pN1.OE))
pobj <- annotate_figure(pobj, left = text_grob("Consumer Density", rot = 90, hjust = 0.3))
pobj <- annotate_figure(pobj, left = text_grob(TeX("$a_0 = 0.18$                   $a_0 = 0.15$                    $a_0 = 0.1$"), rot = 90, face = "bold", hjust=0.4))
#pobj
ggsave(filename = filname, device = "pdf", plot = pobj, width = 9, height = 6)
```
