## Supplementary material for "Multi-scale Effects of Habitat Loss and the Role of Trait Variation": R code for generating data and figures: SI_Figure_S1_S2_plotfile.R

```
# HL shape explanation figure
rm(list = ls())
library(ggplot2)
library(ggpubr)
library(latex2exp) #useful to use latex symbols
library(reshape2) #for reshaping dataframes
loadRData <- function(fileName){ #loads an RData file, and returns it
  load(fileName)
  get(ls()[ls() != "fileName"])
}

###################### Preppeing Data ########################
filname1 <- "SI_Figure_S1.pdf"
filname2 <- "SI_Figure_S2.pdf"

parsweep.curator <- function(filepath, trait) {
  obj <- loadRData(filepath)
  
  cases <- expand.grid(Va=c(T,F),HL=c(T,F),VM=c(T,F))
  cases <- cases[!(cases[,"Va"]==F & cases[,"VM"]==T),]
  
  Data <- data.frame()
  for (j in 1:length(obj)) {
    for (i in 1:nrow(cases)) {
      obj1<-obj[[j]]
      len <- nrow(obj1[[i]]$params)
      tempData <- data.frame(cbind(t=obj1[[i]]$out[,1],N1=rowSums(obj1[[i]]$out[, 2:(len+1), drop=F]),
                                   N2=rowSums(obj1[[i]]$out[,(len+2):(2*len+1), drop=F]),
                                   R1=rowSums(obj1[[i]]$out[, (2*len+2), drop=F]),
                                   R2=rowSums(obj1[[i]]$out[, (2*len+3), drop=F])))
      tempData$type <- if (cases[i,"VM"]==TRUE) "Constant Variation" else if (cases[i,"Va"]==FALSE) "No Variation" else "Heritable Variation"
      tempData$HL <- if (cases[i,"HL"]==TRUE) "Habitat Loss" else "No Habitat Loss"
      tempData$param_val <- as.numeric(names(obj)[j])
      Data <- rbind(Data,tempData)
    }
  }
  Data$type <- factor(Data$type,levels=c("No Variation","Heritable Variation","Constant Variation"))
  Data$HL <- factor(Data$HL, levels = c("No Habitat Loss", "Habitat Loss"))
  Data$Trait <- trait
  Data$Trait <- factor(Data$Trait)
  return(Data)
}
data.curator <- function(filepath, trait) {
  obj <- loadRData(filepath)
  
  cases <- expand.grid(Va=c(T,F),HL=c(T,F),VM=c(T,F))
  cases <- cases[!(cases[,"Va"]==F & cases[,"VM"]==T),]
  
  Data <- data.frame()
  obj1<-obj
    for (i in 1:nrow(cases)) {
      len <- nrow(obj1[[i]]$params)
      tempData <- data.frame(cbind(t=obj1[[i]]$out[,1],N1=rowSums(obj1[[i]]$out[, 2:(len+1), drop=F]),
                                   N2=rowSums(obj1[[i]]$out[,(len+2):(2*len+1), drop=F]),
                                   R1=rowSums(obj1[[i]]$out[, (2*len+2), drop=F]),
                                   R2=rowSums(obj1[[i]]$out[, (2*len+3), drop=F])))
      tempData$type <- if (cases[i,"VM"]==TRUE) "Constant Variation" else if (cases[i,"Va"]==FALSE) "No Variation" else "Heritable Variation"
      tempData$HL <- if (cases[i,"HL"]==TRUE) "Habitat Loss" else "No Habitat Loss"
      tempData$param_val <- obj1[[1]][["params"]][1,trait]
      Data <- rbind(Data,tempData)
    }
  Data$type <- factor(Data$type,levels=c("No Variation","Heritable Variation","Constant Variation"))
  Data$HL <- factor(Data$HL, levels = c("No Habitat Loss", "Habitat Loss"))
  Data$Trait <- trait
  Data$Trait <- factor(Data$Trait)
  return(Data)
}

Data.t0 <- parsweep.curator("Data/e-var-mean_0p2-sd_0p08-parsweep_t0/t0/rawdata/file.RData","t0")
Data.t0.def <- data.curator("Data/e-var-mean_0p2-sd_0p08-a0_0p1/rawdata/file.RData","t0")
Data.k <- parsweep.curator("Data/e-var-mean_0p2-sd_0p08-parsweep_k/k/rawdata/file.RData","k")
Data.k.def <- data.curator("Data/e-var-mean_0p2-sd_0p08-a0_0p1/rawdata/file.RData","k")

Data <- rbind(Data.t0,Data.t0.def,Data.k,Data.k.def)
#levels(Data$Trait)<-list("t[1/2]"="t0","s"="k")
rm(Data.t0,Data.t0.def,Data.k,Data.k.def)

Data <- Data[Data$param_val==c(0,3500,6000,0.001,0.003,0.061),]
Data <- Data[Data$HL=="Habitat Loss",]
for (i in c(0,3500,6000,0.001,0.003,0.061)){
  Data[Data$param_val==i,"param_val"] <- rep(c("low","default","high"),2)[which(c(0,3500,6000,0.001,0.003,0.061)==i)]
}
Data$param_val <- factor(Data$param_val, levels = c("low","default","high"))

HLshape.data <- data.frame()
for (i in 1:length(c(0,3500,6000))){
  tempdata <- data.frame(cbind(t=as.numeric(0:max(Data$t)),
              rate=as.numeric(0.5/(1+exp(-0.003*(0:max(Data$t)-c(0,3500,6000)[i]))))))
  tempdata$par <- "t0"
  tempdata$parval=c("low","default","high")[i]
  HLshape.data <-rbind(HLshape.data,tempdata)
}
for (i in 1:length(c(0.001,0.003,0.061))){
  tempdata <- data.frame(cbind(t=as.numeric(0:max(Data$t)),
                               rate=as.numeric(0.5/(1+exp(-c(0.001,0.003,0.061)[i]*(0:max(Data$t)-3500))))))
  tempdata$par <- "k"
  tempdata$parval=c("low","default","high")[i]
  HLshape.data <-rbind(HLshape.data,tempdata)
}
HLshape.data$par <- factor(HLshape.data$par)
levels(HLshape.data$par)<-list("t[1/2]"="t0","s"="k")
HLshape.data$parval <- factor(HLshape.data$parval, levels = c("low","default","high"))

############# plotting the figure depicting HL shapes ################
pshape <- ggplot(data = HLshape.data, aes(x=t)) +
  scale_colour_manual("Parameter value",
                      values = c("low"="red", "default"="blue", "high"="green")) +
  theme_bw() + theme(plot.title = element_text(hjust = 0.5),
                     # axis.title.x = element_text(size = 15),
                     #axis.title.y = element_text(size = 15),
                     axis.text.x = element_text(angle=90, vjust=0.5),
                     #legend.title = element_text(size = 10),
                     #legend.text = element_text(size = 7),
                     legend.position="bottom",
                     # strip.text = element_text(size = 12),
                     #plot.margin = margin(0.5, 1, 0.5, 0.5, "lines"),
                     panel.grid.major = element_blank(),panel.grid.minor = element_blank(),
                     strip.background = element_rect(colour="black", fill="white"),
                     strip.placement = "outside",
                     panel.border = element_rect(colour = "black"),
                     panel.spacing = unit(0.7, "lines")) +
  scale_x_continuous(expand = c(0,0))
pshape.fig <- pshape + geom_line(aes(y=rate, color=parval), linewidth=0.5) + 
  facet_grid(.~par,scales = "free",labeller = label_parsed)+ xlab("Time") + ylab("Resource removal rate") +
  scale_y_continuous(expand = c(0.01,0))
#pshape.fig

ggsave(filename = filname1, device = "pdf", plot = pshape.fig, width = 8, height = 4)

############## plotting figures depicting time series plots ################
pobj <- ggplot(data=Data,aes(x=t)) + xlab("Time") +
  scale_colour_manual("Parameter value",
                      values = c("low"="red", "default"="blue", "high"="green")) +
  scale_linetype_manual("Patch",values=c("1"="solid","2"="dotted")) +
  theme_bw() + theme(plot.title = element_text(hjust = 0.5),
                     # axis.title.x = element_text(size = 15),
                     #axis.title.y = element_text(size = 15),
                     axis.text.x = element_text(angle=90, vjust=0.5),
                     #legend.title = element_text(size = 10),
                     #legend.text = element_text(size = 7),
                     legend.position="bottom",
                     # strip.text = element_text(size = 12),
                     #plot.margin = margin(0.5, 1, 0.5, 0.5, "lines"),
                     panel.grid.major = element_blank(),panel.grid.minor = element_blank(),
                     strip.background = element_rect(colour="black", fill="white"),
                     strip.placement = "outside",
                     panel.border = element_rect(colour = "black"),
                     panel.spacing = unit(0.7, "lines")) +
  scale_x_continuous(expand = c(0,0)) + scale_y_continuous(expand = c(0.01,0))

ts.fig <- pobj + geom_line(aes(y=N1, color=param_val, linetype="1"), linewidth=0.5, alpha=0.7) + 
  geom_line(aes(y=N2, color=param_val, linetype="2"), linewidth=0.5) +
  facet_grid(Trait~type,scales = "free",labeller=labeller(Trait = as_labeller(c(`t0`="t[1/2]",`k`="s"), label_parsed)))+
  ylab("Consumer Density when HL")
#ts.fig

ggsave(filename = filname2, device = "pdf", plot = ts.fig, width = 8, height = 6)
```
