## Supplementary material for "Multi-scale Effects of Habitat Loss and the Role of Trait Variation": R code for generating data and figures: SI_Figure_S3_plotfile.R

```
#Over-consumption explanation plot in SI

rm(list = ls())
library(ggplot2)
library(ggpubr)

filname <- "SI_Figure_S3.pdf" # plot saved in this file

# function to calculate N1*
neq.fun <- function(C0,C1,a0,e,r) {
  nvec <- (C0/r)*(1 - ((2*a0)/(e*r*C1)))
  nvec[nvec<0] <- NA
  return(nvec)
}
C0 <- 0.5
C1 <- 50
a0 <- 0.1
a0_high <- 0.14
e <- 0.2
e_low <- 0.14
b0 <- 30
b0_low <- 25
b1 <- 500
rvec <- seq(0,0.301,0.001)
data <- data.frame(r = rvec)
data$default <- neq.fun(C0,C1,a0,e,rvec)
data$high_a0 <- neq.fun(C0,C1,a0_high,e,rvec)
data$low_b0 <- neq.fun(C0,C1,a0,e,rvec*(b0_low/b0))
data$low_e <- neq.fun(C0,C1,a0,e_low,rvec)


pobj <- ggplot(data=data,aes(x=r)) + xlab("r") +
  scale_color_manual("",values=c("Habitat Loss"="red","No Habitat Loss"="blue")) +
  scale_linetype_manual("Scenario",values = c("Default"="solid","Other"="dashed")) +
  theme_bw() + theme(plot.title = element_text(hjust = 0.5),
                     axis.title.x = element_text(face = "bold"),
                     axis.title.y = element_text(size=9,face = "bold"),
                     axis.text.x = element_text(angle=90, vjust=0.5),
                     #legend.title = element_text(size = 10),
                     #legend.text = element_text(size = 7),
                     legend.position="bottom",
                     # strip.text = element_text(size = 12),
                     plot.margin = margin(0.5, 1, 0.5, 0.5, "lines"),
                     panel.grid.major = element_blank(),panel.grid.minor = element_blank(),
                     strip.background = element_blank(),
                     strip.placement = "outside",
                     panel.border = element_rect(colour = "black"),
                     panel.spacing = unit(0.7, "lines")) +
  scale_x_continuous(expand = c(0,0)) + scale_y_continuous(expand = c(0,0),limits = c(0,7))

pN1.OE <- pobj + geom_line(aes(y=default, linetype="Default"), linewidth=0.5, na.rm=TRUE) +
  geom_line(aes(y=high_a0, linetype="Other"), linewidth=0.5, na.rm=TRUE) +
  geom_vline(aes(xintercept=(b0)/(b1), color="No Habitat Loss"), linewidth=0.3) +
  geom_vline(aes(xintercept=(b0*0.7)/(b1), color="Habitat Loss"), linewidth=0.3) +
  ylab(expression("N"[1]*"*"))
#pN1.OE

pN2.OE <- pobj + geom_line(aes(y=default, linetype="Default"), linewidth=0.5, na.rm=TRUE) +
  geom_line(aes(y=low_b0, linetype="Other"), linewidth=0.5, na.rm=TRUE) +
  geom_vline(aes(xintercept=(b0*0.85)/(b1), color="No Habitat Loss"), linewidth=0.3) +
  geom_vline(aes(xintercept=(b0*0.85*0.7)/(b1), color="Habitat Loss"), linewidth=0.3) +
  ylab(expression("N"[1]*"*"))
#pN2.OE

pN.OE <- pobj + geom_line(aes(y=default, linetype="Default"), linewidth=0.5, na.rm=TRUE) +
  geom_line(aes(y=low_e, linetype="Other"), linewidth=0.5, na.rm=TRUE) +
  geom_vline(aes(xintercept=(b0)/(b1), color="No Habitat Loss"), linewidth=0.3) +
  geom_vline(aes(xintercept=(b0*0.7)/(b1), color="Habitat Loss"), linewidth=0.3) +
  ylab(expression("N"[1]*"*"))
#pN.OE

################### Making Final Plot ##########################
figure <- ggarrange(pN1.OE+theme(axis.title.x = element_blank(),axis.ticks.x = element_blank(),axis.text.x = element_blank(),plot.title = element_blank()),
                    pN2.OE+theme(axis.title.x = element_blank(),axis.ticks.x = element_blank(),axis.text.x = element_blank(),plot.title = element_blank()),
                    pN.OE+theme(plot.title = element_blank()),
                    labels = "auto", heights = c(0.85,0.85,1),
                    ncol = 1, nrow = 3, #align = "hv", vjust = 1,# hjust = -1,
                    common.legend = TRUE, legend = "bottom")

ggsave(filename = filname, device = "pdf", plot = figure, width = 5, height = 6)
```
