## Supplementary material for "Multi-scale Effects of Habitat Loss and the Role of Trait Variation": R code for generating data and figures: SI_Figure_S4_plotfile.R

```
# We compare equilibrium values over parameter space with no variation case
rm(list = ls())
library(ggplot2)
library(ggpubr)
library(latex2exp) #useful to use latex symbols
library(reshape2) #for reshaping dataframes
loadRData <- function(fileName){ #loads an RData file, and returns it
  load(fileName)
  get(ls()[ls() != "fileName"])
}

###################### Preppeing Data ########################
filname <- "SI_Figure_S4.pdf"

parsweep.curator <- function(filepath, trait) {
  obj <- loadRData(filepath)
  
  cases <- expand.grid(Va=c(T,F),HL=c(T,F),VM=c(T,F))
  cases <- cases[!(cases[,"Va"]==F & cases[,"VM"]==T),]
  
  Data <- data.frame()
  for (j in 1:length(obj)) {
    for (i in 1:nrow(cases)) {
      obj1<-obj[[j]]
      len <- nrow(obj1[[i]]$params)
      tempData <- data.frame(cbind(N1=mean(tail(rowSums(obj1[[i]]$out[, 2:(len+1), drop=F]),n=1)),
                                   N2=mean(tail(rowSums(obj1[[i]]$out[,(len+2):(2*len+1), drop=F]),n=1)),
                                   R1=mean(tail(obj1[[i]]$out[, (2*len+2)],n=1)),
                                   R2=mean(tail(obj1[[i]]$out[, (2*len+3)],n=1))))
      tempData$type <- if (cases[i,"VM"]==TRUE) "Constant Variation" else if (cases[i,"Va"]==FALSE) "No Variation" else "Heritable Variation"
      tempData$HL <- if (cases[i,"HL"]==TRUE) "Habitat Loss" else "No Habitat Loss"
      tempData$param_val <- as.numeric(names(obj)[j])
      Data <- rbind(Data,tempData)
    }
  }
  Data$type <- factor(Data$type,levels=c("No Variation","Heritable Variation","Constant Variation"))
  Data$HL <- factor(Data$HL, levels = c("No Habitat Loss", "Habitat Loss"))
  Data$Trait <- trait
  Data$Trait <- factor(Data$Trait)
  Data <- melt(Data, id.vars=c("R1","R2","type","HL","param_val","Trait"),
                 variable.name = "patch", value.name="density")
  Data$patch <- factor(Data$patch)
  return(Data)
}

Data.a0 <- parsweep.curator("Data/e-var-mean_0p2-sd_0p08-parsweep_a0/a0/rawdata/file.RData","a0")
Data.b0 <- parsweep.curator("Data/e-var-mean_0p2-sd_0p08-parsweep_b0/b0/rawdata/file.RData","b0")
Data.e <- parsweep.curator("Data/e-var-mean_0p2-sd_0p08-parsweep_e/ttmean/rawdata/file.RData","e")
Data.p <- parsweep.curator("Data/e-var-mean_0p2-sd_0p08-parsweep_p/p1/rawdata/file.RData","p")
Data.beta <- parsweep.curator("Data/e-var-mean_0p2-sd_0p08-parsweep_beta/beta/rawdata/file.RData","beta")
Data.d <- parsweep.curator("Data/e-var-mean_0p2-sd_0p08-parsweep_b1/b1/rawdata/file.RData","b1")

Data <- rbind(Data.a0,Data.b0,Data.e,Data.p,Data.beta,Data.d)
Data <- Data[Data$type=="No Variation",]
levels(Data$patch)<-list("1"="N1","2"="N2")
rm(Data.a0,Data.b0,Data.e,Data.p,Data.beta,Data.d)
pobj <- ggplot(data=Data,aes(x=param_val)) + 
  scale_linetype_manual("Patch",values = c("1"="solid", "2"="dotted")) +
  scale_alpha_manual(values = c("1"=0.7, "2"=1),guide = "none") +
  scale_size_manual(values = c("1"=0.5, "2"=0.7),guide = "none") +
  scale_color_manual("",values=c("No Habitat Loss"="blue","Habitat Loss"="red")) +
  theme_bw() + theme(plot.title = element_text(hjust = 0.5),
                     axis.title.x = element_text(face = "bold"),
                     axis.title.y = element_blank(),
                     axis.text.x = element_text(angle=90, vjust=0.5),
                     #legend.title = element_text(size = 10),
                     #legend.text = element_text(size = 7),
                     legend.position="bottom",
                     # strip.text = element_text(size = 12),
                     plot.margin = margin(0.5, 1, 0.5, 0.5, "lines"),
                     panel.grid.major = element_blank(),panel.grid.minor = element_blank(),
                     strip.background = element_blank(),
                     strip.placement = "outside",
                     panel.border = element_rect(colour = "black"),
                     panel.spacing = unit(0.7, "lines")) +
  scale_x_continuous(expand = c(0,0.0001))# + scale_y_continuous(expand = c(0,0),limits = c(0,10))

p.a0 <- pobj + geom_line(data=Data[Data$Trait=="a0",],aes(y=density, color=HL, linetype=patch, alpha=patch, size=patch)) +
  scale_y_continuous(expand = c(0.01,0.01)) + xlab(TeX("$a_0$"))

p.b0 <- pobj + geom_line(data=Data[Data$Trait=="b0",],aes(y=density, color=HL, linetype=patch, alpha=patch, size=patch)) +
  scale_y_continuous(expand = c(0.01,0.01)) + xlab(TeX("$b_0$"))

p.e <- pobj + geom_line(data=Data[Data$Trait=="e",],aes(y=density, color=HL, linetype=patch, alpha=patch, size=patch)) +
  scale_y_continuous(expand = c(0.01,0.01)) + xlab(TeX("$\\epsilon$"))

p.p <- pobj + geom_line(data=Data[Data$Trait=="p",],aes(y=density, color=HL, linetype=patch, alpha=patch, size=patch)) +
  scale_y_continuous(expand = c(0.01,0.01)) + xlab(TeX("$p$"))

p.beta <- pobj + geom_line(data=Data[Data$Trait=="beta",],aes(y=density, color=HL, linetype=patch, alpha=patch, size=patch)) +
  scale_y_continuous(expand = c(0.01,0.01)) + xlab(TeX("$\\beta$"))

p.D <- pobj + geom_line(data=Data[Data$Trait=="b1",],aes(y=density, color=HL, linetype=patch, alpha=patch, size=patch)) +
  scale_y_continuous(expand = c(0.01,0.01)) + xlab(TeX("$b_1$"))

figure <- ggarrange(p.a0+theme(plot.title = element_blank(), axis.title.y = element_blank()),
                    p.b0+theme(plot.title = element_blank(), axis.title.y = element_blank()),
                    p.e+theme(plot.title = element_blank(), axis.title.y = element_blank()),
                    p.p+theme(plot.title = element_blank(), axis.title.y = element_blank()),
                    p.beta+theme(plot.title = element_blank(), axis.title.y = element_blank()),
                    p.D+theme(plot.title = element_blank(), axis.title.y = element_blank()),
                    labels = "auto",
                    ncol = 3, nrow = 2, align = "hv", vjust = 1.2, hjust = 0.2,
                    common.legend = TRUE, legend = "none")
figure <- annotate_figure(figure, left = text_grob("Equilibrium Consumer Density", rot = 90))
figure <- annotate_figure(figure, bottom = get_legend(p.a0))

ggsave(filename = filname, device = "pdf", plot = figure, width = 8, height = 5)

# figure <- pobj + geom_line(aes(y=density, color=HL, linetype=patch, alpha=patch), linewidth=0.5) + 
#   facet_grid(.~Trait,scales = "free",labeller = label_parsed, switch="both")+ xlab("Parameter Values") + ylab("Consumer Density") +
#   scale_y_continuous(expand = c(0.01,0.01))
```
