## Supplementary material for "Multi-scale Effects of Habitat Loss and the Role of Trait Variation": R code for generating data and figures: ReadMe.pdf

### Information about Files and Folders

- Figure_x.R are the files which generate data and plots for the figure x.
- SI_Figure_x.R are the files which generate data and plots for the figure x in SI.
- Bagpack.R, Bagpack.noPD, and MCfun_multcases.R are the background files, which contain all the user-defined functions.
- MCfun_multcases_test.R can be used for exploring the connected patches scenario and MCfun_multcases_test-noPD.R can be used for exploring the isolated patches scenario.
- All the files inside the plotcode folder are files that have the plotting functions. These files are called by the respective Figure_x.R or SI_Figure_x.R files.
- The data generated by the Figure_x.R and SI_Figure_x.R files will be stored automatically in newly generated folder called Data.

### About the MCfun_multcases function

Observe that all the Figure_x.R and SI_Figure_x.R file use the MCfun_multcases function to generate data. It simulates all the scenarios with respect to habitat loss and trait variation for a given set of parameter values. It can also be used to simulate a range of parameter values (parameter space) for one parameter at a time.

Important thing to note that the symbols used for parameters in the code are different from those used in the manuscript. Refer the following table for the same.

| Parameter in the Manuscript | Symbol in Manuscript | Symbol in R code |
|---|---|---|
| per-capita death rate | $a_0$ | a0 |
| maximum per-capita resource consumption rate | $b_0$ | b0 |
| half-saturation constant for resource consumption | $b_1$ | b1 |
| within-patch foraging preference | p | p1 |
| within-patch mating preference | β | beta |
| Allee effect parameter | Θ | theta |
| per-capita resource conversion efficiency | ε | e |
| cross-patch foraging efficiency relative to within-patch | $\varepsilon_c$ | ec |
| maximum growth rate of resources | $r_0$ | C01, C02 * |
| carrying capacity of resources | k | C11, C12 * |
| maximum rate of resource degradation or removal | D | d |
| steepness of logistic resource removal | s | k |
| time when the resource removal rate is D/2 | $t_{1/2}$ | t0 |
| mutation rate | μ | mu |
| number of bins when trait variation is present | B | bins |

*These two resource parameters can have different values for the two patches in our model. But for the purpose of this study C01=C02 and C11=C12

### Example uses of the MCfun_multcases function

- When **evolving trait is ε** and Data stored in folder "e-var-mean_0p2-sd_0p08-a0_0p1"
    - source("Bagpack.R")
      **foldername = "Data/e-var-mean_0p2-sd_0p08-a0_0p1"**
      **trait = c("e")**
      parsweep_trait = c()
      obj <- MCfun_multcases (a0 = 0.1, b0 = 30, b1 = 500, bins=21,
      p1 = 0.7, beta = 0.8, **vartrait = trait**, ToffMinMaxPos = c(0,1,TRUE),
      **ttmin=0, ttmax=0.4, ttmean=0.2, ttsd=0.08,**
      e = 0.2, ec = 0.9, C01 = 0.5, C02 = 0.5, C11 = 50, C12 =50,
      N01 = 3, N02 = 4, R01 = 10, R02 = 10, foldername = foldername,
      mu = 0.3, d = 0.5, k = 0.003, t0 = 3500, theta = 2, times = seq(0, 8000, 1),
      mutations = T, parasweep=parsweep_trait, GeneticMixing=T, Allee = T, HLatEQ = F,
      savedata = T)

- When **evolving trait is ε** and the code runs over parameter range of D. Note the input to d becomes a vector over which the parameter space is scanned.
    - **trait = c("e")**
      **parsweep_trait = c("d")**
      obj <- MCfun_multcases (a0 = 0.1, b0 = 30, b1 = 500, bins=21,
      p1 = 0.7, beta = 0.8, **vartrait = trait**, ToffMinMaxPos = c(0,1,TRUE),
      **ttmin=0, ttmax=0.4, ttmean=0.2, ttsd=0.08,**
      e = 0.2, ec = 0.9, C01 = 0.5, C02 = 0.5, C11 = 50, C12 =50,
      N01 = 3, N02 = 4, R01 = 10, R02 = 10, foldername = foldername,
      mu = 0.3, **d = seq(0,0.5,0.025)**, k = 0.003, t0 = 3500, theta = 2, times = seq(0, 8000, 1),
      mutations = T, parasweep=parsweep_trait, GeneticMixing=T, Allee = T, HLatEQ = F,
      savedata = T)

- When **evolving trait is coupled i.e. p=beta**. Notice that the two coupled traits go as an input to the **vartrait** argument. **ToffMinMaxPos = c(0,1,TRUE)** takes the minimum, maximum, and TRUE (if relationship between coupled traits is positive) or FALSE (if it is negative) as inputs. Minimum and maximum are for the trait values of the coupled trait.
    - **trait = c("p1","beta")**
      parsweep_trait = c()
      obj <- MCfun_multcases (a0 = 0.1, b0 = 30, b1 = 500, bins=21,
      p1 = 0.7, beta = 0.8, **vartrait = trait**, **ToffMinMaxPos = c(0,1,TRUE)**,
      **ttmin=0, ttmax=1, ttmean=0.65, ttsd=0.1,**
      e = 0.2, ec = 0.9, C01 = 0.5, C02 = 0.5, C11 = 50, C12 =50,
      N01 = 3, N02 = 4, R01 = 10, R02 = 10, foldername = foldername,
      mu = 0.3, d = 0.5, k = 0.003, t0 = 3500, theta = 2, times = seq(0, 8000, 1),
      mutations = T, parasweep=parsweep_trait, GeneticMixing=T, Allee = T, HLatEQ = F,
      savedata = T)
