## Supplementary material for "Multi-scale Effects of Habitat Loss and the Role of Trait Variation": R code for generating data and figures: SI_Figure_S1_S2.R

```
# Code for figures with different shapes of HL curves


# # Generate data for default values when a0=0.1 [~1-2 minutes]
# ### Might already have this from previous figure, if not comment out and run the section ###
# rm(list = ls())
# source("Bagpack.R")
# foldername = "Data/e-var-mean_0p2-sd_0p08-a0_0p1"
# trait = c("e")
# parsweep_trait = c()
# obj <- MCfun_multcases(a0 = 0.1, b0 = 30, b1 = 500, bins=21,
#                        p1 = 0.7, beta = 0.8, vartrait = trait, ToffMinMaxPos = c(0,1,TRUE),
#                        ttmin=0, ttmax=0.4, ttmean=0.2, ttsd=0.08,
#                        e = 0.2, ec = 0.9, C01 = 0.5, C02 = 0.5, C11 = 50, C12 =50,
#                        N01 = 3, N02 = 4, R01 = 10, R02 = 10, foldername = foldername,
#                        mu = 0.3, d = 0.5, k = 0.003, t0 = 3500, theta = 2, times = seq(0, 8000, 1),
#                        mutations = T, parasweep=parsweep_trait, GeneticMixing=T, Allee = T, HLatEQ = F, savedata = T)


# Generate data for default values and parameter sweeping t0 [~1-2 minutes]
rm(list = ls())
source("Bagpack.R")

foldername = "Data/e-var-mean_0p2-sd_0p08-parsweep_t0"
trait = c("e")
parsweep_trait = c("t0")
obj <- MCfun_multcases(a0 = 0.1, b0 = 30, b1 = 500, bins=21,
                       p1 = 0.7, beta = 0.8, vartrait = trait, ToffMinMaxPos = c(0,1,TRUE),
                       ttmin=0, ttmax=0.4, ttmean=0.2, ttsd=0.08,
                       e = 0.2, ec = 0.9, C01 = 0.5, C02 = 0.5, C11 = 50, C12 =50,
                       N01 = 3, N02 = 4, R01 = 10, R02 = 10, foldername = foldername,
                       mu = 0.3, d = 0.5, k = 0.003, t0 = c(0,3500,6000), theta = 2, times = seq(0, 8000, 1),
                       mutations = T, parasweep=parsweep_trait, GeneticMixing=T, Allee = T, HLatEQ = F, savedata = T)


# Generate data for default values and parameter sweeping k [~1-2 minutes]
rm(list = ls())
source("Bagpack.R")
foldername = "Data/e-var-mean_0p2-sd_0p08-parsweep_k"
trait = c("e")
parsweep_trait = c("k")
obj <- MCfun_multcases(a0 = 0.1, b0 = 30, b1 = 500, bins=21,
                       p1 = 0.7, beta = 0.8, vartrait = trait, ToffMinMaxPos = c(0,1,TRUE),
                       ttmin=0, ttmax=0.4, ttmean=0.2, ttsd=0.08,
                       e = 0.2, ec = 0.9, C01 = 0.5, C02 = 0.5, C11 = 50, C12 =50,
                       N01 = 3, N02 = 4, R01 = 10, R02 = 10, foldername = foldername,
                       mu = 0.3, d = 0.5, k = c(0.001,0.003,0.061), t0 = 3500, theta = 2, times = seq(0, 8000, 1),
                       mutations = T, parasweep=parsweep_trait, GeneticMixing=T, Allee = T, HLatEQ = F, savedata = T)


#Plot the figure by sourcing the file which plots it
source("plotcode/SI_Figure_S1_S2_plotfile.R")
```
