## Supplementary material for "Multi-scale Effects of Habitat Loss and the Role of Trait Variation": R code for generating data and figures: SI_Figure_S4_S7.R

```
# Code to generate data for parameter space plots

# Parameter scan over a0 while epsilon is evolving [~35-40 mins]
rm(list = ls())
source("Bagpack.R")

foldername = "Data/e-var-mean_0p2-sd_0p08-parsweep_a0"
trait = c("e")
parsweep_trait = c("a0")
obj <- MCfun_multcases(a0 = seq(0.05,0.15,0.005), b0 = 30, b1 = 500, bins=21,
                       p1 = 0.7, beta = 0.8, vartrait = trait, ToffMinMaxPos = c(0,1,TRUE),
                       ttmin=0, ttmax=0.4, ttmean=0.2, ttsd=0.08,
                       e = 0.2, ec = 0.9, C01 = 0.5, C02 = 0.5, C11 = 50, C12 =50,
                       N01 = 3, N02 = 4, R01 = 10, R02 = 10, foldername = foldername,
                       mu = 0.3, d = 0.5, k = 0.003, t0 = 3500, theta = 2, times = seq(0, 8000, 0.5),
                       mutations = T, parasweep=parsweep_trait, GeneticMixing=T, Allee = T, HLatEQ = F, savedata = T)


# Parameter scan over b0 while epsilon is evolving [~35-40 mins]
rm(list = ls())
source("Bagpack.R")

foldername = "Data/e-var-mean_0p2-sd_0p08-parsweep_b0"
trait = c("e")
parsweep_trait = c("b0")
obj <- MCfun_multcases(a0 = 0.1, b0 = seq(15,45,1.5), b1 = 500, bins=21,
                       p1 = 0.7, beta = 0.8, vartrait = trait, ToffMinMaxPos = c(0,1,TRUE),
                       ttmin=0, ttmax=0.4, ttmean=0.2, ttsd=0.08,
                       e = 0.2, ec = 0.9, C01 = 0.5, C02 = 0.5, C11 = 50, C12 =50,
                       N01 = 3, N02 = 4, R01 = 10, R02 = 10, foldername = foldername,
                       mu = 0.3, d = 0.5, k = 0.003, t0 = 3500, theta = 2, times = seq(0, 8000, 0.5),
                       mutations = T, parasweep=parsweep_trait, GeneticMixing=T, Allee = T, HLatEQ = F, savedata = T)

# Parameter scan over b1 while epsilon is evolving
rm(list = ls())
source("Bagpack.R")

foldername = "Data/e-var-mean_0p2-sd_0p08-parsweep_b1"
trait = c("e")
parsweep_trait = c("b1")
obj <- MCfun_multcases(a0 = 0.1, b0 = 30, b1 = seq(110, 910, 40), bins=21,
                       p1 = 0.7, beta = 0.8, vartrait = trait, ToffMinMaxPos = c(0,1,TRUE),
                       ttmin=0, ttmax=0.4, ttmean=0.2, ttsd=0.08,
                       e = 0.2, ec = 0.9, C01 = 0.5, C02 = 0.5, C11 = 50, C12 =50,
                       N01 = 3, N02 = 4, R01 = 10, R02 = 10, foldername = foldername,
                       mu = 0.3, d = 0.5, k = 0.003, t0 = 3500, theta = 2, times = seq(0, 8000, 1),
                       mutations = T, parasweep=parsweep_trait, GeneticMixing=T, Allee = T, HLatEQ = F, savedata = T)


# Parameter scan over p while epsilon is evolving [~17-20 mins]
rm(list = ls())
source("Bagpack.R")

foldername = "Data/e-var-mean_0p2-sd_0p08-parsweep_p"
trait = c("e")
parsweep_trait = c("p1")
obj <- MCfun_multcases(a0 = 0.1, b0 = 30, b1 = 500, bins=21,
                       p1 = seq(0,1,0.05), beta = 0.8, vartrait = trait, ToffMinMaxPos = c(0,1,TRUE),
                       ttmin=0, ttmax=0.4, ttmean=0.2, ttsd=0.08,
                       e = 0.2, ec = 0.9, C01 = 0.5, C02 = 0.5, C11 = 50, C12 =50,
                       N01 = 3, N02 = 4, R01 = 10, R02 = 10, foldername = foldername,
                       mu = 0.3, d = 0.5, k = 0.003, t0 = 3500, theta = 2, times = seq(0, 8000, 1),
                       mutations = T, parasweep=parsweep_trait, GeneticMixing=T, Allee = T, HLatEQ = F, savedata = T)


# Parameter scan over beta while epsilon is evolving [~17-20 mins]
rm(list = ls())
source("Bagpack.R")

foldername = "Data/e-var-mean_0p2-sd_0p08-parsweep_beta"
trait = c("e")
parsweep_trait = c("beta")
obj <- MCfun_multcases(a0 = 0.1, b0 = 30, b1 = 500, bins=21,
                       p1 = 0.7, beta = seq(0,1,0.05), vartrait = trait, ToffMinMaxPos = c(0,1,TRUE),
                       ttmin=0, ttmax=0.4, ttmean=0.2, ttsd=0.08,
                       e = 0.2, ec = 0.9, C01 = 0.5, C02 = 0.5, C11 = 50, C12 =50,
                       N01 = 3, N02 = 4, R01 = 10, R02 = 10, foldername = foldername,
                       mu = 0.3, d = 0.5, k = 0.003, t0 = 3500, theta = 2, times = seq(0, 8000, 1),
                       mutations = T, parasweep=parsweep_trait, GeneticMixing=T, Allee = T, HLatEQ = F, savedata = T)


# Parameter scan over epsilon while epsilon is evolving [~30-35 mins]
rm(list = ls())
source("Bagpack.R")

foldername = "Data/e-var-mean_0p2-sd_0p08-parsweep_e"
trait = c("e")
parsweep_trait = c("ttmean")
obj <- MCfun_multcases(a0 = 0.1, b0 = 30, b1 = 500, bins=21,
                       p1 = 0.7, beta = 0.8, vartrait = trait, ToffMinMaxPos = c(0,1,TRUE),
                       ttmin=0, ttmax=0.4, ttmean=seq(0,0.4,0.02), ttsd=0.08,
                       e = 0.2, ec = 0.9, C01 = 0.5, C02 = 0.5, C11 = 50, C12 =50,
                       N01 = 3, N02 = 4, R01 = 10, R02 = 10, foldername = foldername,
                       mu = 0.3, d = 0.5, k = 0.003, t0 = 3500, theta = 2, times = seq(0, 8000, 1),
                       mutations = T, parasweep=parsweep_trait, GeneticMixing=T, Allee = T, HLatEQ = F, savedata = T)


# Parameter scan over epsilon_c while epsilon is evolving [~17-20 mins]
rm(list = ls())
source("Bagpack.R")

foldername = "Data/e-var-mean_0p2-sd_0p08-parsweep_ec"
trait = c("e")
parsweep_trait = c("ec")
obj <- MCfun_multcases(a0 = 0.1, b0 = 30, b1 = 500, bins=21,
                       p1 = 0.7, beta = 0.8, vartrait = trait, ToffMinMaxPos = c(0,1,TRUE),
                       ttmin=0, ttmax=0.4, ttmean=0.2, ttsd=0.08,
                       e = 0.2, ec = seq(0,1,0.05), C01 = 0.5, C02 = 0.5, C11 = 50, C12 =50,
                       N01 = 3, N02 = 4, R01 = 10, R02 = 10, foldername = foldername,
                       mu = 0.3, d = 0.5, k = 0.003, t0 = 3500, theta = 2, times = seq(0, 8000, 1),
                       mutations = T, parasweep=parsweep_trait, GeneticMixing=T, Allee = T, HLatEQ = F, savedata = T)


# Parameter scan over d while epsilon is evolving [~19-20 mins]
rm(list = ls())
source("Bagpack.R")

foldername = "Data/e-var-mean_0p2-sd_0p08-parsweep_d"
trait = c("e")
parsweep_trait = c("d")
obj <- MCfun_multcases(a0 = 0.1, b0 = 30, b1 = 500, bins=21,
                       p1 = 0.7, beta = 0.8, vartrait = trait, ToffMinMaxPos = c(0,1,TRUE),
                       ttmin=0, ttmax=0.4, ttmean=0.2, ttsd=0.08,
                       e = 0.2, ec = 0.9, C01 = 0.5, C02 = 0.5, C11 = 50, C12 =50,
                       N01 = 3, N02 = 4, R01 = 10, R02 = 10, foldername = foldername,
                       mu = 0.3, d = seq(0,0.5,0.025), k = 0.003, t0 = 3500, theta = 2, times = seq(0, 8000, 1),
                       mutations = T, parasweep=parsweep_trait, GeneticMixing=T, Allee = T, HLatEQ = F, savedata = T)


# Parameter scan over theta while epsilon is evolving [~38-40 mins]
rm(list = ls())
source("Bagpack.R")

foldername = "Data/e-var-mean_0p2-sd_0p08-parsweep_theta"
trait = c("e")
parsweep_trait = c("theta")
obj <- MCfun_multcases(a0 = 0.1, b0 = 30, b1 = 500, bins=21,
                       p1 = 0.7, beta = 0.8, vartrait = trait, ToffMinMaxPos = c(0,1,TRUE),
                       ttmin=0, ttmax=0.4, ttmean=0.2, ttsd=0.08,
                       e = 0.2, ec = 0.9, C01 = 0.5, C02 = 0.5, C11 = 50, C12 =50,
                       N01 = 3, N02 = 4, R01 = 10, R02 = 10, foldername = foldername,
                       mu = 0.3, d = 0.5, k = 0.003, t0 = 3500, theta = seq(0,4,0.2), times = seq(0, 8000, 0.5),
                       mutations = T, parasweep=parsweep_trait, GeneticMixing=T, Allee = T, HLatEQ = F, savedata = T)


# Plot the figure by sourcing the file.
# Parsweep plots (patch-wise equilibrium density vs parameter values) for no variation cases
source("plotcode/SI_Figure_S4_plotfile.R")


# Parsweep plots (patch-wise equilibrium density vs parameter values) for variation in epsilon
source("plotcode/SI_Figure_S7_plotfile.R")
```
