## Supplementary material for "Multi-scale Effects of Habitat Loss and the Role of Trait Variation": R code for generating data and figures: SI_Figure_S8.R

```
# Generating data and figure for comparing the variance scenarios when b1 evolves


# Generate data for default values when b1 evolves [~2-3 minutes]
rm(list = ls())
source("Bagpack.R")
foldername = "Data/b1-var-mean_400-sd_150-ttmin_100-ttmax_700"
trait = c("b1")
parsweep_trait = c()
obj <- MCfun_multcases(a0 = 0.1, b0 = 30, b1 = 500, bins=21,
                       p1 = 0.7, beta = 0.8, vartrait = trait, ToffMinMaxPos = c(0,1,TRUE),
                       ttmin=100, ttmax=700, ttmean=400, ttsd=150,
                       e = 0.2, ec = 0.9, C01 = 0.5, C02 = 0.5, C11 = 50, C12 =50,
                       N01 = 3, N02 = 4, R01 = 10, R02 = 10, foldername = foldername,
                       mu = 0.3, d = 0.5, k = 0.003, t0 = 3500, theta = 2, times = seq(0, 8000, 1),
                       mutations = T, parasweep=parsweep_trait, GeneticMixing=T, Allee = T, HLatEQ = F, savedata = T)


#Plot the figure by sourcing the file which plots it
source("plotcode/SI_Figure_S8_plotfile.R")
```
